# Definition and Signatures of Lung Fibroblast Populations in Development and Fibrosis in Mice and Men

**DOI:** 10.1101/2020.07.15.203141

**Authors:** Xue Liu, Simon C. Rowan, Jiurong Liang, Changfu Yao, Guanling Huang, Nan Deng, Ting Xie, Di Wu, Yizhou Wang, Ankita Burman, Tanyalak Parimon, Zea Borok, Peter Chen, William C. Parks, Cory M. Hogaboam, S. Samuel Weigt, John Belperio, Barry R. Stripp, Paul W. Noble, Dianhua Jiang

**Affiliations:** Department of Medicine and Women’s Guild Lung Institute, Cedars-Sinai Medical Center, Los Angeles, CA 90048, USA; Genomics Core, Cedars-Sinai Medical Center, Los Angeles, CA 90048, USA; Division of Pulmonary and Critical Care Medicine, Keck School of Medicine of University of Southern California, Los Angeles, CA 90033, USA; Department of Medicine, David Geffen School of Medicine, University of California Los Angeles, Los Angeles, CA 90095, USA; Department of Biomedical Sciences, Cedars-Sinai Medical Center, Los Angeles, CA 90048, USA

**Keywords:** lung fibroblasts, scRNA-seq, lipofibroblasts, myofibroblasts, *Ebf1*^+^ fibroblasts, Smooth muscle cells (SMCs), pericytes

## Abstract

The heterogeneity of fibroblasts in the murine and human lung during homeostasis and disease is increasingly recognized. It remains unclear if the different phenotypes identified to date are characteristic of unique subpopulations with unique progenitors or whether they arise by a process of differentiation from common precursors. Our understanding of this ubiquitous cell type is limited by an absence of well validated, specific, markers with which to identify each cell type and a clear consensus on the distinct populations present in the lung. Here we describe single cell RNA sequencing (scRNA-seq) analysis on mesenchymal cells from the murine lung throughout embryonic (E) development (E9.5 – 17.5), at post-natal day (P1 – 15), as well as in the adult and the aged murine lungs before and after bleomycin-induced fibrosis. We carried out complementary scRNA-seq on human lung tissue from a P1 lung, a month 21 lung and lung tissue from healthy donors and patients with idiopathic pulmonary fibrosis (IPF). The murine and human data were supplemented with publicly available scRNA-seq datasets. We consistently identified lipofibroblasts, myofibroblasts, pericytes, mesothelial cells and smooth muscle cells. In addition, we identified a novel population delineated by *Ebf1* (early B-cell factor 1) expression and an intermediate subtype. Comparative analysis with human mesenchymal cells revealed homologous mesenchymal subpopulations with remarkably conserved transcriptomic signatures. Comparative analysis of changes in gene expression in the fibroblast subpopulations from age matched non-fibrotic and fibrotic lungs in the mouse and human demonstrates that many of these subsets contribute to matrix gene expression in fibrotic conditions. Subtype selective transcription factors were identified and putative divergence of the clusters during development were delineated. Prospective isolation of these fibroblast subpopulations, localization of signature gene markers, and lineage-tracing each cluster are under way in the laboratory. This analysis will enhance our understanding of fibroblast heterogeneity in homeostasis and fibrotic disease conditions.

## Introduction

The adult pulmonary mesenchyme includes multiple distinct cell lineages and is centrally involved in the pathogenesis and progression of debilitating respiratory conditions like idiopathic pulmonary fibrosis (IPF)^1^. Pulmonary mesenchymal cells, including specific and unspecific fibroblast subtypes; like myofibroblasts, lipofibroblasts or parenchymal fibroblasts respectively, smooth muscle cells (SMCs) of both airway and vascular, pericytes, and mesothelial cells, undergo dynamic structural, biochemical, and functional changes during organ development and disease. This is particularly true in IPF where dysfunctional tissue repair response by mesenchymal cells is believed to be a critical factor^1^.

Recent studies utilizing single cell omics technologies, including scRNA-seq, have focused on defining the transcriptome of different cell types including lung mesenchymal cells^2–6^. The use of different databases, cells of divergent developmental or disease stages has been confounding, resulting in a range of different transcriptomic signatures being attributed to the same cell population^2,3,5,7–9^. An array of cell “specific” markers has been reported. However highly discriminative markers, especially for fibroblast subpopulations, remain elusive and the majority of markers are non-specific and expressed by multiple cell types.

Major studies, many unpublished and released as preprints, of lung cell types in IPF, interstitial lung disease, chronic obstructive pulmonary disease, sarcoidosis, non-specific interstitial pneumonia and chronic hypersensitivity pneumonitis, have identified anywhere between six and 11 distinct mesenchymal subpopulations^4,6,10–13^. While lipofibroblasts, myofibroblasts, SMCs, pericytes and mesothelial cells are commonly reported in these studies, but the transcriptomic signatures differ. Frequently publications identify subtypes by a mixture of location and/or discriminative gene expression^4,6,10–13^. To add the confusion, clusters identified by high expression of a delineating gene, for example *WIF1*, in one publication are subsequently identified by others as a predefined mesenchymal cell type, like myofibroblasts in which *WIF1* is reported as a delineating marker gene^4,12^. The current approach likely leads to overlap of distinct clusters and does little to resolve the controversies regarding the definitive transcriptomic signature of the pulmonary mesenchymal populations.

Of the known mesenchymal subpopulations, myofibroblasts have been of singular interest given their role as the prominent producers of extracellular matrix (ECM), ability to restore tissue integrity after injury and as the postulated effector cell of pulmonary fibrosis^14^. However, commonly reported marker genes, such as *Acta2,* are expressed prominently by SMCs or other mesenchymal cells making fractionation and further study difficult^15–17^. Similarly the description of lipofibroblasts, a fibroblast subtype prominent in the developing rodent lung characterized by lipid inclusions, and even their presence in the human lung remains controversial^18–21^. The lack of specificity of reported markers for other known mesenchymal lung components is true also for SMCs, pericytes and mesothelial cells. Further, whether these mesenchymal subpopulations transition or acquire a myofibroblast like phenotype is yet to be determined. To date, no consistently reported signature for the different mesenchymal subtypes has been reported. A better definition of specific markers and detailed genetic lineage of these mesenchymal cell subtypes is needed to better understand lung development and disease.

Therefore, in this study we undertook a longitudinal scRNA-seq analysis of mesenchymal cells from murine and human lungs at different developmental stages, adult and aged murine lungs following bleomycin induced fibrosis, and lung tissue from healthy donors and patients with IPF. We accessed and re-analyzed scRNA-seq data from relevant published studies. We identified the genetic programs of murine and human lung mesenchymal cells from embryonic development to adulthood and in disease. We examined mesenchymal subtypes for known marker gene expression and specification and identified novel markers genes that were more specific for each mesenchymal subtype than those in the literature. Comparative analysis of the genetic profiles of the mesenchymal cell subtypes in the non-fibrotic and fibrotic lung was performed. These data provide a longitudinal and comprehensive genetic definition of murine and human lung mesenchymal subtypes and suggest all mesenchymal subtypes may contribute to pulmonary fibrosis. These findings contribute to the study and understanding of lung development and may aid the development of targeted therapies for the treatment of pulmonary fibrosis.

## Results

### scRNA-seq on E17.5 murine lung identified six fibroblast subpopulations

Myofibroblasts are essential components of the growing secondary septa, and most prominent during alveolarization in late embryogenesis; when lipofibroblasts also emerge, and postnatal stages of lung development in mice^18^. The transcriptome and specific markers for neither myofibroblasts nor lipofibroblasts have been clearly described. Therefore, in order to identify a clear transcriptomic profile for lipofibroblasts and myofibroblasts we first performed scRNA-seq on E17.5 embryonic murine lungs. We collected live cells from three E17.5 murine lungs (Fig. S1A). Sequencing libraries were prepared from FACS sorted cells using the 10x Genomics Chromium system. After quality control (QC) (Fig. S1B, C), samples were integrated (Fig. 1A). The cells were visualized in two dimensions according to their gene expression profiles using Uniform Manifold Approximation and Projection (UMAP). Mesenchymal, immune, epithelial and endothelial cell clusters were well separated by distinct gene expression profiles (Fig. 1B, S1D-F). The mesenchymal fraction was subset from epithelial, immune and endothelial cells and the fraction purity was confirmed (Fig. S2A-C, Supplementary Table 1). Lipofibroblasts (*Plin2* and *Tcf21*^+^), myofibroblasts (*Acta2* and *Pdgfra*^+^), proliferative fibroblasts (*Hmmr* and *Mki67*^+^) and mesothelial cells (*Wt1* and Upk3b^+^) were identified by discrete expression of widely reported marker genes (Fig. 1C)^2,7,8,17,22^. Two major populations could not be identified using any known markers in the literature. One subpopulation was identified as *Ebf1*^+^ fibroblasts due to the condensed and specific expression of this transcription factor. The remaining cluster, after all others had been identified, were named intermediate fibroblasts due to the low expression of genes from multiple other populations (Fig. 1C).

**Figure 1.**
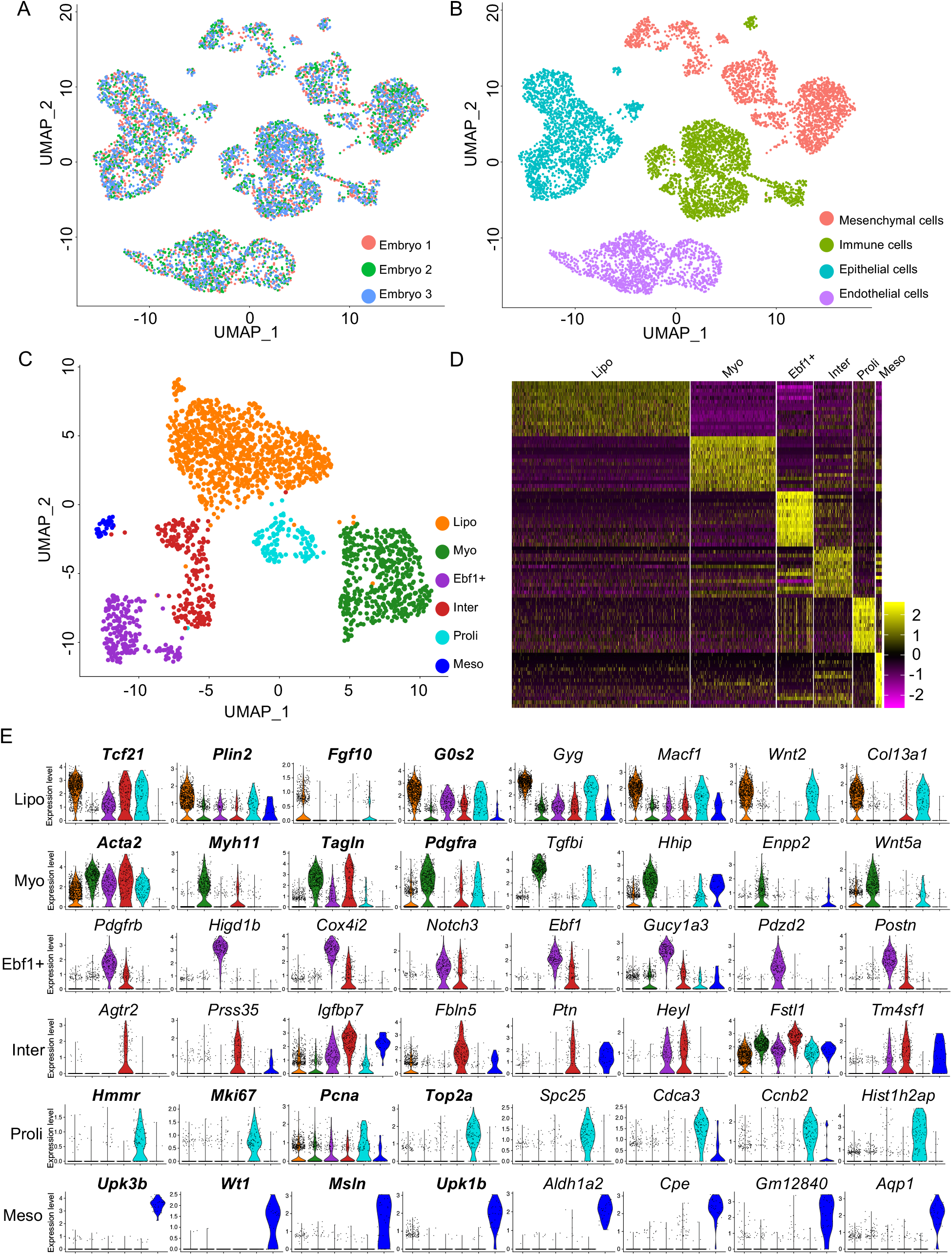
scRNA-sequencing on E17.5 lung identified fibroblast subtypes. UMAP visualization of cell origin, grouping (**A**) and cell type clustering (**B**) in E17.5 mouse lung total cell scRNA-seq data. (**C**) Six fibroblast clusters were defined. (**D**) Heat map of lung fibroblasts normalized signal show fibroblast subtypes changes by top 15 genes (rows) for individual subtype cells (columns). (**E**) Violin plot representation showing relative expression of the fibroblasts cluster signature genes. Genes in bold and normal text represent the classical and novel signature genes, respectively. Lipo, lipofibroblasts, Myo, myofibroblasts, Ebf1^+^, Ebf1^+^ fibroblasts, Inter, intermediate fibroblasts, Proli, proliferative fibroblasts, Meso, Mesothelial cells.

To validate the subpopulation identification, linear dimensional reduction was performed using two further assays. Firstly, the cells were visualized using t-Distributed Stochastic Neighbor Embedding (t-SNE) with the proliferative fibroblasts and mesothelial cells groups excluded (Fig. S2D). Secondly, principal component analysis (PCA) was visualized using Independent Component Analysis on module data (k-means) and metagene data. In both instances the remaining mesenchymal cells separated into four clear clusters in both two and three dimensions (Fig. S2E). *Pdgfrb* expression was present in all subpopulations and highest in the *Ebf1*^+^ population (Fig. 1E, S2F, G) visualized by violin plot, t-SNE and UMAP analysis, suggesting that *Ebf1*^+^ fibroblast cluster might include pericytes. Heatmap visualization of the top 15 differentially expressed genes of each cluster revealed a distinct pattern of gene expression in each identified cluster (Fig. 1D) and a selection of the top specific genes for each subpopulation were visualized by dot plots (Fig. S2H). The expression of eight specific genes from each subpopulation were visualized by violin plots (Fig. 1E).

To further validate the definition of the mesenchymal subpopulation, we performed another unbiased analysis – single cell ATAC-seq (scATAC-seq) on E17.5 mouse lung. After QC (Fig. S3A-B) and clustering major cell types (Fig. S3C-F), mesenchymal cells were extracted and clustered mesenchymal cells. By checking the mesenchymal cell cluster specific genes identified above, we defined similar mesenchymal cell clusters, lipofibroblast, *Ebf1*^+^ fibroblast, myofibroblast, intermediate fibroblast and mesothelial cell (Fig. S3G-I) and similar cluster-specific growth factors and transcriptional factors were confirmed (Fig. S3J).

Self-organizing maps (SOM) were used to visualize coincidental gene sets in each subpopulation of E17.5 fibroblasts (Supplementary Methods). Multiple subtype-specific gene signatures were determined, including extracellular space, collagen-containing extracellular matrix, structural constituent of ribosome, ribosome, DNA-binding transcription activator activity, RNA polymerase, nucleus, cell adhesion, plasma me, extracellular region, extracellular space, stress fiber, plasma membrane, Golgi cisterna membrane and protein glycosylation. Notably, lipofibroblasts displayed opposite gene signatures to that of myofibroblasts (Fig. S4A).

A customizable suite of single-cell R-analysis tools (SCRAT) based on SOM machine learning was used to analyze for sample similarity and perform pseudo-time analysis (Supplementary Methods). The correlation-spanning tree and trajectory report suggested a directed hierarchical relationship between the fibroblast subpopulations. The correlation-spanning tree and k−nearest neighbor graph began from the lipofibroblast cluster, bifurcated to intermediate fibroblasts and finally bifurcated to *Ebf1*^+^ fibroblasts and myofibroblasts (Fig. S4B-E). The top 15 activated and inhibited regulators among the differentially expressed genes of each subpopulation are reported in Fig. S5 by Ingenuity Pathway Analysis (IPA).

### Identification of murine lung fibroblast subpopulations throughout lung development and following bleomycin induced fibrosis

scRNA-seq datasets from earlier developmental stages (E9.5, 10.5, 11.5, 12.5, 14.5, 16.5) were examined for the conserved expression of the transcriptomic profiles of each subpopulation identified in the E17.5 dataset. The mesenchymal cells from E9.5, E10.5 and E11.5 datasets were extracted and combined^23^ (Fig. S6A). Distinct endoderm and mesoderm clusters were identified by visualizing known markers, endoderm specific transcription factors, *Nkx2-1* and *Foxa2,* and mesoderm specific transcription factors, *Tbx5* and *Osr1* (Fig. 2A, B, S6B-C). However, the fibroblast subtype-specific transcriptomic profiles identified in the E17.5 dataset were not distinct in the combined E9.5-11.5 datasets. This suggests the differentiation fate of the mesodermal cells at E9.5-E11.5 lung was not yet determined.

**Figure 2.**
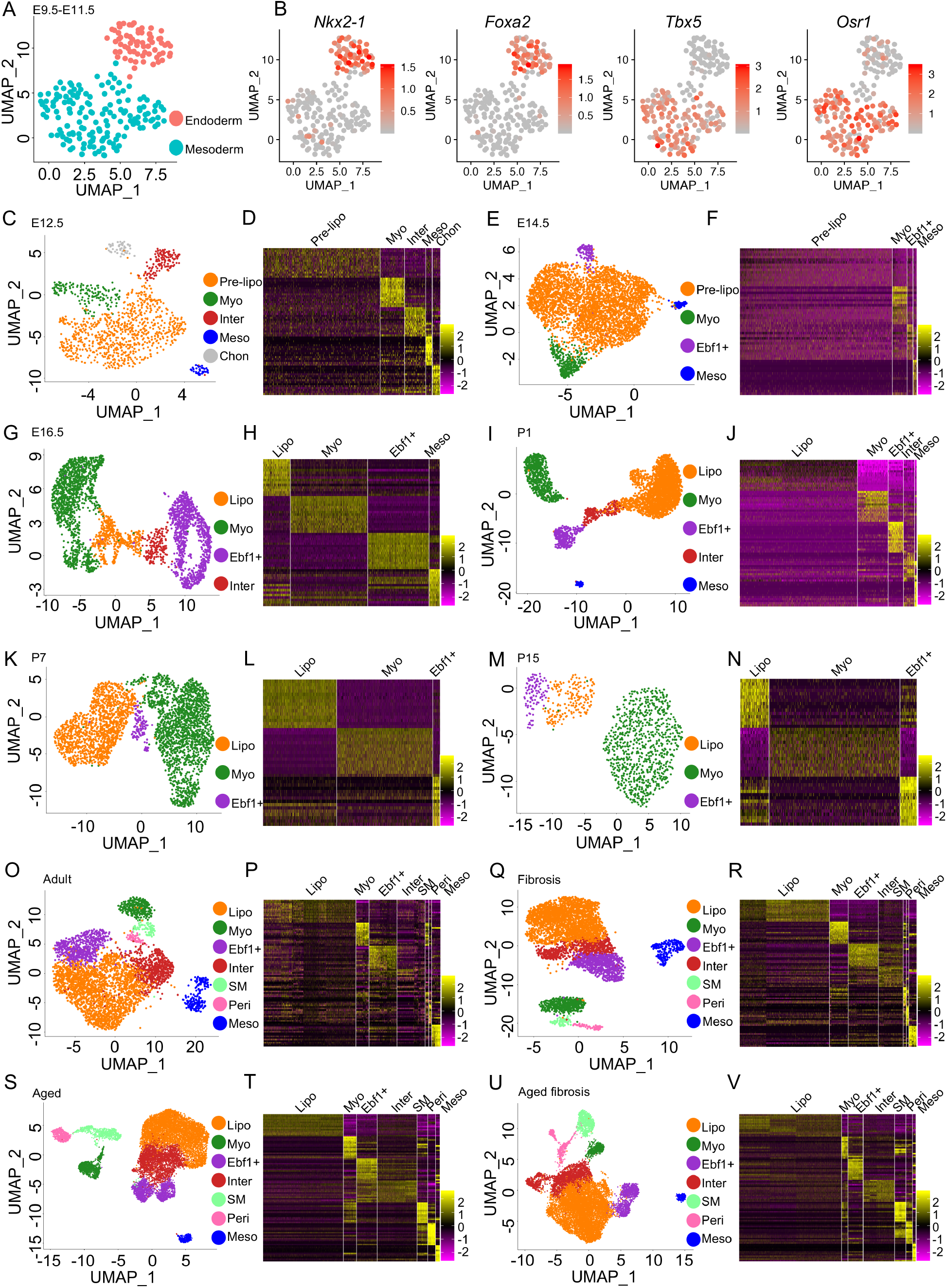
Identification of lung fibroblast subtypes in different time points. UMAP visualization of E9.5-E11.5 lung endoderm and mesoderm (**A**) and the expression of specific transcription factors (**B**). Fibroblast subtype classification and heatmaps based on tope 15 genes in E12.5 (**C**, **D**), E14.5 (**E**, **F**), E16.5 (**G**, **H**), P1 (**I**, **J**), P7 (**K**, **L**) and P15 (**M**, **N**) mouse lung. Lung fibroblast subtype classification and heatmaps based on tope 15 genes in young (**O**, **P**, **Q**, **R**) and aged (**S**, **T**, **U**, **V**) mice before (**Q**, **P**, **S**, **T**) and after (**Q**, **R**, **U**, **V**) bleomycin injury. Pre-lipo, Pre-lipofibroblasts, Lipo, lipofibroblasts, Myo, myofibroblasts, *Ebf1^+^*, *Ebf1*^+^fibroblasts, Inter, intermediate fibroblasts, Meso, Mesothelial cells, Chon, Chondrocytes.

CD45^−^ lung cells from an E12.5 murine lung dataset and all cells from an E14.5 murine lung dataset were extracted from recently published studies^24,25^. The mesenchymal cells were subset from the data following QC (Fig. S7A-E, S8A-E, Supplementary Table 1). Mesenchymal subpopulations were identified using known markers and the transcriptomic profiles identified at E17.5 (Fig. 2C-F). Each cluster had a distinct gene expression profile (Fig. 2D, 2F, S7F, S8F). αSMA-GFP; Tbx4-Cre; Rosa26-tdTomato (Tbx4-lineage^+^, αSMA^+^) fibroblasts were sorted from E16.5 murine lungs using the same FACS strategy reported in our previous study (Fig. S9A)^2^. scRNA-seq analysis was carried out on the sorted E16.5 cells (Fig. S9B-G, Supplementary Table 1). The subpopulations were identified as described previously (Fig. 2G, H). The gene profile of each cluster was homologous to the corresponding mesenchymal cluster in the E17.5 murine lung (Fig. S9H). scRNA-seq data from P1 lungs, Pdgfra-GFP^+^ mesenchymal cells from P7 and P15 murine lungs, were accessed from published studies^3,17^. QC and data integration, if necessary, were performed as described previously (Fig. S10A). The major cell types were identified, and the mesenchymal cells extracted from the P1 mouse lung (Fig. S10B-E, Supplementary Table 1). After removing the endothelial and immune cells in the P15 mouse lung dataset (Fig. S12A-D) and confirming the purity of the subset mesenchymal population (Fig. S10E, 11B, 12E), the mesenchymal cells were clustered, and the subpopulations identified (Fig. 2I, K and M) and each cluster had a distinct gene expression profile (Fig. 2J, L, N, S10F, S11C and S12F).

scRNA-seq data from adult normal murine lungs were accessed from our and others’ previously published studies (Fig. S13A-H, Supplementary Table 1)^5,15,26–28^. The mesenchymal cells were extracted, integrated, clustered and the purity of the mesenchymal faction was confirmed (Fig. S14A-C). The transcriptomic profile of the identified subpopulations was homologous to that observed in earlier datasets (Fig. 2O-P, S14D). Adult murine lung mesenchymal cells, 21 days after bleomycin induced injury, from two of our previously reported studies were extracted, integrated and clustered as described above^15,28^ (Fig. S15A-D). After clustering, corresponding mesenchymal subpopulations to the non-fibrotic controls were identified (Fig. 2Q-R, S15E).

As IPF is a disease of aging, to further investigate pulmonary fibroblast lineage and the fibroblast subtypes present during development and disease, mesenchymal cells (EPCAM/CD31/CD45^−^) were collected from three control aged mouse lungs and aged matched lungs 14 days after bleomycin injury. QC, integration, mesenchymal fraction extraction and purity confirmation were performed as described previously (Fig. S16A-I, S17A-B, S18A-I and S19A-B). The mesenchymal cell subpopulations identified in the aged normal and fibrotic lungs were similar to those identified in adult mouse lungs (Fig. 2S and U) and each cluster showed a distinct gene expression profile (Fig. 2T and V, S17C and S19C).

### Lipofibroblast specific and timepoint specific signature genes

We identified a distinct lipofibroblast cluster in murine lungs at each developmental and disease stage. We determined the differentially expressed genes of lipofibroblasts at each stage and visualized the top genes from representative developmental stages using volcano plots (Fig. S20A). An extended selection of time point-specific lipofibroblast genes were visualized by dot plots (Fig. S2F, S7F, S8F, S9H, S10F, S11C, S12F, S14D, S15E, S17C, S19C). Several genes are commonly used to identify lipofibroblasts, including *Tcf21, Plin2, Fgf10* and *G0s2* (G0/G1 Switch 2)^7,8,29^. To address the specification and expression of these genes, we examined the transcript level of these four representative genes and top four time point-specific genes using violin plots (Fig. 3A-B). We found either high background or very low transcript of these four known genes in some time points. Among the top differentially expressed genes in murine lung lipofibroblasts; *Limch1* (LIM and calponin homolog domains 1), *Gyg* (glycogenin), *Macf1* (Microtubule actin cross-linking factor 1), *Mfap4* (microfibril associated protein 4), *Npnt* (nephronectin), *Wnt2*, *Col13a1* (collagen type XIII alpha 1 chain), and *Inmt* (indolethylamine N-methyltransferase) were consistently expressed, and discriminative in the lipofibroblast clusters at all time points (Fig 3B). *Gyg* encodes a member of the glycogenin family and showed very specific expression patterns in pre-lipofibroblasts and lipofibroblasts (Fig. 3C, S20B). Among all the novel lipofibroblast specific genes identified, *Gyg*, *Macf1*, *Wnt2* and *Co13a1* were the most specific and consistently expressed compared to canonical markers (Fig. 3C).

**Figure 3.**
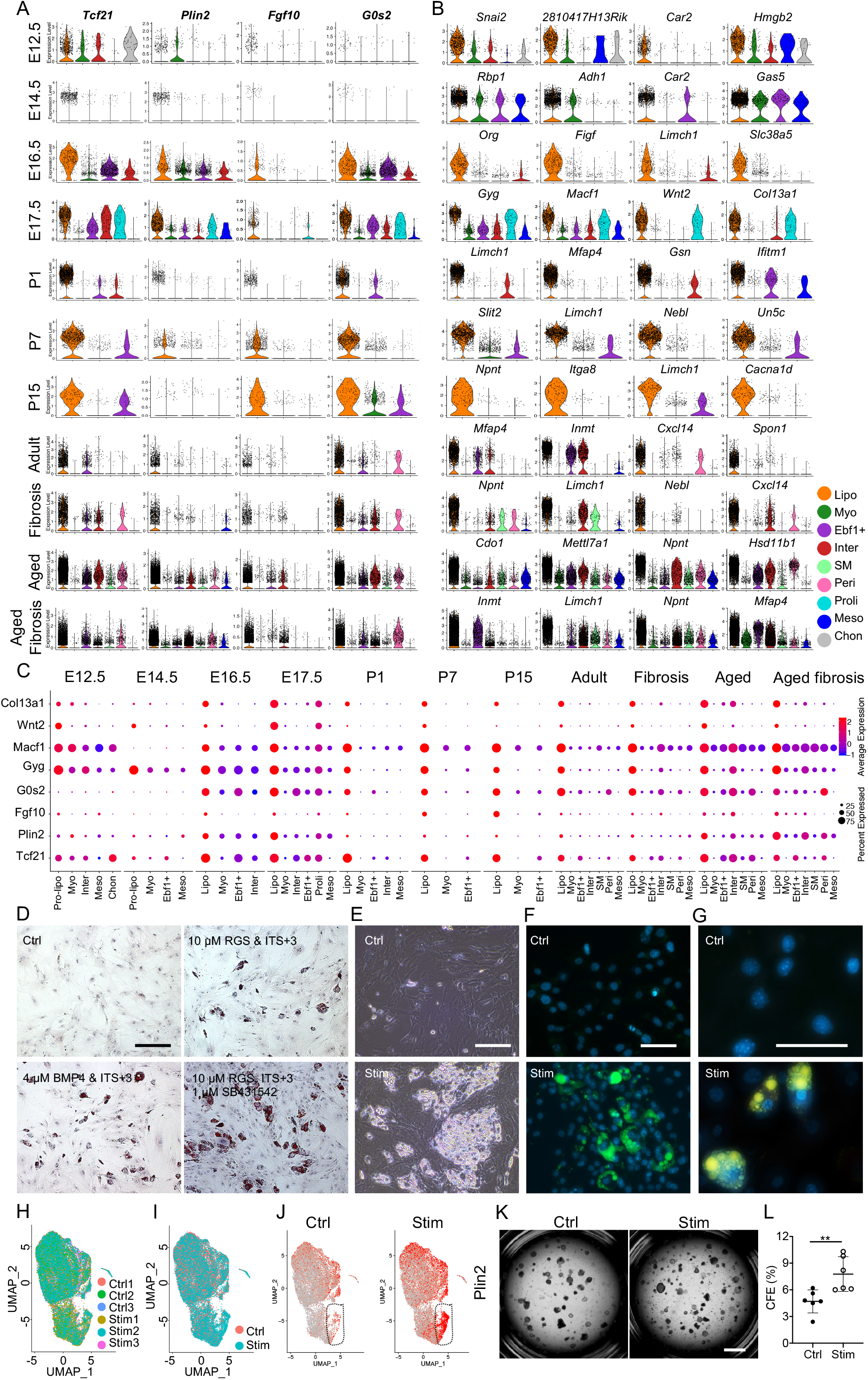
Identification and induction of lung lipofibroblast specific markers. Visualization of 4 known lipofibroblast markers (**A**) and top 4 timepoint specific genes (**B**) by violin plots. (**C**) Comparison of known and novel lipofibroblast markers in each developmental timepoints, normal and fibrosis lungs. (**D**) Representative Oil Red O staining and of murine lung fibroblasts and a lipofibroblast like phenotype was induced by stimulation by different conditions. (**E**) Morphologies of the stimulated lipofibroblast like cells and control cells were visualized by representative phase contrast images. Immunofluorescent images of the neutral lipid stained by BODPIY 493/503 (**F**) or the lipid stained Nile Red (**G**) in control and stimulated cells to confirm the presence of lipid droplets in the stimulated lipofibroblast like cells. Sample integration and cell distribution (**H**, **I**) of the control and stimulated cells by scRNA-seq and transcript of lipofibroblast common marker, Plin2 (**J**), were visualized by UMAP. (**K** - **L**) Colony formation assays were performed to examine the supporting potentials of the control and stimulated fibroblasts. Pre-lipo, Pre-lipofibroblasts, Lipo, lipofibroblasts, Myo, myofibroblasts, Ebf1+, Ebf1+ fibroblasts, Inter, intermediate fibroblasts, SMC, smooth muscle cells, Peri, Pericytes, proli, proliferative fibroblasts, Meso, Mesothelial cells, Chon, Chondrocytes. Scale bar, 20 μm (**D**, **E**, **F**, **G**) and 1 mm (**K**).

Lipofibroblasts were sorted by FACS from adult murine lungs using the top differentially expressed genes that encoded cell surface proteins in the scRNA-seq dataset, represented by Cd249 (aka *Enpep*, Glutamyl aminopeptidase) (Fig. S21A-C). The top differentially expressed genes in bulk-sequenced Cd249^+^ fibroblasts in comparison to Cd249^−^ fibroblasts overlapped substantially with the highly discriminative lipofibroblast genes in the scRNA-seq analysis, like *Limch1, Col13a1, Fgf10*, and *Tcf21* (Fig. S21D). Cultured fibroblasts in which a lipofibroblast-like phenotype had been induced using conventional methods displayed pronounced lipid inclusions (Fig. 3D-G). scRNA-seq analysis of lipofibroblast-like cells demonstrated that stimulated lipofibroblast-like cells displayed high transcript expression of canonical makers, like *Plin2*, as reported by others (Fig. 3H-J, S21D). However, the *in vivo* lipofibroblast transcriptomic signature was not localized to the *Plin2^high^* population that emerged in the stimulated cells (Fig. S21E). Colony forming assays using lipofibroblast-like cells demonstrated that they were more supportive of AEC2 colony formation than unstimulated cells (Fig. 3K, L).

### Delineation of SMCs and myofibroblasts

The transcriptomic profiles of myofibroblasts and SMCs have not yet been definitively determined and these populations are yet be to clearly distinguished from each other at the mRNA level without localization. *Acta2, Myh11* (myosin-11), *Tagln* (transgelin) and *Pdgfra* are widely reported myofibroblast marker genes but are highly expressed in other mesenchymal subtypes^16,17,22,30–32^. In the current study, we identified clear myofibroblast clusters in embryonic and postnatal, adult and aged normal and bleomycin injured mouse lung and also identified clear SMC clusters in adult and aged normal and bleomycin injured mouse lung (Fig. 2O-V). We examined the differentially expressed genes of the SMC and myofibroblasts subpopulations and visualized them using volcano (Fig. 4A) and dot plots (Fig. S2F, S7F, S8F, S9H, S10F, S11C, S12F, S14D, S15E, S17C, S19C). *Acta2*, *Myh11* and *Tagln* were preferentially expressed in the myofibroblast cluster in E12.5, E14.5, E16.5, E17.5, P7 and P15 murine lungs with limited expression in other clusters (Fig. 4B, S22A). In the P1 mouse lung, these genes showed little transcript expression in the myofibroblast cluster (Fig. 4B). *Tgfbi* (transforming growth factor beta induced), *Hhip* (hedgehog interacting protein), *Enpp2* (ectonucleotide pyrophosphatase/phosphodiesterase 2), *Egfem1* (EGF-like and EMI domain-containing protein 1), *P2ry14* (P2Y purinoceptor 14), *Wnt5a*, *Nnat* (neuronatin), *Mustn1* (musculoskeletal, embryonic nuclear protein 1), *Actg2* (actin gamma 2, smooth muscle) and *Cnn1* (calponin 1) were among the top differentially expressed genes. These genes were found to be highly discriminative and conserved between time points in myofibroblasts (Fig. 4A-B, S22B). SMC related gene expressing clones were detected in some datasets but a separate SMC cluster in the embryonic and early postnatal lung datasets could not be detected (Fig. 2C-N, S23). These data suggest that SMCs in the embryonic and early postnatal day lung cannot be definitively distinguished from myofibroblasts at the mRNA level. In the adult and aged normal and fibrotic murine lung, distinct SMC clusters were identified (Fig. 2O-V). *Acta2*, *Myh11*, *Actg2* and *Actc1* (Actin alpha cardiac muscle 1), commonly reported SMC marker genes, were among the top DE genes from the adult and aged SMC clusters (Fig. 4C-D, S22I). In the mature adult and aged murine lungs, SMC clusters were identifiable but were not clearly separated from myofibroblast clusters (Fig. 2O-V and 4D). Myofibroblasts showed similar gene profiles with embryonic lungs and we visualized four of the most specific and highly expressed genes, *Tgfbi*, *Hhip*, *Enpp2* and *Wnt5a* in all the mouse lungs of different timepoints (Fig. 4B-D, S22C-H, J). We did not observe increased myofibroblast cell number in fibrotic lungs compared to control lungs (Fig. 2O-V and 4D). More time point-specific myofibroblast and SMC genes were visualized by dot plots (Fig. S2F, S7F, S8F, S9H, S10F, S11C, S12F, S14D, S15E, S17C, S19C).

**Figure 4.**
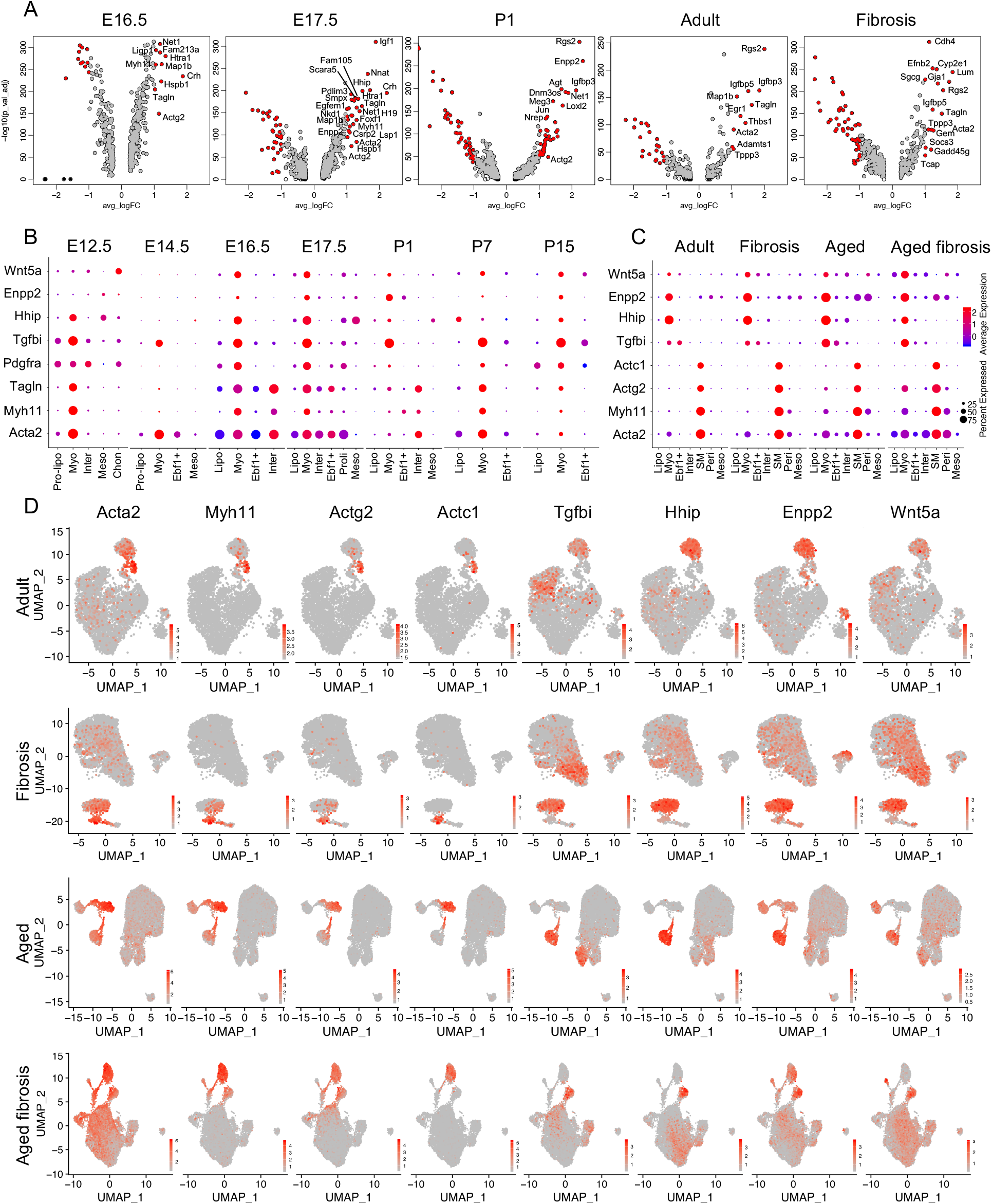
Identification of murine lung myofibroblast and SMC cluster. (**A**) Visualization of differentially expressed genes of myofibroblasts at each time point by Volcano plots. Genes in red, p-value < 10^−5^, fold-change (logFC)> 1, genes in black, p-value < 10^−5^, logFC < 1, genes in grey, p-value > 10^−5^, logFC > 1. (**B**) Comparison of known and novel myofibroblast markers in embryonic and postnatal mouse lungs. Visualization of myofibroblasts and SM cells in adult and aged normal and fibrosis mouse lung by dot plots (**C**) and UMAPs (**D**). Pre-lipo, Pre-lipofibroblasts, Lipo, lipofibroblasts, Myo, myofibroblasts, *Ebf1*^+^, *Ebf1*^+^ fibroblasts, Inter, intermediate fibroblasts, SM, smooth muscle cells, Peri, Pericytes, proli, proliferative fibroblasts, Meso, Mesothelial cells, Chon, Chondrocytes.

### Identification of an *Ebf1*^+^ fibroblast subtype and pericytes

A distinct, previously unidentified cluster of mesenchymal cells delineated by *Ebf1* expression, emerged at E14.5 and was identified at all time points examined (Fig. 2E-N). Differential expression analysis revealed time point-specific genes for the *Ebf1*^+^ cluster in each dataset (Fig. 5A). The transcriptomic profile of embryonic *Ebf1*^+^ fibroblasts represented by *Ebf1*, *Gucy1a3* (Guanylate cyclase soluble subunit alpha-3), *Pdzd2* (PDZ Domain Containing 2), *Postn* (Periostin), *Pdgfrb, Higd1b* (HIG1 Hypoxia Inducible Domain Family Member 1b), *Cox4i2* (Cytochrome C Oxidase Subunit 4i2) and *Notch3* was consistent up to P1 (Fig. 5B, S24A, C-H). From P7 onwards the transcriptomic profile was better represented by *Ebf1*, *Serpinf1* (Serpin Family F Member 1), *Postn, Col14a1* and *Pi16* (Peptidase Inhibitor 16) (Fig. 5C and S24B, I, J). In adult and aged mesenchymal cells, two *Ebf1*^+^ clusters were identified (Fig. 2O-V). One *Ebf1*^+^ cluster we identified as pericytes due to condensed expression of known pericyte markers *Cspg4* (*Ng2*) and *Pdgfrb* (Fig. 2O-V, 5D-E, S24K). The other distinct cluster was also *Ebf1*^+^ and expressed the novel transcriptomic signature identified from P7 onwards (Fig. 5D-E, S24K). Discriminative marker genes for the *Ebf1*^+^ cluster in the E14.5-P1 lungs included genes, for example *Higd1b*, *Cox4i2* and *Notch3*, subsequently identified among the top differentially expressed genes of adult/aged lung pericytes (Fig. 5E, S24K). These data suggest that pericytes and adult *Ebf1*^+^ fibroblasts diverge during early post-natal development but may share a common lineage.

**Figure 5.**
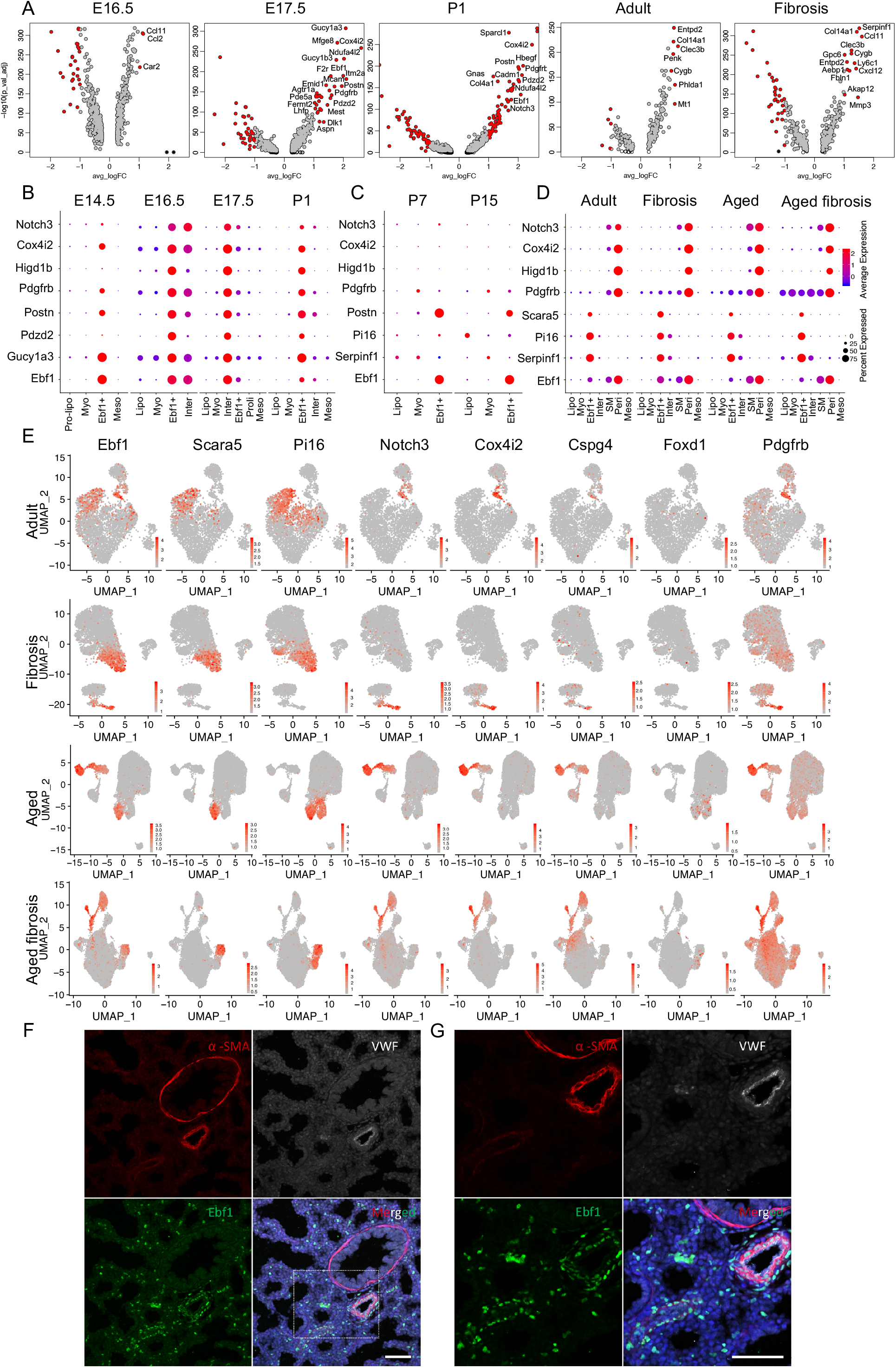
Identification of mouse lung pericytes and *Ebf1*^+^ fibroblasts. (**A**) Visualization of differentially expressed genes of *Ebf1*^+^ fibroblasts at each time point by Volcano plots. Genes in red, p-value < 10^−5^, fold-change (logFC) > 1, genes in black, p-value < 10^−5^, logFC < 1, genes in grey, p-value > 10^−5^, logFC > 1. Visualization of Ebf1+ fibroblast specific genes in embryonic (**B**) and postnatal (**C**) mouse lungs, and *Ebf1*^+^fibroblasts and pericytes in adult and aged normal and fibrosis lung (**D**) by dot plots. Top 3 specific genes of *Ebf1*^+^fibroblasts and 5 known genes of pericytes were visualized by UMAP (**E**). (**F**-**G**) aSMA, VWF and Ebf1 staining on E17.5 mouse lung section to visualize Ebf1 protein localization. Pre-lipo, Pre-lipofibroblasts, Lipo, lipofibroblasts, Myo, myofibroblasts, *Ebf1*^+^, *Ebf1*^+^ fibroblasts, Inter, intermediate fibroblasts, SM, smooth muscle cells, Peri, Pericytes, proli, proliferative fibroblasts, Meso, Mesothelial cells, Chon, Chondrocytes. Scale bar, 50 μm.

Traditional pericyte markers, like *Cspg4*, displayed low transcript expression while *Pdgfrb* was condensed in pericytes but with high background in other clusters. *Foxd1*, a common marker of pericyte lineage, was weakly expressed in *Ebf1*^+^ fibroblasts and undetectable in any pericyte cluster in the murine lung (Fig. 5E)^33^. Novel pericyte markers identified in our analysis were expressed at a greater level, with greater specification than commonly used pericyte marker genes (Fig. 5D, E, S24K). More timepoint-specific *Ebf1*^+^ fibroblast and pericyte genes were visualized by dot plots (Fig. S2F, S7F, S8F, S9H, S10F, S11C, S12F, S14D, S15E, S17C, S19C).

At protein level, co-staining for Ebf1, the endothelial cell marker, vWF (von Willebrands factor), and *α*SMA, indicated that Ebf1^+^ cells were not only perivascular but also in interstitial lung tissue (Fig. 5F-G). This further suggests that the *Ebf1*^+^ population consists of both pericytes and a distinct fibroblast subtype.

### Matrix gene expression in normal and bleomycin-injured murine lungs

As significant increased myofibroblast number was not detected in our analysis, which denied the well-known hypothesis of mesenchymal cells transition into myofibroblasts in fibrotic lung. To determine the possible mechanism, age matched non-fibrotic and fibrotic mesenchymal cells were combined for further analysis (Fig. 6A, 6G). Major ECM associated genes in fibrosis, represented by *Col1a1* and *Fn1* displayed an increased expression in the fibrotic mesenchymal cells compared to mesenchymal cells from non-fibrotic aged matched controls (Fig. 6C-F, I-L, S25). The presence of all previously identified fibroblast subtypes in the integrated datasets was confirmed (Fig. 6B, H). UMAP visualization confirmed increased expression of the known matrix related genes and the expression of novel myofibroblast markers we had identified in all the mesenchymal clusters (Fig. S26). These data suggest that bleomycin-induced fibrotic injury increases the expression of the ECM related genes in all mesenchymal cell subtypes.

**Figure 6.**
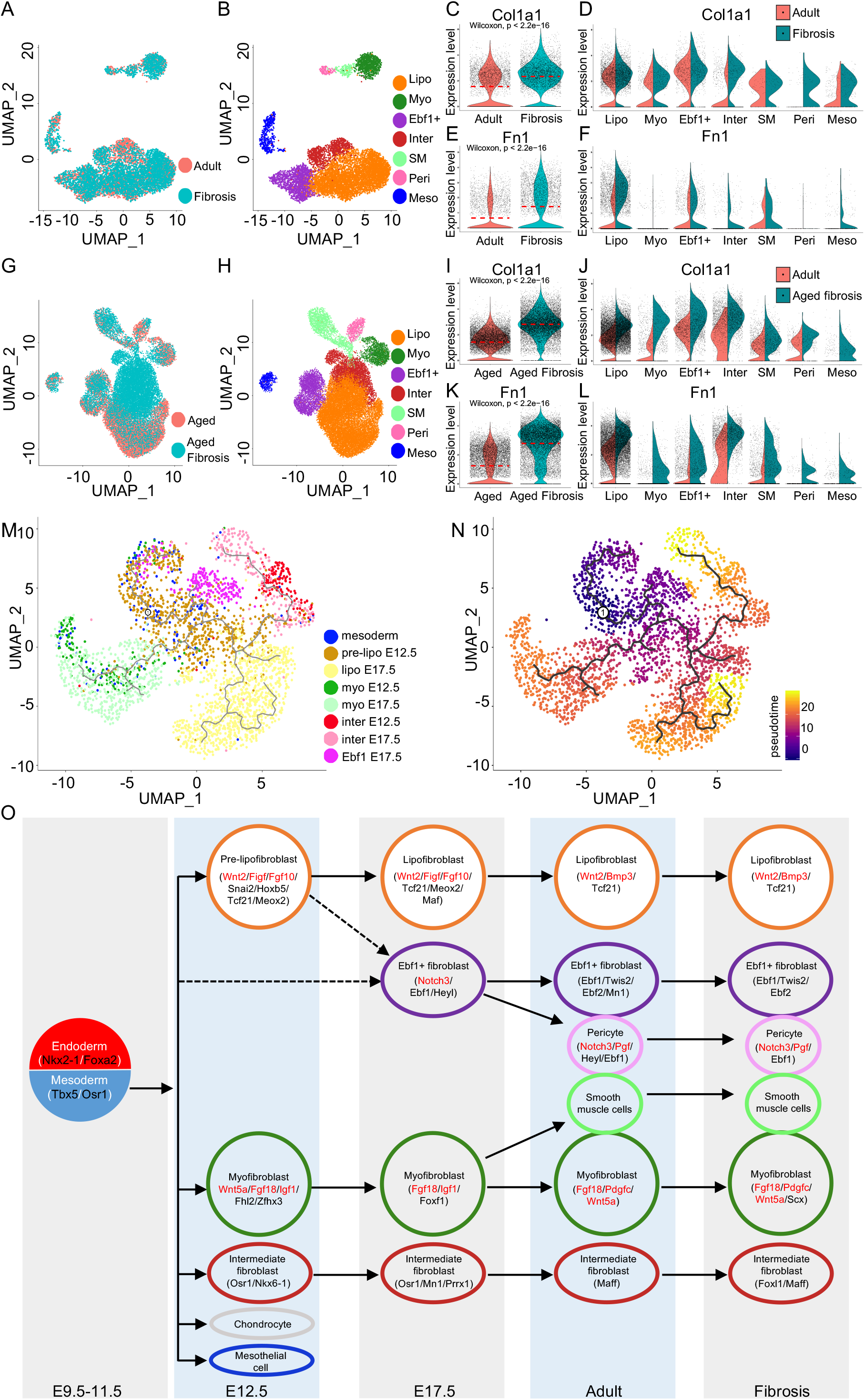
Lineage graph of mouse lung mesenchymal cell subtypes. Cell integration of adult and fibrosis mesenchymal cells (**A**), aged and aged fibrosis mesenchymal cells (**G**). Cell subtype definition of integrated adult and fibrosis (**B**), aged and aged fibrosis (**H**) mesenchymal cells. Comparison of Col1a1 (**C**, **D**, **I** and **J**) and Fn1 (**E**, **F**, **K** and **L**) expression in total mesenchymal cells (**C**, **E**, **I** and **K**) and mesenchymal cell subtypes (**D**, **F**, **J** and **L**) of adult and fibrosis lungs, aged and aged fibrosis lungs. (**M** and **N**) Lineage bifurcation and differentiation potentials of mesenchymal cell subtypes in embryonic lungs. (**O**) Lineage graph of mouse lung mesenchymal cell subtypes labelled by specific transcription factors and growth factors. Lipo, lipofibroblasts, Myo, myofibroblasts, *Ebf1*^+^, *Ebf1*^+^ fibroblasts, Inter, intermediate fibroblasts, Proli, proliferative fibroblasts, Meso, Mesothelial cells

*Col14a1* and *Col13a1*, which we previously reported as matrix fibroblast^2^ were found specific expressed in lipofibroblast or *Ebf1*^+^ clusters specific genes at different timepoints (Fig. S27A). Vim (Vimentin) is an often used as mesenchymal cell marker and in some instances reportedly a gene specific for myofibroblast^22^, however, here we found that *Vim* transcript expression was highest in endothelial cells and was detectable in mesenchymal and immune cells. It was rarely detected in epithelial cells (Fig. S27B). This suggests that *Vim* should not be used as mesenchymal cell marker. We also checked commonly used mesenchymal cell marker, *Pdgfra* and *Pdgfrb*, and found *Pdgfrb* expression overlapped with *Pdgfra* expression in some datasets while in others the expression of these two genes was well separated with high background overall (Fig. S28). In adult and fibrotic mouse lungs, *Pdgfra* was well separated from *Acta2*^+^ cells and *Pdgfrb*^+^ cells, but showed good overlap with *Tcf21* (Fig. S29A, C, D, E, G, H). *Pdgfrb* showed some overlap with one of the *Ebf1* expressing fibroblast cluster, pericytes (Fig. S29B and F).

### Differentiation potential of the embryonic mesenchymal cell clusters

To investigate the differentiation potential of the mesenchymal cell clusters at different embryonic stages, the identified mesoderm cells from the E9.5-E11.5 datasets were integrated with the E12.5 and E17.5 mesenchymal cell clusters. The integrated data was projected onto SCRAT for sample similarity and pseudo-time analysis^2^. The major fibroblast clusters were identified (Fig. 6M). The mesodermal cells were dispersed throughout the other clusters suggesting that the mesodermal cells may be pluripotent progenitor cells. E12.5 pre-lipofibroblasts and E17.5 lipofibroblasts were closely associated but did not integrate (Fig. 6M). This implies a direct hierarchical relation between these two clusters. This was confirmed by pseudotime analysis (Fig. 6N). Myofibroblasts and intermediate fibroblasts from E12.5 integrated with the corresponding subpopulation from the E17.5 dataset (Fig. 6M-N). This may suggest that these cell types were terminally differentiated cells at the earlier embryonic stage. E17.5 *Ebf1*^+^ fibroblasts were separated into two sub-clusters and showed greater differentiation potential compared to myofibroblasts and intermediate fibroblasts (Fig. 6M-N). This supports our observation that *Ebf1*^+^ fibroblasts and pericytes in the adult lung may be related and it is possible that these two populations, at E17.5, are the progenitors of the corresponding population in the adult lung.

Furthermore, the genetic programs in mesenchymal subpopulations were confirmed with scATAC-seq analysis (Fig. S3J). Transcription factors and growth factors were identified from single cell RNA-seq were confirmed with scATAC-seq analysis (Fig. S3J).

To summarize the genetic program of the mesenchymal subpopulations we identified transcription factors and growth factors specific for each cluster that were conserved between time points. The conserved transcription factors and growth factors identified are illustrated in Fig. 6O.

### scRNA-seq on human lungs identified up to eight fibroblast subtypes

Single cell lung suspensions of explanted healthy and IPF donor lung tissue were generated. All live cells were sorted by FACS and scRNA-seq performed, as described for the murine lung data, on the *EPCAM* negative population. After QC, the mesenchymal cells were identified using canonical markers and subset for further analysis (Fig. S30A-E). The results of scRNA-seq on P1, month 21, healthy and IPF donor human lung tissue from publicly available datasets were re-analysed and integrated, where appropriate, with the scRNA-seq data generated in our laboratory (Fig. S30F)^4,10,13,27^. Patient sample characteristics and human scRNA-seq dataset details are summarised in Supplementary Table 2. Up to eight mesenchymal subpopulations in each data set were identified (Fig. 7A-D). We consistently identified lipofibroblasts, myofibroblasts, SMCs, pericytes, a population with a homologous transcriptomic profile of the murine *Ebf1*^+^ fibroblasts, an intermediate fibroblast subtype and mesothelial cells. The human mesenchymal subpopulations had a distinct and highly conserved transcriptomic profile that was remarkably similar to the corresponding murine lung subpopulation (Fig. 7E-H).

**Figure 7.**
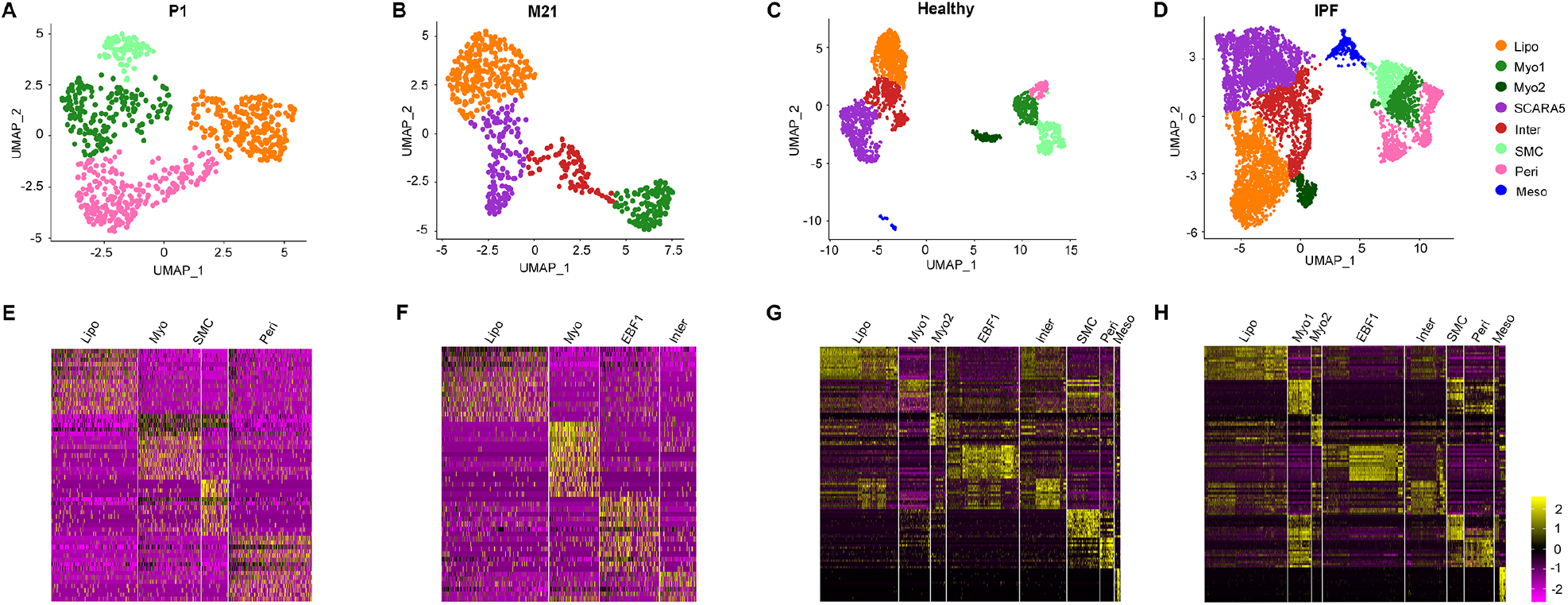
Identification of lung fibroblast subtypes in the human lung. UMAP visualization of mesenchymal subtypes in P1 (**A**), M21 (**B**), healthy control (**C**) and IPF donor (**D**) lungs. Heatmap of scaled gene expression of the top 15 differentially expressed genes (rows) in each cluster of cells (columns) in P1 (**E**), M21 (**F**), control (**G**) and IPF (**H**) dataset. Lipo, lipofibroblasts, Myo, myofibroblasts, *EBF1*, *EBF1* subpopulation, Inter, intermediate fibroblasts, SMC, smooth muscle cells, Peri, pericytes, Meso, mesothelial cells.

### Lipofibroblasts in the human lung

As we observed in the murine lung, canonical lipofibroblasts markers with the exception of *TCF21*, were found to poorly discriminate the lipofibroblast cluster identified in the human lung (Fig. 8A, B). Novel marker genes represented by *A2M* (alpha-2-macroglobulin)*, RARRES2* (retinoic acid receptor responder 2, chimerin) and *GPC3* (Glypican 3), and those identified in the murine lipofibroblasts represented by *LIMCH1, MACF1*, better delineated the human lipofibroblast cluster than canonical markers (Fig. 8B, C). The top differentially expressed genes in human lipofibroblasts in each dataset, in comparison to all other mesenchymal cells, were determined using the MAST statistical framework. These genes were visualised using volcano plots and consistently included *TCF21, LIMCH1, A2M, RGCC* (regulator of cell cycle) and genes related to reported lipofibroblast functions, for example lipid/retinoic acid processing and/or storage (Fig. 8D). Comparative analysis of lipofibroblasts from control and IPF donor lungs identified that collagen and ECM related genes were among the most differentially expressed (Fig. 8E). The most highly expressed and specific transcription factors in lipofibroblasts conserved between datasets were *TCF21, NR2F1* (nuclear receptor subfamily 2 group F member 1, also known as COUP-TF1) and *LMO4* (LIM domain only 4) (Fig. 8F). The complete differentially expressed transcriptomic profile of human lipofibroblasts in each dataset is available in Supplementary Table 3.

**Figure 8.**
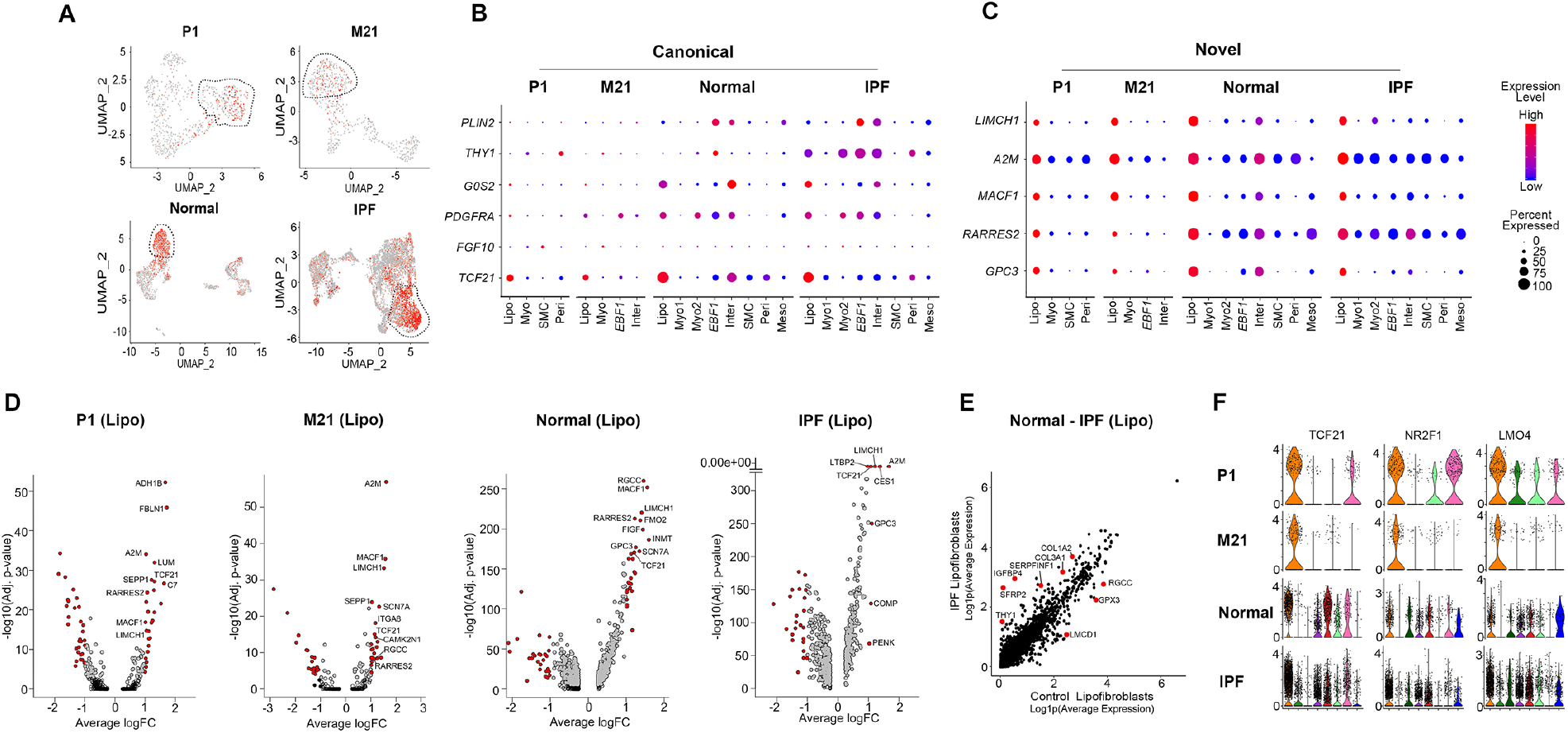
Identification of human lung lipofibroblasts and lipofibroblast specific markers. (**A**) UMAP visualization of *TCF21* (lipofibroblast cluster circled). Dot plot visualization of canonical (**B**) and novel (**C**) lipofibroblast marker genes. Dot size corresponds to the percentage of cells expressing the gene and color to average expression level. (**D**) Volcano plot visualization of differentially expressed lipofibroblast genes. Genes in red, p < 10^−5^; average log fold-change (Avg_logFC)> 1, Genes in black, p < 10^−5^; Avg_logFC < 1, Genes in grey, p> 10^−5^; Avg_logFC > 1. (**E**) Comparative analysis of changes in gene expression in lipofibroblasts from healthy vs IPF donor lungs. (**F**) Violin plot representation of conserved transcription factors in the lipofibroblast cluster. Lipo, lipofibroblasts, M, month, P, post-natal day.

### Myofibroblasts and SMCs in human lungs

Myofibroblasts and SMCs were closely associated and had a homologous transcriptomic profile with common markers, for instance *ACTA2 and TAGLN*, prominently expressed in both populations (Fig. 9A-C). In the P1 and healthy donor lung datasets the SMC and myofibroblast clusters were distinct (Fig. 9A, B). In the IPF lung myofibroblasts also prominently expressed myosin heavy chain genes, like MHY11, and increased their expression of other genes traditionally associated exclusively with SMCs (Fig. 9C, D). Specific, delineating, marker genes to differentiate the human myofibroblast cluster from SMCs could not be identified. SMCs could be discriminated from myofibroblasts using some commonly reported marker genes represented by *CNN1, SYNPO2* (synaptopodin-2)*, ACTG2*, in the non-fibrotic datasets (Figure 9D, E). In addition to commonly used SMC markers, the P1 dataset expressed the reported human airway SMC marker *HHIP* while other datasets expressed reported vascular SMC markers, *NTRK3* (Neurotrophic Receptor Tyrosine Kinase 3) and *MEF2C* (myocyte enhancer factor 2C) (Fig. 9D)^34^. In the adult human lung two myofibroblast subpopulations were identified (Fig. 7C-D). The first (Myo1), highly expressed commonly reported myofibroblast marker genes. The second (Myo2), was distinct, expressed *ACTA2*, and increased expression of *TGFBI* in IPF. Myo2 had a gene profile homologous to that of “Classical Myofibroblasts” in a recent pre-print publication^12^. The top differentially expressed genes determined using MAST in the myofibroblast clusters were visualized by volcano plot (Fig. 9F, G). Commonly reported SMC marker genes, for example *CNN1, SYNPO2*, were consistently among the top differentially expressed genes determined using the MAST statistical framework in the SMC clusters (Figure. 9D, E, H).

**Figure 9.**
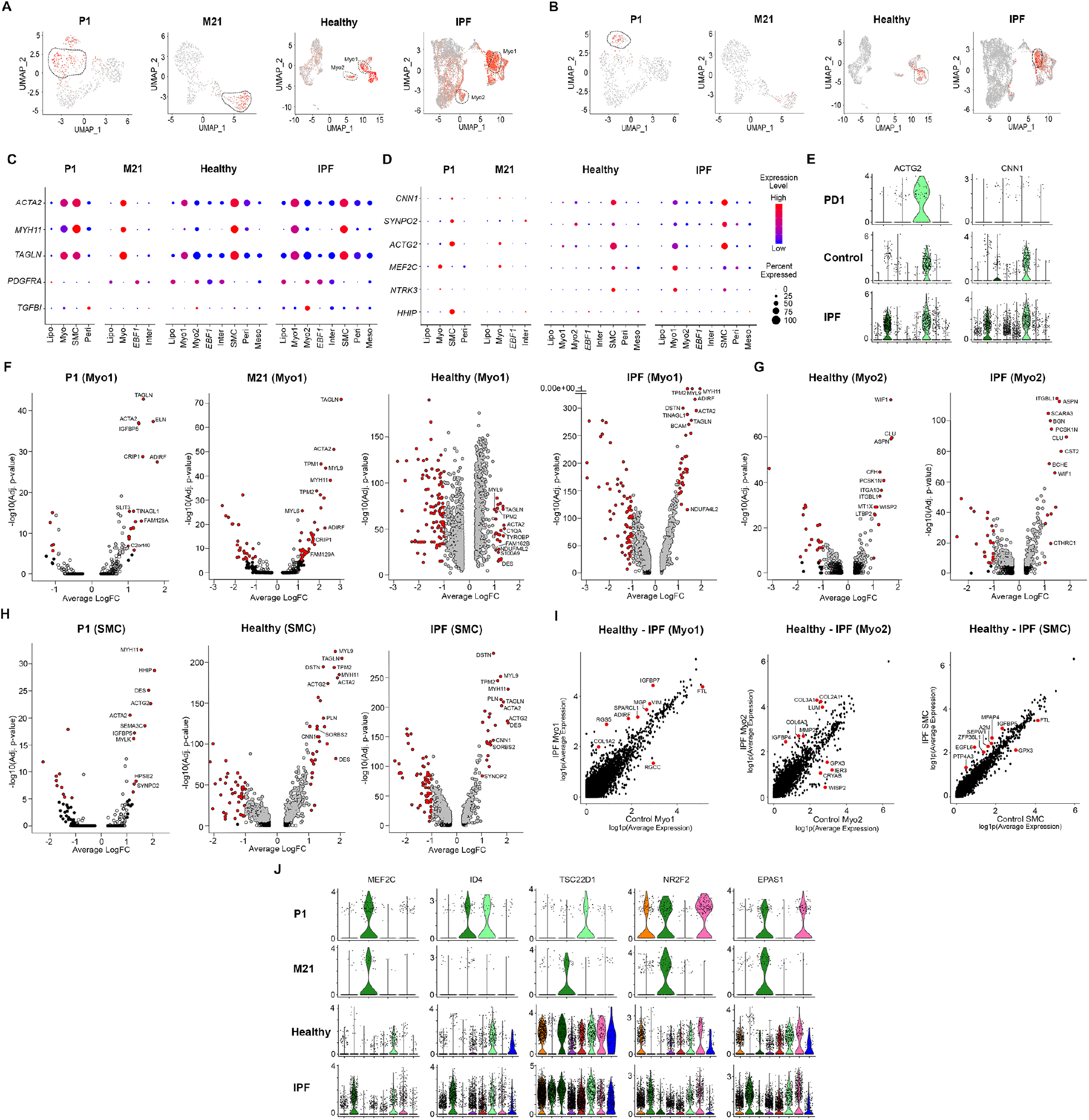
Identification of human lung myofibroblast and SMC clusters. UMAP visualization of *TAGLN* (myofibroblast clusters circled) (**A**) and *ACTG2* (**B**) (SMC cluster circled). Dot plot representation of myofibroblast (**C**) and SMC (**D**) markers. Dot size corresponds to percentage of cells expressing the gene and color to average expression level. (**E**) Violin plot representation of widely reported SMC genes. Volcano plot visualization of differentially expressed myofibroblast (**F**, **G**) and SMC (**H**) cluster genes. Genes in red, p < 10^−5^; average log fold-change (Avg_logFC) > 1. Genes in black p < 10^−5^; Avg_logFC < 1. Genes in grey, p> 10^−5^; Avg_logFC > 1. (**I**) Comparative analysis of changes in gene expression in myofibroblast and SMC subpopulations from healthy vs IPF donor lungs. (**J**) Violin plot representation of conserved transcription factors in the SMC/myofibroblasts clusters. Myo, myofibroblasts, M, month, P, post-natal day, SMC, smooth muscle cell.

Comparative analysis of the differentially expressed genes in the myofibroblast and SMC clusters from healthy and IPF donor lungs identified genes related to ECM production (for example *VIM*, *COL2A1*, *COL3A1*), matrix metalloproteinase genes (for example, *MMP2*) and IGF binding proteins (for example *IGFBP4*, *6*) (Fig. 9I). The top differentially expressed genes transcription factors conserved between datasets were *MEF2C* (myocyte-specific enhancer factor 2C), *ID4* (inhibitor of differentiation 4), *TSC22D1* (TSC22 domain family member 1), *NR2F2* (nuclear receptor subfamily 2 group F member 2) and *EPAS1* (endothelial PAS domain-containing protein 1 (Fig. 9J). The complete, differentially expressed, transcriptomic profile of human myofibroblasts and SMCs in each dataset is available in Supplementary Table 3.

### Pericytes and *Ebf1* fibroblasts in the human lung

Discrete expression of genes commonly associated with pericytes were identified in three of the four datasets (Fig. 10A, C). Conserved expression of a further list of possible novel pericyte marker genes in these clusters represented by *NDUFA4L2* (NADH dehydrogenase 1 alpha subcomplex, 4-like 2), *PAG1* (phosphoprotein associated with glycosphingolipid-enriched microdomains 1) and *FAM162B* (family with sequence similarity 162 member B) among others were identified (Fig. 10C). The top differentially expressed genes, determined using MAST, in the pericyte cluster were visualised using volcano plots and included known pericyte marker genes and genes identified as potential novel markers (Fig. 10E).

**Figure 10.**
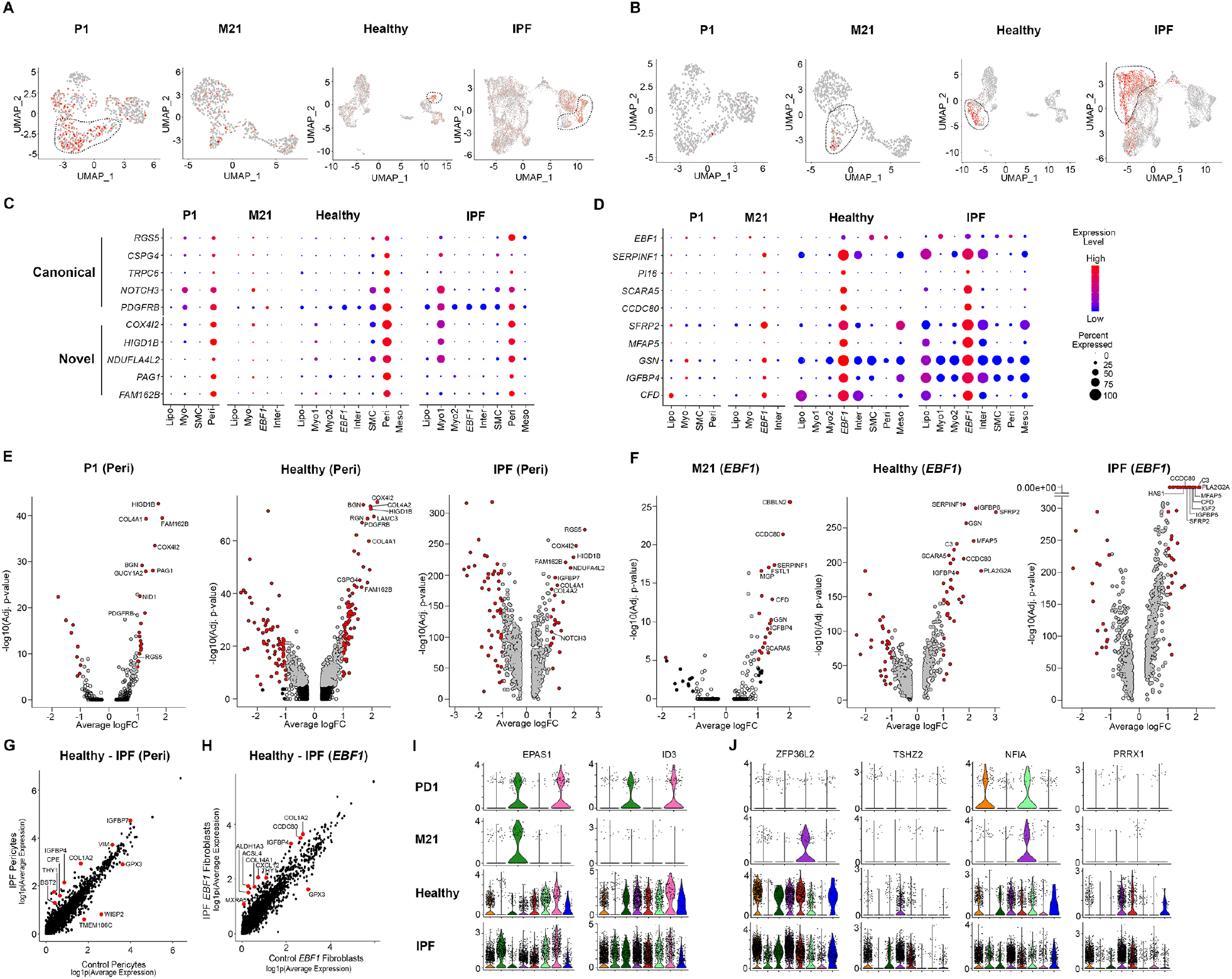
Identification of human lung pericytes and *EBF1* fibroblasts. UMAP visualization of *PDGFRB* (pericyte cluster circled) (**A**) and *SCARA5* (*EBF1* subpopulation circled) (**B**). Dot plot representation of canonical and novel pericyte genes (**C**) and the *EBF1* subpopulation transcriptomic signature (**D**). Dot size corresponds to percentage of cells expressing the gene and color to average expression level. Volcano plot visualization of differentially expressed pericyte (**E**) and *EBF1* (**F**) cluster genes. Genes in red: p < 10^−5^; average log fold-change (Avg_logFC) > 1. Genes in black, p < 10^−5^; Avg_logFC < 1. Genes in grey, p> 10^−5^; Avg_logFC> 1. Comparative analysis of changes in gene expression in pericytes (**G**) and *EBF1* (**H**) fibroblasts from healthy vs IPF donor lungs. Violin plot representation of conserved transcription factors in the pericyte (**I**) and EBF1 (**J**) clusters. M, month, Peri, pericyte, P, post-natal day.

A mesenchymal population with a homologous transcriptomic signature to the novel *Ebf1*^+^ subpopulation in the murine lung was identified in three of four human datasets (Fig. 10B, D). The top differentially expressed genes in this cluster, determined using MAST, in each dataset were visualised using volcano plots (Fig. 10F). The most significant genes in this cluster were also prominent in the corresponding murine population, such as *SCARA5* (scavenger receptor class A member 5) and *SERPINF1* (serpin family F member 1), but also included genes prominent in humans but not mice, for example *CCDC80* (coiled-coil domain containing 80) (Fig. 10D, F). Comparative analysis of differentially expressed genes in pericytes and *EBF1* fibroblasts from healthy and IPF donor lungs included CXCL chemokine and ECM related genes (for example *COL1A2*, *COL14A1*) (Fig. 10G). *EPAS1* and *ID3*, were the only transcription factors with a relatively discrete expression among the differentially expressed pericyte genes (Fig. 10I). *ZFP36L2* (zinc finger protein 36 C3H1 type-Like 2), *TSHZ2* (teashirt zinc finger homeobox 2), *NFIA* (nuclear factor 1A) and *PRRX1* (paired related homeobox 1) were consistently identified among the top genes in the *EBF1* clusters (Fig. 10J) The complete, differentially expressed, transcriptomic profile of human pericytes and the *EBF1* sub-population in each dataset is available in Supplementary Table 3.

### Expression of ECM related genes in the IPF lung

All healthy and IPF donor lung mesenchymal cells were integrated (Fig. 11A, B). The major subpopulations were identified (Fig. 11C). ECM related gene expression was significantly increased in mesenchymal cells from IPF lungs in comparison to the cells from healthy donors (Fig. 11D). Consistent with our data in the murine lung, all mesenchymal subpopulations identified, not solely myofibroblasts, increased their expression of major ECM related genes as represented by *COL1A1*, *COL1A2*, *COL3A1* and *FN1* (Fig. 11E).

**Figure 11.**
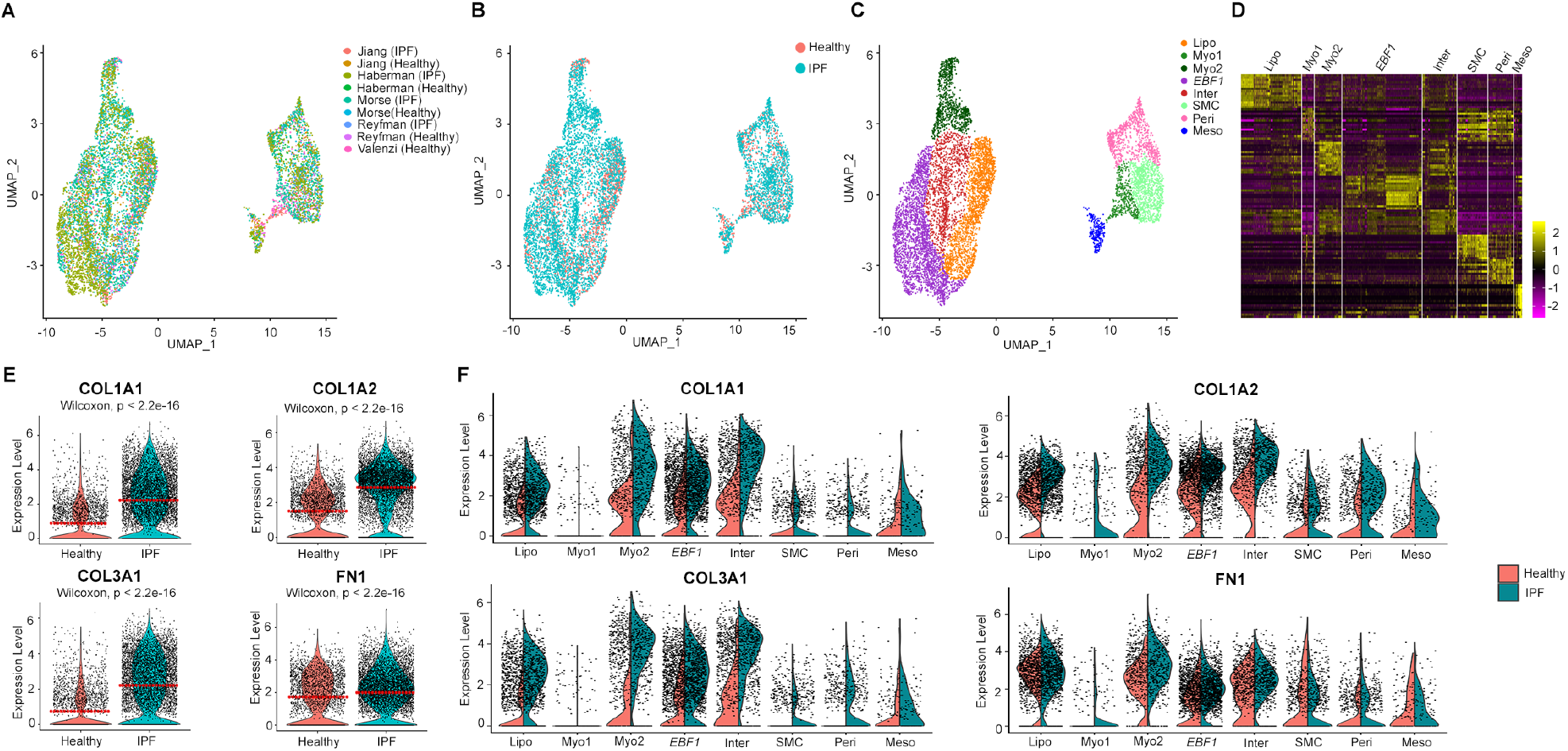
Expression of extracellular matrix associated genes in the human lung. (**A**, **B**, **C**) UMAP visualization of the integrated and clustered mesenchymal cells from normal/IPF donor lungs and published data. Violin plot representation of representative ECM related genes in healthy vs. IPF donor lung mesenchymal cells (**D**) and the change in expression of these genes in each identified cluster (**E**). Wilcoxon, p < 2.2e-16 per comparison. Lipo, lipofibroblasts, Myo, myofibroblast subpopulation, *EBF1*, *EBF1* subpopulation, Inter, intermediate fibroblasts, SMC, smooth muscle cells, Peri, pericytes, Meso, mesothelial cells.

## Discussion

Due to recent advances in omic technologies, and the application of techniques like scRNA-seq and scATAC-seq to the lung, we have begun to appreciate that mesenchymal cells are a conglomeration of distinct subpopulations and not a homogenous population delineated by collagens, such as Col1a1, Col3a1 etc., or Pdgfr-*α* or -*β* expression. Despite this marker genes with which to indicate distinct cell types, a lack of information on subpopulation origin(s) and functions remain. To date there is surprisingly little information on the molecular identity, localization and relative abundance of different stromal populations^35^. There is little if any consensus in the literature on differential expression, by these cells, of widely reported markers in health and disease. Therefore, in this comprehensive longitudinal study, we have addressed this gap in our understanding of mesenchymal cells in the healthy and fibrotic lung by analyzing the transcriptome of all mesenchymal cells as they emerged in the lung during embryonic development, tracked the identified populations to adulthood and examined the changes in gene expression in the fibrotic lung. We have systematically investigated the expression of all commonly reported mesenchymal markers and demonstrated that many of these markers are poorly discriminative for the target population. Our comparative analysis of the changes in gene expression in each mesenchymal subpopulation in the fibrotic lung of both mice and humans suggests all mesenchymal subtypes contribute to ECM production in fibrosis. Further, these data suggest that there is little evidence of trans-differentiation of fibroblast subtypes and that fibroblast fate is determined early in lung development.

### Pulmonary lipofibroblasts in lungs

Lipofibroblasts have been consistently reported in the rodent lung but rarely in the human lung leading to controversy in the literature regarding their existence, identification, and relevance to human disease^20,21^. Traditionally lipofibroblasts have been identified histologically by the presence of intracellular lipid droplets, markers of an adipose like phenotype, enzymatic properties, characteristic cytokines, and canonical marker genes like *Plin2*, *Lpl*, and *Fgf10* among others^20,36^. In recent lineage tracing studies, *Tcf21* was demonstrated to be preferentially expressed in adult murine lung lipofibroblasts^27^. Previously, both the *Pdgfra*^+^ and the *Fgf10*^+^ lineage lung stromal cell population were reported to include lipofibroblasts^7,37,38^. The use of lipid dyes, and/or associated genes like *PLIN2*, to distinguish and quantitate lipofibroblasts in the lung is not ideal. Lipid droplets exist in virtually all normal cells and closely associated resident lung cell types like macrophages, endothelial cells, mast cells and AEC2s express canonical lipofibroblast marker genes like *PLIN2*^39–44^. Further PPRAγ signaling, traditionally associated with lipofibroblasts, is prominent in both macrophages and epithelial cells and proteins related to lipid metabolism and handling are also expressed prominently by alveolar macrophages^43–48^. *In vitro* studies, in which an overt lipofibroblast like phenotype has been induced in cultured fibroblasts, typically using PPARγ agonists have likely contributed to the use of these markers. Traditional lipofibroblast markers can be readily detected at both the mRNA and protein level in stimulated fibroblasts in which a lipofibroblast-like phenotype has been induced.

In the present study, we found that canonical lipofibroblast marker genes delineated a population of fibroblasts clearly in the rodent lung between E16.5-E17.5, when lipofibroblasts emerge and are readily detectable, and P15, reportedly when the prevalence of lipofibroblasts in the rodent lung peaks^49,50^. These genes were less effective at later developmental stages in the rodent lung and, other than *TCF21*, ineffective at identifying the lipofibroblast cluster in humans. Our novel lipofibroblast signature, in keeping with the recent lineage tracing study, included *Tcf21/TCF21* and was consistently discriminative for the associated, transcriptomically distinct, cluster of cells in all datasets^8^. The top differentially expressed genes in bulk sequenced fibroblasts sorted using the novel lipofibroblast cell surface markers (Cd249) identified in the scRNA-seq analysis overlapped substantially with the transcriptomic signature of lipofibroblasts we consistently identified. The postulated lipofibroblast lineage marker *Fgf10* was expressed in murine, but not human, lipofibroblasts supporting the findings of a previous publication^7^. *Pdgfra/PDGFRA* expression was not consistently expressed in any single cluster in mouse or human mesenchymal cells. A limited proportion of human lipofibroblasts were *PDGFRA*^+^ and supporting the notion that Pdgfra^+^ fibroblasts include lipofibroblasts in the adult murine lung lipofibroblast cluster *Pdgfra* expression was prominent^37^. Fibroblasts, in which a lipofibroblast-like phenotype had been induced using conventional methods reported in the literature, displayed an overt lipofibroblast phenotype, prominent lipid inclusions, and significant expression of canonical marker genes. These cells were supportive of AEC2 3D organoid growth as reported for *Pdgfra*^+^ lipofibroblasts*^37^*. However, these *in vitro* cells did not display a similar transcriptomic profile to lipofibroblasts *in vivo*.

These data suggest that the canonical markers are only efficacious at identifying lipofibroblasts in the murine lung at specific developmental stages and ineffective at delineating lipofibroblasts in the human post-natal lung. Traditional marker genes are a prominent feature of *in vitro* lipofibroblast like cells but the transcriptomic signature of cultured lipofibroblast like cells is not similar to lipofibroblasts *in vivo.* Our findings may explain the difficulty in identifying human lung lipofibroblasts *in vivo* using conventional marker genes and support the recent lineage tracing study documenting *Tcf21* as a lipofibroblast lineage marker.

### Myofibroblasts and SMCs

Myofibroblasts have long been considered the primary drivers of ECM deposition in fibrosis, and the key effector cells in IPF combining of the synthesizing features of fibroblasts with the cytoskeletal contractile characteristics of SMCs^1,9,16,22^. The definition of myofibroblasts however has been almost entirely dependent on *α*SMA (*Acta2*) expression. As noted repeatedly in the literature myofibroblasts and SMCs express a number of common cell selective markers like *α*SMA (*Acta2*), SM22*α*(*Tagln*), desmin (*Des*) and vimentin (*Vim*) among others^16,22,51–53^. As myofibroblasts are also reportedly capable of producing calponin, encoded by *Cnn1*, and numerous other putative SMC markers it has been virtually impossible to distinguish myofibroblasts from true SMCs. In our analysis we also found this to be the case when using commonly reported myofibroblast/SMC markers. Even reportedly SMC “specific” markers like myosin heavy chain genes, for instance *Mhy11/MYH11*, were prominently expressed in both cell types^16,54^. Widely used markers were not discriminative for either population or the expression of these genes was comparable in both populations. This was particularly evident in the IPF lung, and aged fibrotic murine lung, where gene expression in these populations was highly homologous.

In the adult, healthy, lungs of both mice and humans we found that myofibroblasts and MCs clustered distinctly. We successfully identified discriminative marker genes for myofibroblasts in the murine lung. This was not the case in the human lung where the transcriptomic differences between myofibroblasts and SMCs were either very subtle or non-existent as suggested by others^53^. The distinct myofibroblast clusters were *Thy1/THY1*^−^ as suggested by Sanders et al. but the postulated myofibroblast marker S100A4 was expressed by all mesenchymal subtypes^22,52,54^. Neither *Pdgfra* nor *Pdgfrb* expression were discriminative for myofibroblasts in mice in keeping with the observations of previous publication, which both reported that Acta2^+^ cells were Pdgfra^− 22,30,55^. Similarly, in the human lung *PDGFRA* and *PDGFRB* expression was not discriminative for myofibroblasts. However, the expression of these genes did increase in IPF myofibroblasts. This phenomenon not observed in the fibrotic murine lung.

We identified a number of SMC associated genes that displayed discrete expression in the SMC clusters, like *Actg2/ATCG2* in both species, *Actc1* in mice, or *NTRK3/MEF2C* in humans^16,34,56^. Therefore, it was possible, using a select number of reported SMC markers, to distinguish SMCs from myofibroblasts. It should be noted that some of these genes, for instance *Hhip/HHIP* appear to be species specific. *Hhip* displayed discrete expression in murine myofibroblasts. In humans, as previously reported, *HHIP* displayed discrete expression in the SMC cluster alongside traditional SMC markers^34^.

SMCs and myofibroblasts are believed to share common lineage(s) with *Fgf10, Axin2, Gli1* and *Wt1* lineages all suggested to include myofibroblasts^56–59^. The contribution of these lineages to the distinct subsets is yet to be definitively resolved with reports leaning towards *Fgf10/Wt1*^+^ cells as predominantly fibroblast/mesothelial and *Gli1/Axin2* as the predominantly giving rise to myofibroblast/SMCs^7,56,59^. These studies are frequently limited by the dependence on *α*SMA or *Acta2* as the marker for myofibroblasts and/or SMCs. As discussed, this marker is not discriminative. Authors have long equated the increase in *α* SMA^+^ cells with contractile myofibroblasts^60^. We noted increased *Acta2/ACTA2* expression in the fibrotic murine and human lung as reported by others. However, our data does not support the hypothesis that this increase can be attributed to an expansion of the myofibroblast population^22^. We observed an increase in *Acta2/ACTA2* expression in multiple mesenchymal subtypes without an associated increase in myofibroblast number.

Our findings highlight that continued reliance on traditional myofibroblast/SMC markers is likely to yield ambiguous data, as expression of any one marker, like *α*SMA^+^, is likely to be dynamic or species specific. SMCs can be discriminated from myofibroblasts if the correct markers are selected. These data advocate for a strategy of using multiple markers for subpopulation discrimination, as conducted by a recent publication, which combined fluorescence *in situ* hybridization (FISH) co-localization of a traditional marker (*ACTA2*) with novel subtype specific markers. Using this approach in future lineage tracing studies will likely yield a more definitive answer on myofibroblast/SMC lineage. Further, spatial transcriptomics studies may be required to definitively determine a myofibroblast specific signature with which fractionate these cells.

### *Ebf1*^+^ mesenchymal cells and pericytes

We identified a novel mesenchymal subpopulation delineated by *Ebf1* in the E17.5 murine lung with a transcriptomic signature that could not be attributed to any known mesenchymal subtype. In the embryonic lung, this population co-expressed markers for pericytes. In the post-natal lung, the *Ebf1*^+^ populations diverged and became distinct. The first *Ebf1*^+^ population displayed discrete expression of known pericyte markers. The second, was closely associated with the other fibroblast subtypes, had a unique transcriptomic signature, and could be delineated in most datasets. These data suggest that the novel *Ebf1*^+^ fibroblast population and pericytes may share a common developmental lineage. Alternatively, in the embryonic datasets pericytes and *Ebf1*^+^ fibroblasts were indivisible due to a common transcriptomic and therefore clustered together. In the human post-natal lung, a mesenchymal population with a highly homologous transcriptomic signature to the murine *Ebf1*^+^ fibroblasts were identified along with a distinct pericyte cluster. As embryonic human data was unavailable for this study it was not possible to determine if the divergence of the *EBF1* and pericyte populations was also evident in the human lung.

There is little in the literature on the role of *Ebf1* in fibroblasts. However, in a recent study an *Ebf1^high^* fibroblast population was identified as a distinct cluster in a scRNA-seq analysis of wound fibroblasts^61^. Recent pre-print publications identified an “adventitial fibroblast” subtype with a similar transcriptomic signature to the *Ebf1/EBF1* population in our study^11,12^. The *in-situ* hybridization localization of *SFRP2, SERPINF1, PI16* prominent genes in the *Ebf1/EBF1* cluster we identify in a recent study are compatible with the results of our Ebf1 immunofluorescence localizing a proportion of Ebf1^+^ fibroblasts to the adventitia^12^. Ebf1 deletion was demonstrated to have critical effects on Foxd1^+^ stromal progenitors, a lineage that includes pericytes^62,63^. Further reports document that cells expressing the pericyte marker Ng2^+^ (aka Cspg4) require Ebf1 for their function and a recent study reported an *Rgs5^+^* subgroup of PDGFR *β* pericytes with a transcriptomic signature characterized by *Ebf1*, as well as *Ndufla4l2, Cox4i2* and *Higd1* all genes we identify as discrete pericyte markers^64,65^. These reports are supportive of our identification of an *Ebf1^+^/EBF1* fibroblast population as a distinct subtype and our hypothesis that this subtype and pericytes may share a common lineage.

### Pericytes in fibrotic lungs

We found that commonly reported pericyte markers identified a distinct cluster of cells in the adult murine and human lungs. However, transcript expression of *Cspg4/CSPG4* and *Rgs5/RGS5*, prototypical pericyte marker genes, were low in both murine and human lung mesenchymal cells while *Pdgfrb/PDGFRB* had high background expression in almost all other mesenchymal subtypes ^22,55,66^. More novel markers were expressed at greater levels and were more discriminative for pericytes. We did not observe a *Pdgfra/PDGFRA*^+^ pericyte population as reported by a previous study ^66^. It has been reported that pericytes (for example Gli1^+^, Foxd1^+^ lineage, Ng2^+^ and Foxj1^+^ cells) may give rise to alpha-SMA^+^ myofibroblasts during fibrosis and/or acquire a myofibroblast like phenotype^22,33,55,63,66^. In our analysis transcript expression of myofibroblast like genes, like *Acta2* and *Myh11*, increased in the adult and aged fibrotic murine lung pericyte cluster consistent with these reports. This increase in “myofibroblast” gene expression was not clear in the distinct human pericyte cluster. However, expression of pericyte markers, like *RGS5, PDGFRB*, and *NOTCH3*, became pronounced in the IPF myofibroblast/SMC clusters. It is possible that some pericytes may have acquired a myofibroblast like phenotype, as described by others, and therefore have clustered with myofibroblasts in our human analysis.

### ECM in fibrotic lungs

Myofibroblasts have long been reported as the driver of ECM deposition in the fibrotic lung^1,9,22^. The present study demonstrates that all identified fibroblast subpopulations, not just myofibroblasts, increase their expression of transcripts for ECM components; collagens (*COL1A1, COL1A2, COL3A1*) and fibronectin (*FN1*), in both the fibrotic murine and IPF lung. These data are supportive of the previous work, which reported a dramatic expansion of Col-EGFP^+^ cells in the bleomycin injured lung, with only a minority of cells expressing both Col-EGFP and Acta2-RFP^60^. They are also in keeping with a growing body of research challenging the assumption that *α*SMA is a consistent marker of collagen producing cells, and the focus on the myofibroblast as the major pathological cell type in IPF^22,57,60^.

### Commonly used markers

We previously reported that *Col14a1* and *Col13a1* represented distinct matrix fibroblast clusters in adult and fibrosis mouse lungs^2^. Here, in all murine lung mesenchymal cells, we found that *Col13a1* was expressed alongside the known and novel lipofibroblast specific genes at all timepoints and *Col14a1* was mostly expressed in lipofibroblast clusters in embryonic and P1 lungs and in *Ebf1*^+^ clusters in later postnatal, adult and aged lungs. *Col14a1* and *Col13a1*, previously reported as matrix fibroblast markers, have been demonstrated in this comprehensive analysis as discriminative for distinct mesenchymal subtypes. While these findings is at odds with our previous report^2^, this analysis benefits from a greater number of cells in the adult dataset, integration with independent datasets, and a longitudinal analysis. PDGFRA and PDGFRB are commonly used to differentiate distinct mesenchymal subtypes. In our study, *Pdgfrb* expression was well overlapped with *Pdgfra* expression in some datasets in the murine lung while in others the expression of these two genes was well separated with high background overall. In adult and fibrotic mouse lungs, *Pdgfra* was well separated from *Acta2^+^* cells and *Pdgfrb^+^* cells, but showed good overlap with *Tcf21^+^* lipofibroblasts, which denied that it as a marker of myofibroblasts in adult lungs. *Pdgfrb* showed some overlap with one of the *Ebf1* expressing fibroblast clusters, pericytes, with high background in other clusters. This suggested that Pdgfrb could be used as a pericyte marker, but its specification at protein level needs to be further validated. Vimentin is an often used as mesenchymal cell marker and in some instances reportedly a gene specific for myofibroblast^22^. However, in out scRNA-seq datasets, *Vim* transcript expression was highest in endothelial cells and was detectable in mesenchymal and immune cells. It was rarely detected in epithelial cells. This suggests that *Vim* should not be used as mesenchymal cell marker. More comprehensive lineage tracing experiments using subpopulation-specific transcription factors for each population are needed to verify these definitions.

### Conclusion

This study is the first definitive description of the transcriptome of all mesenchymal subtypes from embryonic development, to adulthood and in the fibrotic lung. We have demonstrated that mesenchymal fate decisions occur during embryonic development and the identified transcriptomic signatures remain distinct into adulthood and in the aged lung. We did not find evidence of trans-differentiation between mesenchymal subtypes even in the diseased lung. Comparative analysis between the murine and human lung demonstrated that the transcriptomic signature of the subtypes is remarkably conserved between species. Canonical markers, in general, were poorly discriminative for their associated mesenchymal subtype. Novel markers we have identified were consistently discriminative for each subtype irrespective of developmental stage. This comprehensive analysis provides a wealth of new markers and transcriptomic information with which to study these cell types and will enhance the study of mesenchymal cells in health and disease.

## Materials and Methods

### Study approval

The use of human tissues for research were approved by the Institutional Review Board (IRB) of Cedars-Sinai Medical Center and were under the guidelines outlined by the IRB (Pro00032727). All animal experiments performed in this study were approved by Cedars-Sinai Medical Center Institutional Animal Care and Use Committee (IACUC008529).

### Bleomycin instillation

Detailed methods can be found in Supplementary Methods.

### Mouse lung tissue isolation

Wild-type C57/Bl6J mice from an in-house colony were used in all experiments. Animals were randomly assigned to treatment groups. Animals of both genders were used without bias. Mice were considered adult at 8-to 12-weeks-old and aged at between 82-to 95-weeks-old. All mice had access to autoclaved water and pelleted mouse diet *ad libitum* were housed in a pathogen free facility at Cedars-Sinai Medical Center. For the isolation of embryonic murine lung tissues breeding cages; containing a male and two female mice, were monitored intensively following the addition of the male to the breeding cage. The presence of a female with a vaginal plug was considered embryonic day 0.5 (E0.5). Adult (12-16weeeks old), aged (82-95 weeks old), or pregnant mice were deeply anaesthetized by intraperitoneal injection (I.P.) of Ketamine (100mg/kg) and Xylazine (10 mg/kg) followed by exsanguination. Adequate depth of anesthesia was determined by lack of a withdrawal reflex to paw, followed by tail pinch, prior to the start of any surgical intervention. In adult mice the lungs were cleared of blood by flushing phosphate buffered saline (#10010023, Gibco, Thermo Fisher Scientific, Waltham, MA, USA) through the pulmonary artery via cardiac puncture prior to isolation. Pregnant mice were euthanized at the indicated time and the embryos were quickly isolated after removal of the uterus, and the lungs of the embryos were resected. The lungs of embryos, adult, and aged mice were transferred to a 15ml tube containing ice-cold PBS and processed immediately.

### Murine lung dissociation and cell isolation

Murine lung tissues were dissociated using a standard protocol in our laboratory that we have reported previously^67^. Isolated tissues were taken immediately to a sterile laminar flow tissue culture hood where they were, rinsed in fresh PBS and then minced finely using a scissors in a 100mm^2^ Petri dish. The minced lung tissue was then suspended in a digestion media containing 0.125% vol/vol Trypsin-EDTA (#25300056, Gibco, Thermo Fisher Scientific, Waltham, MA, USA), 1mg/ml Bovine Serum Albumin (#15260037, Gibco, Thermo Fisher Scientific, Waltham, MA, USA), 100 U/ml DNase 1 (#DN25, Sigma Aldrich, St. Louis, MO, USA), 1 mg/ml Collagenase IV (#LS004209, Worthington Biochemical, Lakewood, NJ, USA) and transferred a tissue culture incubator at 37°C for 30 minutes. At 10-minute intervals the lung digestion solution was titurated 10 times using a 10 ml glass pipette. Following the incubation period, the supernatant and remaining tissue was passed through a 100 μm strainer into a 50ml tube. The strainer was washed with Dulbecco’s Modified Eagle Medium (#11995065, Gibco, Thermo Fisher Scientific, Waltham, MA, USA) containing 10% vol/vol fetal bovine serum (#SH3062601, HyClone, GE Healthcare, Chicago, IL USA). The tube was then centrifuged at 1600 rpm for 10 minutes at 4°C and the pellet resuspended in Hank’s Buffered Saline Solution (#14175095, Gibco, Thermo Fisher Scientific, Waltham, MA, USA) containing 0.2 mM EGTA, 10mM HEPES (#15630106, Gibco, Thermo Fisher Scientific, Waltham, MO, USA), 2% vol/vol FBS and 1% vol/vol antibiotic-antimycotic (#15240062, Gibco, Thermo Fisher Scientific, Waltham, MA, USA) referred to hereafter as HBSS^+^. Red blood cells were preferentially lysed by treating the isolated cells with 1X RBC lysis buffer (# 00-4333-57, eBiosceinces, Thermo Fisher Scientific, Waltham, MA, USA) for 45 seconds, followed by immediate dilution in 20 ml HBSS^+^. Cells were centrifuged again and resuspended in fresh HBSS^+^ prior to florescence-activated cell sorting (FACS).

### *In vitro* culture of murine lung fibroblasts

Detailed methods can be found in Supplementary Methods.

### *In vitro* 3D organoid culture with cultured lipofibroblast like cells

Detailed methods can be found in Supplementary Methods.

### Fluorescence activated cell sorting (FACS)

Detailed methods can be found in Supplementary Methods.

### Human lung dissociation and cell isolation

Freshly isolated human lung tissues were obtained from Cedars-Sinai Medical Center and UCLA and were dissociated using a standard protocol in our laboratory^67^. Lung tissue was taken to a sterile tissue culture hood, transferred to a Petri dish and rinsed in PBS. Airways >2 mm were resected from the surrounding tissues and discarded along with the visceral pleura. The remaining tissue was finely minced with a scissors and then a straight razor blade. The minced lung tissue was then washed in Ham’s/F12 media (#11320033, Gibco, Thermo Fisher Scientific, Waltham, MO, USA) at 4°C for 20 minutes to remove blood and then centrifuged at 600 rpm for 5 minutes in a pre-cooled centrifuge. The media was removed, and the tissue transferred to a 50 ml conical tube containing 2 mg/ml Dispase II (Thermo Fisher Scientific, Waltham, MA, USA) in DMEM/F12 overnight at 4°C with gentle agitation. The next day the suspension was heated to 37°C for 30 minutes, and the centrifuged for 5 minutes at 4°C. The supernatant was removed, and any large pieces of tissue finely minced again with a straight razor blade. The tissue was then titurated in a digestion media containing 10 U/ml elastase (#LS002280, Worthington Biochemical, Lakewood, NJ, USA) and incubated for 30 minutes at 37°C. An equal volume of HBSS^+^ was then added, the solution titurated and then centrifuged at (600 g, 5 minutes, 4°C). The supernatant was removed, and the tissue incubated at 37°C for 15 minutes with DNase I solution. The suspension was titurated and transferred to a 70 m cell strainer over a new 50 ml tube. The strainer was rinsed three times with 10 ml HBSS^+^. The suspension was centrifuged (600 g, 5 minutes, 4°C) and the cells resuspended in 1 X RBS lysis buffer for 2 minutes on ice, the solution diluted with HBSS then centrifuged (600 g, 5 minutes, 4°C). The supernatant was removed, and the cells resuspended in appropriate solution for further analysis.

### scRNA-sequencing

mRNA from single cells sorted from lung into lysis plates was reverse transcribed to complementary DNA (cDNA) and amplified as previously described. Library preparation and sequencing were performed as described previously. Sequencing libraries for cDNA from single cells were prepared as per the Single Cell 3′ v2 Reagent Kits User Guide (10x Genomics, Pleasanton, CA, USA). Cellular suspensions were loaded on a Chromium Controller instrument (10x Genomics) to generate single-cell Gel Bead-In-EMulsions (GEMs). GEM-reverse transcription (RT) was performed in a Veriti 96-well thermal cycler (Thermo Fisher Scientific, Waltham, MA, USA). GEMs were collected and the cDNA was amplified and purified with SPRIselect Reagent Kit (Beckman Coulter, Brea, CA, USA). Indexed sequencing libraries were constructed using Chromium Single-Cell 3′ Library Kit for enzymatic fragmentation, end-repair, A-tailing, adapter ligation, ligation cleanup, sample index PCR, and PCR cleanup. The barcoded sequencing libraries were quantified by quantitative PCR using the KAPA Library Quantification Kit for Illumina platforms (KAPA Biosystems, Roche Holding AG, Basel, Switzerland). Sequencing libraries were loaded on a NextSeq500 (Illumina, San Diego, CA, USA) with a custom sequencing setting (26bp for Read 1 and 98bp for Read 2) to obtain a sequencing depth of ~200K reads per cell.

### scRNA-seq data analysis

Detailed scRNA-seq analysis could be found in Supplementary Methods. The demultiplexed raw reads were aligned to the transcriptome using STAR (version 2.5.1) with default parameters, using human GRCh38 (or mouse mm10) transcriptome reference from Ensembl version 84 annotation, containing all protein coding and long non-coding RNA genes. Expression counts for each gene in all samples were collapsed and normalized to unique molecular identifier (UMI) counts using Cell Ranger software version 3.0 (10X Genomics). The result is a large digital expression matrix with cell barcodes as rows and gene identities as columns. Seurat suite version 3.0 was used for downstream analysis. Quality control before analysis on each individual sample were performed on the number of genes detected in each cell (“nFeature_RNA”), number of transcripts detected in each cell (“nCount_RNA”), and percentage of mitochondria related genes (“percent_mt”) in each cell. For clustering, principal-component analysis (PCA), T-distributed Stochastic Neighbor Embedding (t-SNE) and Uniform Manifold Approximation and Projection (UMAP) were performed for dimension reduction. Batch correction was performed if sample integration was needed. Trajectory analysis was performed by package monocle3 as previously described^2^. The bioinformatics methodology is described in full in the Supplementary Methods. Details on the cell numbers pre- and post-QC and the proportion of cells in each of the major factions (Immune, Endothelial, Epithelial, Mesenchymal) in murine and human lung datasets can be seen in Supplementary Table 1 and 2 respectively.

### scATAC-seq and data analysis

Cells from E17.5 murine lung were isolated in the same way as for scRNA-seq. Cell nuclei isolation was optimized from 10x genomics protocols (https://support.10xgenomics.com/single-cell-atac/sample-prep/doc/demonstrated-protocol-nuclei-isolation-from-mouse-brain-tissue-for-single-cell-atac-sequencing) and previous publication^68^. Library preparation was perform following 10x genomics protocols^68^ (https://support.10xgenomics.com/single-cell-atac/library-prep/doc/user-guide-selecting-the-correct-single-cell-atac-user-guide) and sequencing was performed on Illumina® HiSeq 3000/4000. Raw sequencing data is demultiplexed and converted to fastq format by using bcl2fastq v2.20. Cell Ranger ATAC software v1.1.0 (10X Genomics) is used for barcodes identification, reads alignment, duplicate marking, peak calling and cell calling with default parameter. Briefly, each barcode sequence is checked against a ‘whitelist’ of correct barcode sequences, and the frequency of each whitelist barcode is counted. Raw reads are aligned to the human reference genome GRCm38 using BWA-MEM^69^ with default parameters, then duplicated reads that have identical mapping positions on the reference are marked. For peak calling, the number of transposition events at each base-pair along the genome is counted, then signal above a threshold are determined as peak signal after modeling. For cell calling, barcodes with high fraction of fragments overlapping called peak are selected, then odds ratio of 100000 is used to separate the barcodes that correspond to real cells from the non-cell barcodes. Finally, a count matrix is generated consisting of the counts of fragments ends within each peak region for each barcode. For further QC, clustering and gene accessibility visualization were performed following online vignette^70^ (https://satijalab.org/signac/articles/mouse_brain_vignette.html). Briefly, a Seurat object was generated on count matrix and fragments, and QC was performed by removing cells that are outliers for QC metrics: pct_reads_in_peaks, peak_region_fragments, blacklist_ratio, nucleosome_signal. After normalization and linear dimensional reduction, non-linear dimension reduction and clustering, gene accessibilities were visualized by UMAP. Cell types and mesenchymal cell sub-clusters were defined by checking unknown cell type markers.

### Bulk RNA-seq analysis

Detailed methods can be found in Supplementary Methods.

### Histology and Immunofluorescence staining

Detailed methods can be found in Supplementary Methods.

### Bioinformatics Methods

Detailed methods including Read alignments, Quality control, cell clustering, doublet calling and annotation can be found in Supplementary Methods.

### Statistics

The statistical difference between groups in the bioinformatics analysis was calculated using the Wilcoxon Signed-rank test. For the scRNA-seq data the lowest p-value calculated in Seurat was p < 2.2e-10^−16^. For all other data the statistical difference between groups was calculated using GraphPad and the exact value was shown.

#### Data availability

The GEO accession numbers for mouse lung raw and processed scRNA-seq and scATAC-seq data accessed and reported in this paper are listed below: E16.5 mouse lung Tbx4-lineage^+^, a-SMA^+^ cells, GSEXXXXX (in submission); E17.5 mouse lung, GSEXXXXX (in submission); aged mouse lung and aged fibrosis mouse lung sorted mesenchymal cells, GSEXXXXX (in submission); E9.5-E11.5 mouse lung, GSE87038; E12.5 mouse lung CD45^−^ cells, GSE119228; E14.5 mouse lung, GSE108097; P1 mouse lung, GSE122332; P7 and P15 mouse lung Pdgfra-GFP^+^ cells, GSE118555; adult mouse lung, GSE111664, GSE133747, GSE121611, GSE131800 and GSE104154; adult fibrosis lung, GSE131800 and GSE104154.

The GEO accession numbers for human lung raw and processed scRNA-seq data accessed and reported in this paper are listed below: adult human lung, GSEXXXXX (in submission) and IPF human lung, GSEXXXXX (in submission); P1 and M21 human lung, LungMAP: https://lungmap.net/; published adult human lung, GSE135893, GSE128033, GSE122960 and GSE128169; published IPF human lung, GSE135893, GSE128033 and GSE122960. Codes for data procession and analysis are in submission to GitHub at https://github.com/jiang-fibrosis-lab.

## Acknowledgments

The authors thank the members of Noble and Jiang laboratory for support and helpful discussion during the course of the study. This work was supported by National Institutes of Health grants R35-HL150829, R01-HL060539, R01-AI052201, R01-HL077291 (PWN), and R01-HL122068 (DJ and PWN), and P01-HL108793 (PWN and DJ). SCR was supported by European Union’s Horizon 2020 research and innovation program under the Marie Skłodowska-Curie grant agreement 797209. We thank the lung research community for the open sharing of datasets available.

## Author contributions

JL, PWN, and DJ conceived the study. DJ, JL, PWN, XL, and SR designed the study. XL and SR performed most of the experiments and analyzed the data. JL, GH, XL, CY, ND, YW, and DJ analyzed single cell RNA transcriptome data. GH analyzed single cell RNA transcriptome data, performed flow cytometry analysis, and prepared figures. GH, XL, FT, AB, NL, TX, TP, and SR took part in mouse, cell culture, and biological experiments. PC, CH, WCP, and BS analyzed and interpreted data. SSW and JB provided human samples and interpreted data. XL, SR, PWN, and DJ wrote the paper.

## Supplementary Figure Legends

**Figure S1. scRNA-sequencing on E17.5 mouse lung.** Violin plot showing the number of genes (nFeature_RNA), number of read counts (nCount_RNA) and percentage of mitochondria genes (percent.mt) detected in each cell before (**A**) and after QC (**B**). (**C**) Heatmap of top 15 differential expression genes comparing each cell types. (**D**) Dot plot visualization of relative expression of known cell type specific markers of each cell type. (**E**) UMAP visualization of relative expression of known cell type specific markers used for cell type clustering. Mesen, mesenchymal cell, Epi, epithelial cell, Immu, immune cell, Endo, endothelial cell.

**Figure S2. Clustering of E17.5 mouse lung fibroblasts.** UMAP visualization of mesenchymal cell integration (**A**) and clustering (**B**) of the fibroblasts from E17.5 mouse lung. (**C**) UMAP visualization of relative expression of known cell type specific markers to validate the purity of fibroblasts. Visualization of mesenchymal cell clustering (**D**) by t-SNE from E17.5 mouse lung. (**E**) 3-D and 2-D visualization of PCA analysis on the E17.5 lung mesenchymal cells to validate the clustering. Visualization of Pdgfrb transcript by t-SNE (**F**) and UMAP (**G**) from E17.5 mouse lung. (**F**) Dot plot visualization of specific genes of each subpopulation.

**Figure S3. scATAC-seq on E17.5 mouse lung.** Cell quality of the nuclei before (**A**) and after (**B**) QC. Distribution (**C**), cell type gene accessibilities (**D**) and cell types (**E**) of the nuclei visualized by UMAP. (**F**) Heatmap of top 50 genes of each cell type. (**G**) Cluster specific gene accessibilities were visualized by UMAP. (**H**) Mesenchymal cell sub-clusters were defined. (**I**) Heatmap of top 15 genes of each mesenchymal cell cluster. (**J**) Cluster specific growth factors and transcription factors. Lipo, lipofibroblast, Myo, myofibroblast, *Ebf1*^+^, *Ebf1*^+^fibroblast, Inter, intermediate fibroblast, Meso, mesothelial cell.

**Figure S4. Differentiation potential of the E17.5 lung fibroblast subtypes.** (**A**) Metagene profile for each subtype in E17.5 mouse lung fibroblasts. Lineage bifurcation of E17.5 mouse lung fibroblast subtypes by Correlation Spanning Tree (**B**-**C**) and group (**D** -**E**) and k−nearest neighbour graph (k=30) (**F**). Lipo, lipofibroblast, Myo, myofibroblast, *Ebf1^+^*, *Ebf1*^+^ fibroblast, Inter, intermediate fibroblast, Proli, proliferative fibroblast.

**Figure S5. Visualization of IPA analysis of each clusters of E17.5 fibroblasts.** Top 15 activated and inhibited regulators of lipofibroblast (**A**), myofibroblast (**B**), *Ebf1*^+^fibroblast (**C**) and intermediate fibroblast (**D**) of E17.5 mouse lung. Scale bar, 50 μm.

**Figure S6. Clustering of E9.5-E11.5 mouse lung single cells.** (**A**) Cell integration of E9.5, E10.5 and E11.5 mouse lung single cells. (**B**) Heatmap of top 30 genes of endoderm and mesoderm cells. (**C**) Cell type markers were visualized by UMAP.

**Figure S7. Clustering of E12.5 mouse lung single cells.** (**A**) Cell type definition of E12.5 mouse lung single cell. (**B**) Heatmap of top 15 gene of E12.5 mouse lung cell type. (**C**) Visualization of cell type markers by UMAP. (**D**) Clustering of mesenchymal cells and (**E**) purity of mesenchymal cells. (**F**) Visualization of specific genes of each cluster by dot plots. Mesen, mesenchymal cell, Epi, epithelial cell, Immu, immune cell, Endo, endothelial cell, Unkn, unknown cell type.

**Figure S8. Clustering of E14.5 mouse lung single cells.** (**A**) Cell type definition of E14.5 mouse lung single cell. (**B**) Heatmap of top 15 gene of E14.5 mouse lung cell type. (**C**) Visualization of cell type markers by UMAP. (**D**) Clustering of mesenchymal cells and (**E**) purity of mesenchymal cells. (**F**) Visualization of specific genes of each cluster by dot plots. Mesen, mesenchymal cell, Epi, epithelial cell, Immu, immune cell, Eryth, erythroid cell.

**Figure S9. scRNA-seq on α-SMA-GFP^+^, Tbx4-lineage fibroblasts from E16.5 mouse lung.** (**A**) Flow sorting for α-SMA-GFP^+^, Tbx4-lineage fibroblasts. Violin plot showing nFeature_RNA and percent.mt detected in each cell before (**B**) and after QC (**C**). (**D**) UMAP visualization of lung cell types. (**E**) UMAP visualization of cell type markers. Clustering (**F**) and purity (**G**) of mesenchymal cells. (**H**) Visualization of cluster specific genes by dot plots.

**Figure S10. scRNA-sequencing on P1 mouse lung.** (**A**) Data integration of two batches of scRNA-seq data. (**B**) Cell type definition of P1 lung single cells. (**C**) Heatmap of top 15 differential expression genes comparing each cell types. (**D**) UMAP visualization of relative expression of cell type specific markers. (**E**) UMAP visualization of relative expression of known cell type specific markers to validate the purity of fibroblasts. (**F**) Visualization of specific genes of each cluster by dot plots. Mesen, mesenchymal cell, Epi, epithelial cell, Immu, immune cell, Endo, endothelial cell.

**Figure S11. Clustering of P7 mouse lung fibroblasts.** (**A**) UMAP visualization of clustering of the Pdgfra-GFP fibroblasts from P7 mouse lung. (**B**) UMAP visualization of relative expression of known cell type specific markers to confirm the purity of fibroblasts. (**C**)Visualization of mesenchymal subpopulation specific gene expression in P7 mouse lung mesenchymal cells by dot plots.

**Figure S12. Clustering Pdgfra-GFP cells of P15 mouse lung.** (A-B) UMAP visualization of clustering and cell type clustering in P15 mouse lung single cell. (C) UMAP visualization of relative expression of known cell type specific markers. Clustering (D) and purity (E) of mesenchymal cells in P15 mouse lung. (F) Specific genes of each mesenchymal cell clusters visualized by dot plots. Mesen, mesenchymal cell, Immu, immune cell, Endo, endothelial cell.

**Figure S13. Identification of adult mouse lung mesenchymal cells.** UMAP visualization of sample integration from published data by Aran (**A**), Raredon (**C**) and Reyfman (**E**). (**C**) UMAP visualization of mesenchymal cell identification from published data by Aran (**B**), Raredon (**D**), Reyfman (**F**), Parimon (**G**) and Xie (**H**). Mesenchymal cells were circled by dotted lines. Mesen, mesenchymal cell, Immu, immune cell, Epi, epithelial cell, Endo, endothelial cell.

**Figure S14. Clustering of adult mouse lung fibroblasts.** UMAP visualization of mesenchymal cell integration (**A**) and clustering (**B**) of the fibroblasts from adult mouse lung. (**C**) UMAP visualization of relative expression of known cell type specific markers to validate the purity of fibroblasts. (**D**) Specific genes of each mesenchymal cell cluster visualized by dot plots.

**Figure S15. Clustering of fibrosis mouse lung fibroblasts.** Cell types of fibrosis mouse lung from published papers were defined (**A**-**B**). Mesenchymal cells were circled by dotted lines. (**C**) UMAP visualization of mesenchymal cell integration and clustering of the fibroblasts from fibrosis mouse lung. (**D**) UMAP visualization of relative expression of known cell type specific markers to validate the purity of fibroblasts. (**E**) Specific genes of each mesenchymal cell cluster visualized by dot plots. Mesen, mesenchymal cell, Immu, immune cell, Epi, epithelial cell, Endo, endothelial cell.

**Figure S16. scRNA-seq on aged mouse lung.** Violin plot showing nFeature_RNA, nCount_RNA and percent.mt detected in each cell of each sample before (**A**-**C**) and after QC (**D**-**F**). UMAP visualization of mesenchymal cell integration (**G**) and cell types (**H**) of the single cells from aged mouse lung. (**I**) UMAP visualization of relative expression of known cell type specific markers.

**Figure S17. Clustering of aged lung mesenchymal cells.** (**A**) UMAP visualization of mesenchymal cell clustering from aged mouse lung. (**B**) UMAP visualization of relative expression of known cell type specific markers to validate the purity of fibroblasts. (**C**) Specific genes of each mesenchymal cell cluster visualized by dot plot.

**Figure S18. scRNA-seq on aged fibrosis mouse lung.** Violin plot showing nFeature_RNA, nCount_RNA and percent.mt detected in each cell of each sample before (**A** -**C**) and after QC (**D** -**F**). UMAP visualization of mesenchymal cell integration (**G**) and cell types (**H**) of the single cells from fibrosis mouse lung. (**I**) UMAP visualization of relative expression of known cell type specific markers.

**Figure S19. Clustering of aged fibrosis lung mesenchymal cells.** (**A**) UMAP visualization of mesenchymal cell clustering from aged fibrosis mouse lung. (**B**) UMAP visualization of relative expression of known cell type specific markers to confirm the purity of fibroblasts. (**C**) Specific genes of each mesenchymal cell cluster visualized by dot plot.

**Figure S20. Visualization of novel lipofibroblast marker expression in mouse lung lipofibroblasts of different timepoints.** (**A**) Visualization of differentially expressed genes of lipofibroblasts at each time point by Volcano plots. Genes in red, p-value < 10^−5^, fold-change (logFC) > 1, genes in black, p-value < 10^−5^, logFC < 1, genes in grey, p-value > 10^−5^, logFC > 1. (**B**) Visualization of Gyg, Macf1, Wnt2 and Col13a1 expression in mouse lung mesenchymal cells of different timepoints.

**Figure S21. Cd249^+^ fibroblast FACS and scRNA-seq of lipofibroblast-like cells.** (**A**) Enpep (Cd249) transcript in mouse lung mesenchymal cell. (**B**) FACS gating strategy to obtain Cd249^+^ fibroblasts with concentration match isotype control overlay. (**C**) Mean (±SD) percentage Cd249^+^ fibroblasts vs Cd249^−^ fibroblasts obtained in each mouse lung. **(D**) Heatmap and volcano plot representation of the differentially expressed genes identified in bulk RNA-seq analysis of Cd249^+^ in comparison to Cd249^−^ fibroblasts. (**E**) UMAP visualization and violin plot representation of canonical lipofibroblast marker genes in cultured lipofibroblast-like cells and controls. Wilcoxon, p < 2.2e-16 per comparison. (**F**) UMAP visualization of the top differentially expressed genes identified in the scRNA-seq analysis of the in vivo lipofibroblast cluster in in vitro stimulated lipofibroblasts-like cells.

**Figure S22. Identification of myofibroblasts and SMC markers.** Visualization of known (**A**) and timepoint specific genes for myofibroblasts (**B**) by violin plots in embryonic and postnatal mouse lung mesenchymal cells. UMAP visualization of novel myofibroblast markers in embryonic and postnatal mouse lung mesenchymal cells (**C**-**H**). Visualization of SMC specific markers (**I**) and novel myofibroblasts markers (**J**) in adult and aged normal and fibrosis mouse lung mesenchymal cells.

**Figure S23. SMC specific gene expression in embryonic and postnatal lung mesenchymal cells.** SMC specific gene expression in E9.5-E11.5 (**A**), E12.5 (**B**), E14.5 (**C**), E16.5 (**D**), E17.5 (**E**), P1 (**F**), P7 (**G**) and P15 (**H**) mouse lung mesenchymal cells. Broad or rare expression of these genes indicated no distinct SMC clusters in these mesenchymal cell datasets of these timepoints

**Figure S24. Identification of** *Ebf1***^+^ and pericyte subtypes.** (**A** -**B**) Visualization of timepoint specific genes for *Ebf1*^+^ fibroblasts by violin plots in embryonic and postnatal mouse lung mesenchymal cells. (**C** -**J**) UMAP visualization of *Ebf1*^+^fibroblast specific markers in embryonic and postnatal mouse lung mesenchymal cells. (**K**) violin plot visualization of *Ebf1*^+^ fibroblast and pericyte specific genes in adult and aged normal and fibrosis mouse lung mesenchymal cells.

**Figure S25. ECM related genes expression in normal and fibrosis lung total mesenchymal cells and mesenchymal cell subtypes**. *Acta2* (**A**, **B**, **G** and **H**), *Col1a2* (**C**, **D**, **I** and **J**), and *Col3a1*(**E**, **F**, **K** and **L**) transcripts in total normal and fibrosis lung mesenchymal cell (**A**, **C**, **E**, **G**, **I** and **K**) and mesenchymal cell subtypes (**B**, **D**, **F**, **H**, **J** and **L**) of adult (**A** -**F**) and aged (**G** -**L**) mouse were visualized by violin plots.

**Figure S26. Expression of fibrotic related gene and novel myofibroblast genes.** Transcript of known fibrotic related genes, *Col1a1*, *Col1a2*, *Acta2*, *Fn1* and myofibroblast specific genes, *Enpp2*, *Hhip*, *Tgfbi*, *Wnt5a* in adult (**A**), fibrosis (**B**), aged (**C**) and aged fibrosis (**D**) mesenchymal cells.

**Figure S27. Transcript of Col14a1 in lung mesenchymal cells and Vim in lung total cells of different timepoints**. (**A**) In embryonic stages, Col14a1 was mainly expressed in pre-lipofibroblasts/lipofibroblasts, however, after birth, Col14a1 expression switched to Ebf+ fibroblasts and in adult and aged lungs Col14a1 kept its transcripts in *Ebf1*^+^ fibroblasts. (**B**) In early embryonic stages (E9.5-E11.5), Vim showed transcripts in both mesoderm and endoderm cells. In later embryonic, postnatal, adult and fibrosis stages, Vim showed highest transcripts in endothelial cells, showed weakened transcripts in immune cells and mesenchymal cells and was rarely detectable in epithelial cells. Endo, endoderm, Meso, mesoderm, Mesen, mesenchymal cells, Immu, immune cells, Epi, epithelial cells, Endo, endothelial cells, Unkn, unknown.

**Figure S28. Expression of *Pdgfra* and *Pdgfrb* in mouse lung mesenchymal cells of different timepoints**. (**A**) *Pdgfra* transcripts in mesenchymal cells of different time points. Pdgfra expression was mainly in myofibroblasts in embryonic and postnatal lung mesenchymal cells and switched to lipofibroblasts in adult and aged normal and fibrosis mesenchymal cells with high background in other subtypes. (**B**) *Pdgfrb* showed higher expression in *Ebf1*^+^ fibroblasts and pericytes with some background in other subtypes.

**Figure S29. Co-expression of genes in normal and fibrosis lung mesenchymal cells.** Visualization of blend expression of *Acta2* and *Pdgfra* (**A** and **E**), *Pdgfrb* and *Ebf1* (**B** and **F**), *Pdgfrb* and *Pdgfra* (**C** and **G**), *Tcf21* and *Pdgfra* (**D** and **H**) in normal (**A**-E) and fibrosis (**E** -**H**) mouse lung mesenchymal cells.

**Figure S30. Representative quality control, sub-setting and integration of human scRNA-seq data.** Violin plot showing number of genes (nFeature_RNA), number of read counts (nCount_RNA) and the percentage of transcripts mapping to mitochondrial genes (percent.MT) detected in each cell before (**A**, **C**) and after (**B**, **D**) QC. (**E**) Dot plot representation of common marker genes used to identify and subset mesenchymal from epithelial, endothelial and immune cells in each dataset. Dot size corresponds to percentage of cells in a cluster expressing the gene and color to expression level. (**F**) UMAP visualization to the integration of scRNA-seq data sets of healthy and IPF donor lung mesenchymal cells. Mesen, Mesenchymal, Epi, Epithelial, Immu, Immune, Endo, Endothelial.

