## Supplemental information for "Definition and Signatures of Lung Fibroblast Populations in Development and Fibrosis in Mice and Men"

### Supplementary Methods

#### Bleomycin instillation

Bleomycin instillation were described previously[1]. Briefly, under anesthesia the trachea was surgically exposed. 1.25U/kg bleomycin (Hospira, Lake Forest, IL) in 25 µl PBS was instilled into the mouse trachea with a 25-G needle inserted between the cartilaginous rings of the trachea. Control animals received saline alone. The tracheostomy site was sutured, and the animals monitored intensively until active. Animals were randomly allocated to control or treatment groups. Bleomycin treated mice were actively monitored by trained animal welfare staff until sacrificed. Mice were sacrificed at indicated and lung tissues were collected.

#### *In vitro* culture of murine lung fibroblasts

Murine lungs were dissociated as described above. The entire cell suspension was then plated on appropriately sized tissue culture plasticware. Fibroblasts were cultured in advanced DMEM/F12 (#12634010, Thermo Fisher Scientific), with 10 % vol/vol FBS (HyClone) or to induce a lipofibroblast like phenotype with the addition of 1 % vol/vol ITS+3 liquid media supplement (#I2771, Merck Milipore, Burlington, MA, USA) to the culture media. Media was changed every other day and the cells sub cultured when 80 % confluent. Fibroblasts cultured to passage three (P3) were considered free of contaminating cells based on previous experience in the laboratory. At P3 the cultured fibroblasts were then stimulated with one or a combination (as indicated) of 10 µM rosiglitazone (#72622, Stem Cell Technologies, Cambridge, MA, USA), 1 µM SB4315442 (#1614, R&D Systems, Minneapolis, MO, USA), 4 µM, rhBMP4 (#314-BP, R&D Systems) for

14 days. Matched control cells isolated from the same lung were stimulated in an identical media, for an identical duration, with the appropriate vehicle.

***In vitro* 3D organoid culture with cultured lipofibroblast like cells**

Fibroblasts in which a lipofibroblast like phenotype had been induced *in vitro,* and their associated control cells, were cultured in Matrigel/Medium (1:1) mixture in the presence of flow sorted type 2 epithelial cells ($Cd45^-$ $Cd31^-$ $Cd24^-$ $Cd34^-$ $Sca1^-$ $CD326^+$, Supplementary Table s1). 100 $\mu$l Matrigel/medium mix containing 3 x $10^3$ AEC2 cells and 2 x $10^5$ control or lipofibroblast like cells were plated into each 24 well 0.4 4 $\mu$m Transwell insert. 400 $\mu$l of medium was added in the lower chambers. Medium was described previously[2]. Matrigel was from Corning (#354230, Corning, One Riverfront Plaza, NY, USA). Half of the media in each well was changed every other day. Cultures were maintained in humidified 37°C and 5% $CO_2$ incubator. Colonies were visualized with a Zeiss Axiovert40 inverted fluorescent microscope (Carl Zeiss AG, Oberkochen, Germany). Number of colonies with a diameter of ≥50 µm from each insert was counted and colony-forming efficiency (CFE) was determined by the number of colonies in each culture as a percentage of input epithelial cells at day 14 after plating.

**Fluorescence activated cell sorting (FACS)**

For staining cells were counted and diluted to a maximum of $10^6$ cells/100 µl in $HBSS^+$. The cell suspension was then divided into Eppendorf's for staining. Controls included unstained, single color controls and where appropriate isotype controls. The cells were pelleted in a pre-cooled (4°C) centrifuge at 1600 rpm for 5 minutes and resuspended in appropriately diluted primary antibody. The cell suspension was incubated with the antibody and live/dead maker (fixable viability) on ice in the dark for between 30 minutes to 1 hour. The cells were then washed by gentle pipetting with $HBSS^+$. The cell suspension was centrifuged, the supernatant carefully removed, and the cells washed again. In total three wash steps were performed after each incubation. If the antibody was directly conjugated following the final wash the cells were resuspended in 500 µl $HBSS^+$ and passed through a cell strainer cap into a falcon test tube by centrifugation prior to FACS. If the antibody was not directly conjugated following the final wash the cells were incubated with appropriate secondary antibody for 30 minutes to 1 hour on ice protected from light. The cells were then washed as described previously and resuspended prior to analysis. The antibodies were used at optimized concentrations and are listed in Supplementary Methods Table 1. Where DAPI was used as the live/dead marker it was added to the cell suspension prior 5-10 minutes prior to sorting. Total mesenchymal cells were analyzed using Becton Dickinson Fortessa or sorted using a 13-color Becton Dickinson FACS Aria III (BD Biosciences, San Jose, CA) from both mice and human single cell lung suspensions using negative selection as: live cells (fixable viability or DAPI negative) and EPCAM/PECAM1/PTPRC negative.

**RNA-seq analysis**

Total RNA was extracted from $CD249^+$ and $CD249^-$ fibroblasts flow sorted from murine lungs using the RNeasy Micro Kit (Qiagen, Valencia, CA, USA) according to manufacturer's instructions. Total RNA was stored at -80°C until the day of analysis. RNA integrity of an aliquot from each sample was analyzed using a Bioanalyzer (Agilent Technologies, Santa Clara, CA, USA) and only samples with a RIN ≥8 retained for analysis. A minimum of 50 ng and maximum of 400 ng total RNA in a maximum of 20 $\mu$l was sequenced (1x75 bp single-end sequencing, average 25 million reads/sample) using a NextSeq 500 (Illumia, San Diego, CA, USA). Library preparation, library QC and differential expression analysis was performed by the Cedars-Sinai Genomics Core facility.

**Histology and Immunofluorescence staining**

To prepare the mouse lungs for histology; after being deeply anaesthetized and sacrificed as described previously, the trachea was cannulated. The left lung cleared of blood by perfusion of the pulmonary artery with PBS via cardiac puncture. The lungs were then inflated with 0.5ml pf 10% neutral buffered formalin. The tissues were fixed overnight, and the following day embedded in Optimal Cutting Temperature Compound (OCT, Sakura Finetek USA, Torrance, CA, USA) and flash frozen. Cryosections (5 $\mu$m) were cut using a cryostat (Lecia, Buffalo Grove, IL, USA) onto Superfrost Plus Microscope Slides (Fisherbrand, Fisher Scientific, Waltham, MA, USA). Immunofluorescence was performed using primary antibodies raised against the following antigens and used at the indicated dilutions to stain slides overnight at 4°C: α-Smooth Muscle - Cy3™, (#C6198, mouse monoclonal, clone 1A4, Sigma-Aldrich, clone 1A4), Ebf1, (#AF5165, polyclonal goat IgG, R&D Systems, Minneapolis, MN, USA), Von Willebrand Factor, VWF (,#ab6994, polyclonal rabbit IgG, Abcam, Eugene, OR, USA).

To stain intracellular lipid droplets in lipofibroblast like cells and controls the media was removed and the cells washed with PBS. The cells incubated with 10 $\mu$M Bodipy 493/505 (#D3922, Invitrogen, Carlsbad, CA, USA) or 1 mg/ml Nile Red (N1142, Invitrogen), protected from light in a tissue culture incubator at 37°C. The Bodipy solution was removed and the cells washed twice with PBS. The cells were fixed *in situ* with 4 % vol/vol formaldehyde at room temperature for 15 minutes protected from light and stained with 10 $\mu$g/ml DAPI for 10 minutes prior to imaging. For Oil Red O (#O0625, Merck Millipore Burlington, MA, USA) staining after fixation the cells were dehydrated with 100 % 1, 2-Propanediol solution for 5 m minutes. This step was repeated and then 2 ml/ 10 $cm^2$ 0.5 % Oil Red O solution diluted in 1, 2-Propanediol solution was added and incubated for 30 minutes at 37°C. The stain was aspirated and differentiated with 85 % 1, 2-Propanediol solution for 1 minute. The cells were then rinsed with $dH_2O$ 2-3 times, counterstained with Mayer's Hematoxylin for 10-15 minutes at room temperature. The cells were rinsed 4-5 times with $dH_2O$ and then imaged. Stained sections were imaged using Zeiss 780 reverse Laser Scanning Confocal Microscope.

### Bioinformatics Methods

#### Read alignments

The demultiplexed raw reads were aligned to the transcriptome using STAR (version 2.5.1) with default parameters, using human GRCh38 (or mouse mm10) transcriptome reference from Ensembl version 84 annotation, containing all protein coding and long non-coding RNA genes. Expression counts for each gene in all samples were collapsed and normalized to unique molecular identifier (UMI) counts using Cell Ranger software version 3.0 (10X Genomics). The result was a large digital expression matrix with cell barcodes as rows and gene identities as columns.

#### Quality control, cell clustering, doublet calling and annotation

Expression profiles of cells from different subjects and publicly available datasets were analyzed and clustered separately using the R software package Seurat (version 3.0)[3]. For each individual sample the total number of genes detected per cell ('nFeature_Gene'), number of transcripts per cell ('nFeature_RNA') and percentage of transcripts mapping to mitochondrial genes ('percent.mt' or ‘percent.MT’) were visualized. Samples with less than 5 cells and/or less than 200 detected genes per cell were excluded from further analysis. Quality control based on these metrics included to the exclusion of outliers, low quality cells with low gene and/or transcripts detected per cell and/or cells with a high number of transcripts mapping to mitochondrial genes. Doublets were identified as outliers with a dramatically higher number of detected genes per cell than the median ±interquartile range genes detected per cell. After QC unique molecular identifiers (UMIs, 10x) were then normalized across cells, scaled per $10^4$ and converted to log scale using the ‘NormalizeData’ function. These data were converted to z-scores using the ‘ScaleData’ command and highly variable genes were selected with the ‘FIndVariableGenes’ function. To integrate multiple samples integration anchors were identified in the list of samples (individual Seruat Objects) with the ‘FindIntegrationAnchors’ command and the list of samples integrated with the ‘IntegrateData’ function. Principal components were calculated for these selected genes with the ‘RunPCA’. The optimum dimensionality of the dataset for downstream clustering was determined using both the JackStraw and Elbow plot methods. Clusters of similar cells were detected using the Louvain method for community detection including only biologically meaningful principle components to construct the shared nearest neighbor map and an empirically set resolution, as implemented in the ‘FindClusters’ and ‘FindNeighbors’ functions.

Clusters were assigned an identity to a given cluster based on expression of tissue compartment markers (e.g. *EPCAM, CDH1, PECAM1, EMCN, PTPRC, CD86, COL1A1, COL1A2, WT1).* The mesenchymal fractions were identified by expression of known marker genes commonly reported in the literature and separated for further analysis using the ‘SubsetData’ command. RNA markers for each cluster were identified using the ‘FindAllMarkers’ command in Seurat and examining the top differentially expressed genes in each cluster for homology with the known marker genes. Where a cluster could not be identified using known marker genes, they were identified by a highly discriminative gene that was among the most differentially expressed in that cluster. Differentially expressed genes for each mesenchymal subpopulation relative to all other mesenchymal cells were identified using the ‘MAST’ statistical framework implemented in the ‘FindMarkers’ command[4]. To obtain the most sensitive and specific differentially expressed genes for each subpopulation we identified genes with a p-value less than $10^{-5}$ and an average log fold-change greater than 1.

Comparative analysis of changes in gene expression between mesenchymal cells of the same subpopulation from healthy and fibrotic lungs was performed by calculating the log1p (average expression) values for each gene in Seurat and visualizing these on a scatterplot. The genes that changed most significantly between conditions were identified and annotated using the 'MAST' statistical framework implemented in the 'FindMarkers' command and the top genes annotated on the relevant figure.

Enriched genes were annotated as transcription factors from the differentially expressed genes of each cluster by imputing the top differentially expressed genes into the NCBI, EMBL-EBI, UniprotKB Gene Ontology database. Genes were identified as transcription factors if included under the "DNA binding, transcription factor activity" categorisation in the returned results.

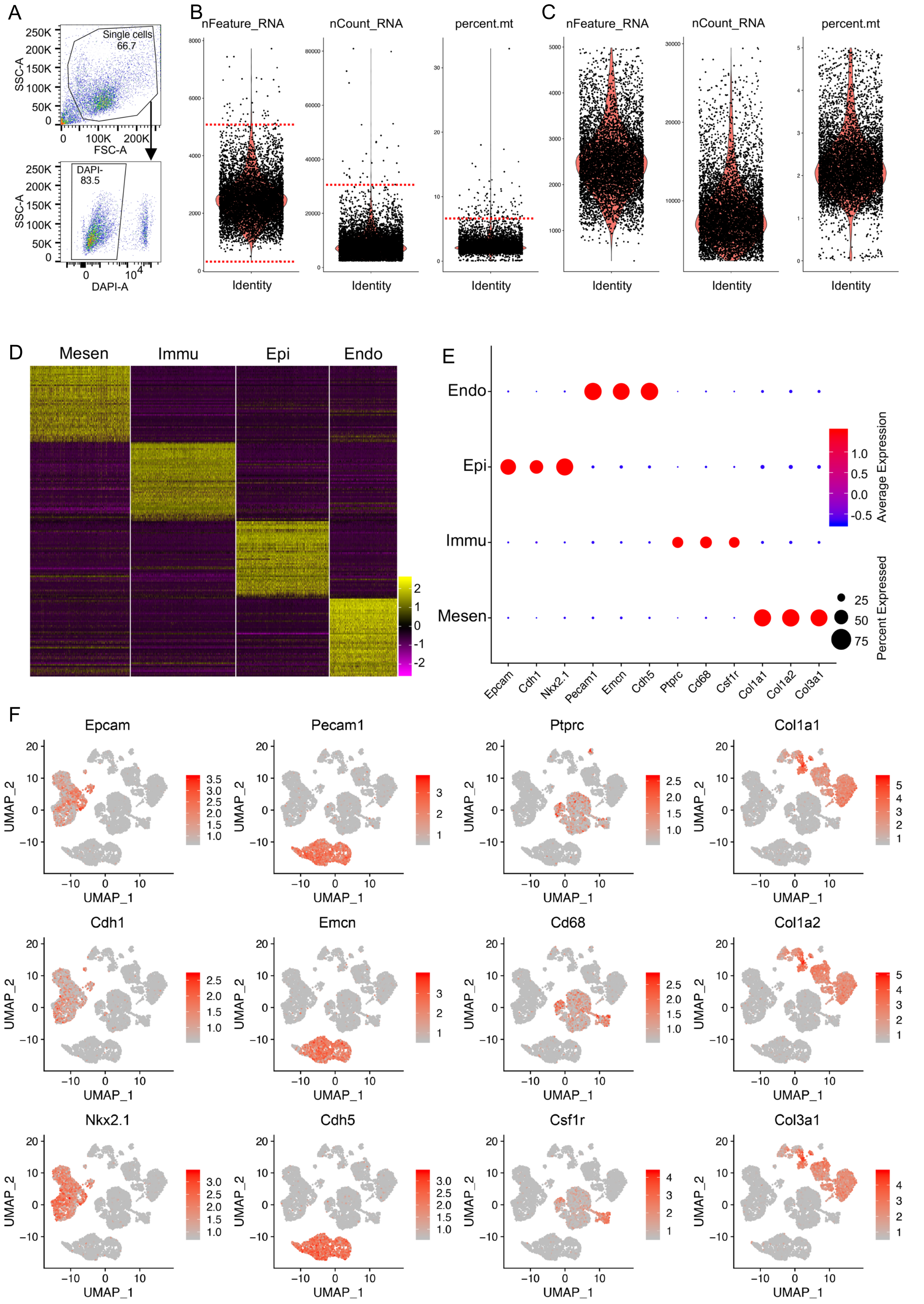


**Figure S1. Single cell RNA-sequencing on E17.5 mouse lung.** Violin plot showing the number of genes (nFeature_RNA), number of read counts (nCount_RNA) and percentage of mitochondria genes (percent.mt) detected in each cell before (**A**) and after QC (**B**). (**C**) Heatmap of top 15 differential expression genes comparing each cell types. (**D**) Dot plot visualization of relative expression of known cell type specific markers of each cell type. (**E**) UMAP visualization of relative expression of known cell type specific markers used for cell type clustering. Mesen, mesenchymal cell, Epi, epithelial cell, Immu, immune cell, Endo, endothelial cell.

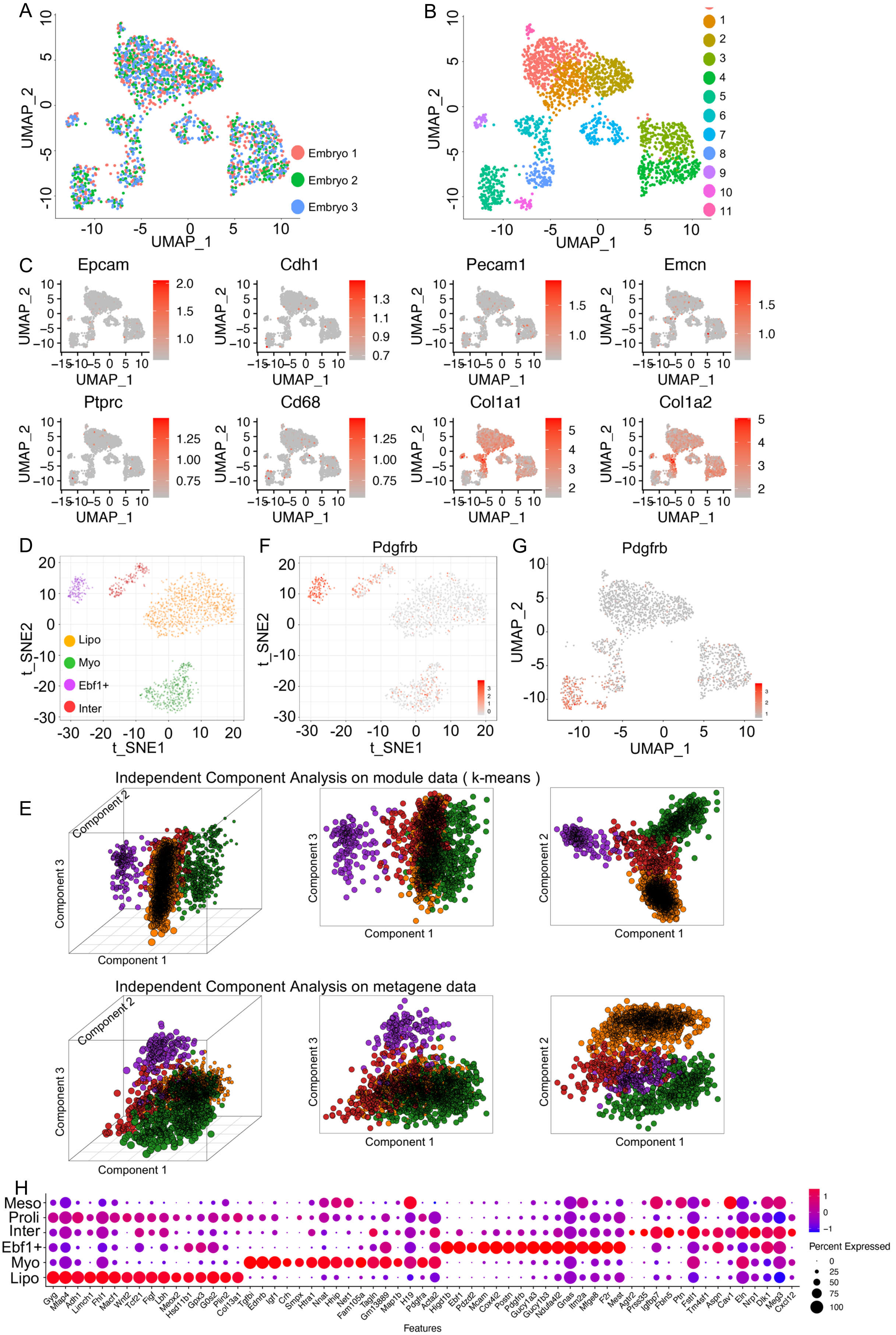


**Figure S2. Clustering of E17.5 mouse lung fibroblasts.** UMAP visualization of mesenchymal cell integration (**A**) and clustering (**B**) of the fibroblasts from E17.5 mouse lung. (**C**) UMAP visualization of relative expression of known cell type specific markers to validate the purity of fibroblasts. Visualization of mesenchymal cell clustering (**D**) by t-SNE from E17.5 mouse lung. (**E**) 3-D and 2-D visualization of PCA analysis on the E17.5 lung mesenchymal cells to validate the clustering. Visualization of Pdgfrb transcript by t-SNE (**F**) and UMAP (**G**) from E17.5 mouse lung. (**H**) Dot plot visualization of specific genes of each subpopulation.

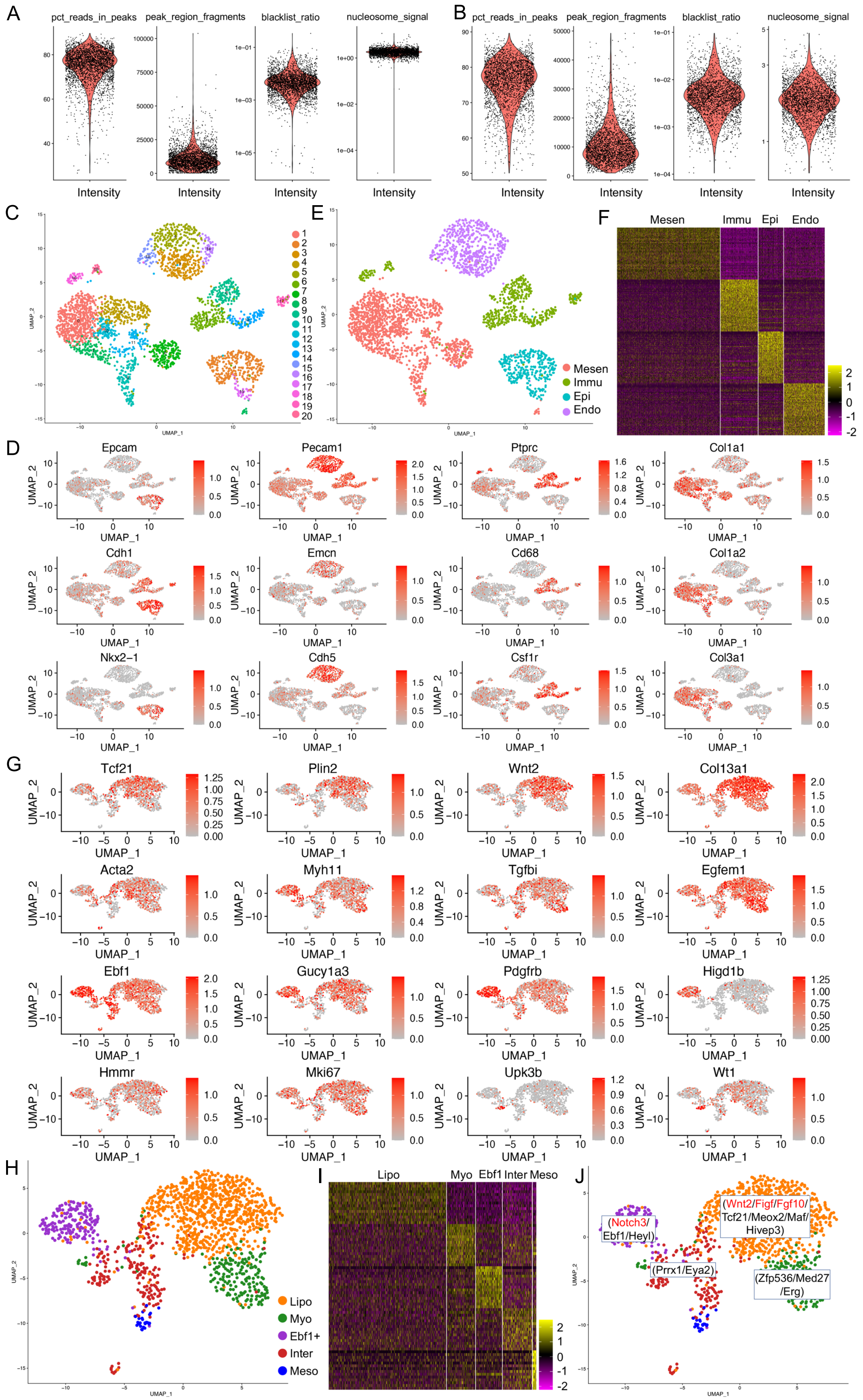


**Figure S3. scATAC-seq on E17.5 mouse lung.** Cell quality of the nuclei before (**A**) and after (**B**) QC. Distribution (**C**), cell type gene accessibilities (**D**) and cell types (**E**) of the nuclei visualized by UMAP. (**F**) Heatmap of top 50 genes of each cell type. (**G**) Cluster specific gene accessibilities were visualized by UMAP. (**H**) Mesenchymal cell sub-clusters were defined. (**I**) Heatmap of top 15 genes of each mesenchymal cell cluster. (**J**) Cluster specific growth factors and transcription factors. Lipo, lipofibroblast, Myo, myofibroblast, *Ebf1*⁺, *Ebf1*⁺ fibroblast, Inter, intermediate fibroblast, Meso, mesothelial cell.

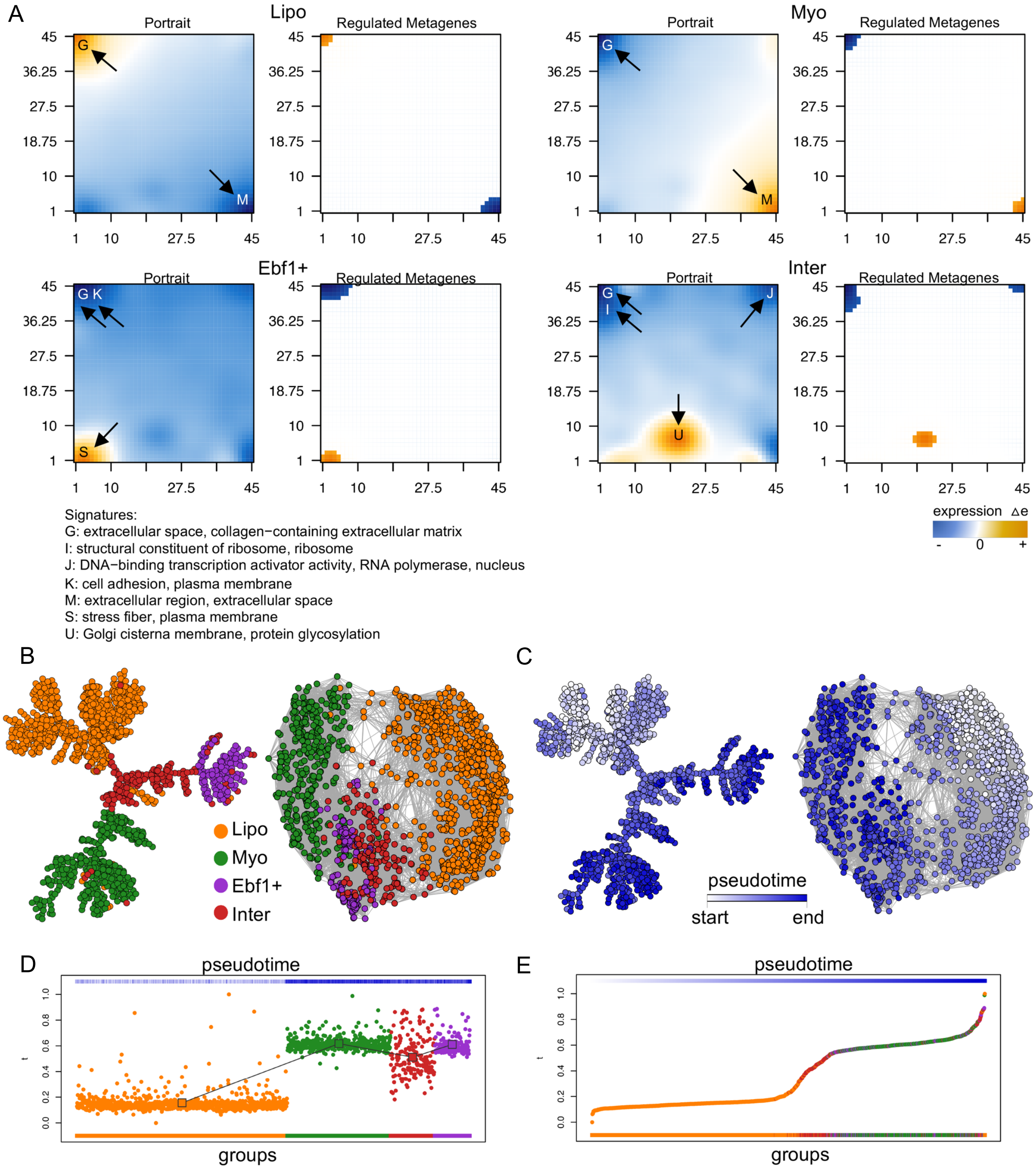


**Figure S4. Differentiation potential of the E17.5 lung fibroblast subtypes.** (**A**) Metagene profile for each subtype in E17.5 mouse lung fibroblasts. Lineage bifurcation of E17.5 mouse lung fibroblast subtypes by Correlation Spanning Tree (**B**-**C**) and group (**D**-**E**) and k−nearest neighbour graph (k=30) (**F**). Lipo, lipofibroblast, Myo, myofibroblast, Ebf1+, Ebf1+ fibroblast, Inter, intermediate fibroblast, Proli, proliferative fibroblast.

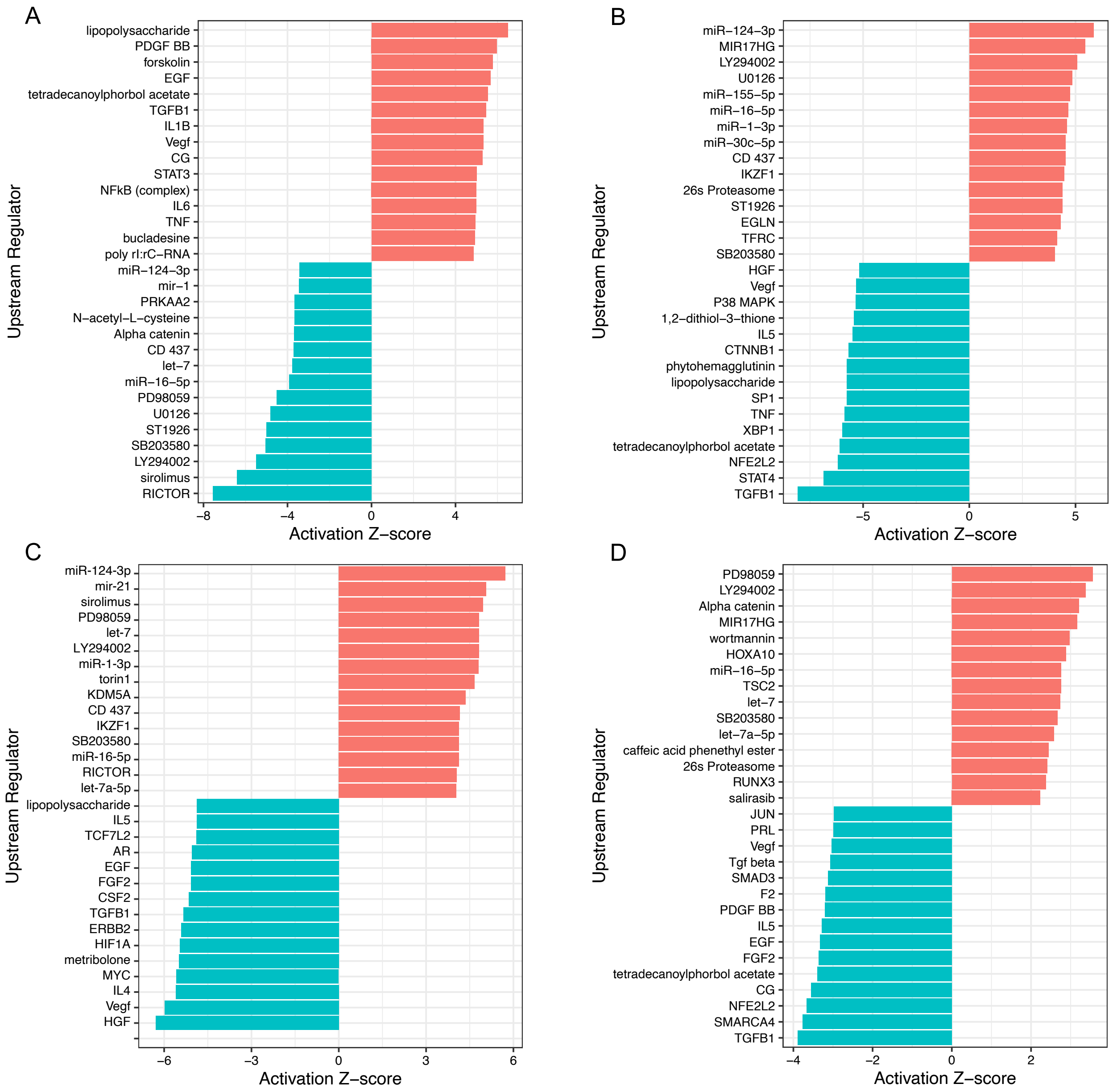


**Figure S5. Visualization of IPA analysis of each clusters of E17.5 fibroblasts.** Top 15 activated and inhibited regulators of lipofibroblast (**A**), myofibroblast (**B**), Ebf1+ fibroblast (**C**) and intermediate fibroblast (**D**) of E17.5 mouse lung.

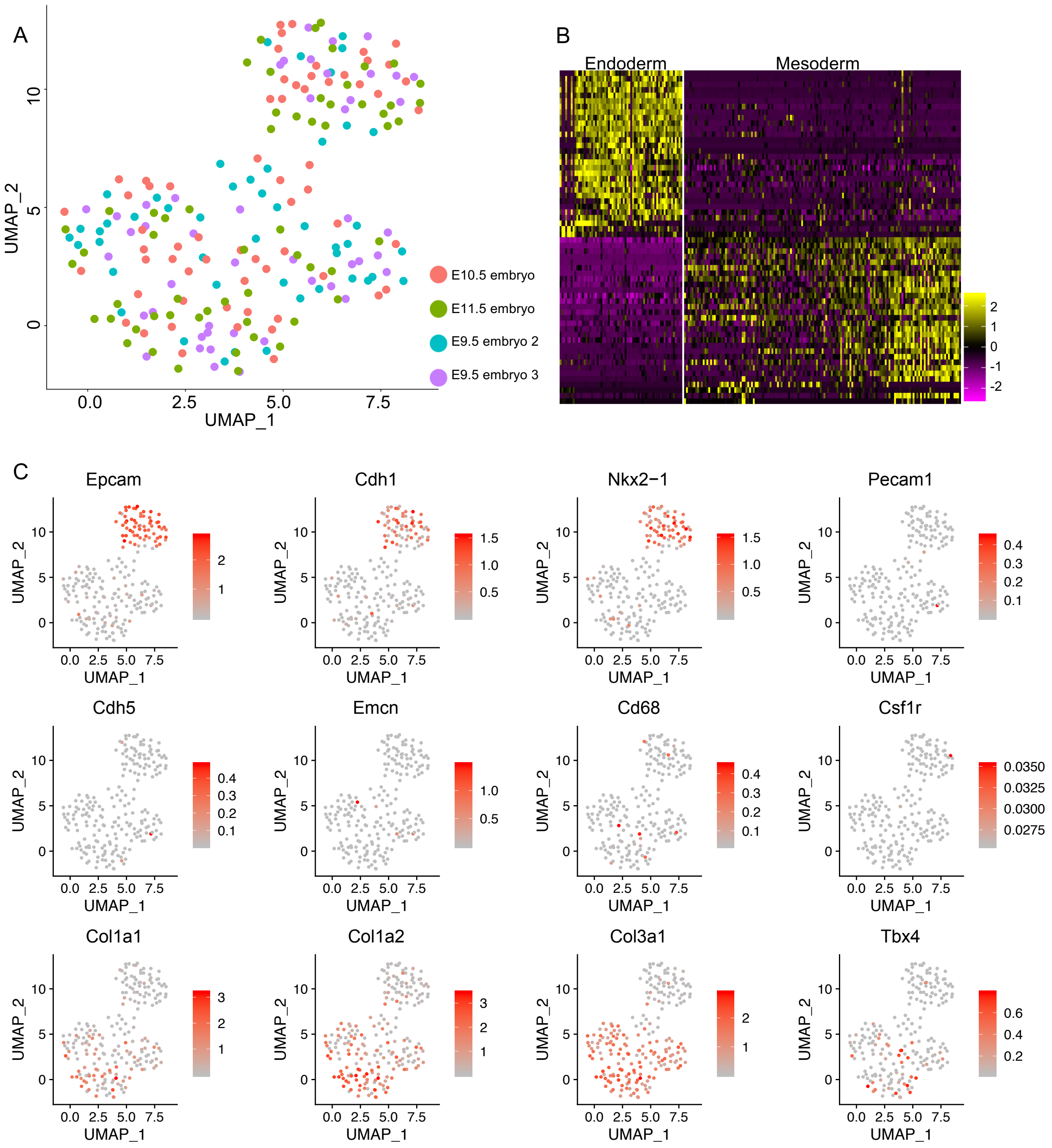


**Figure S6. Clustering of E9.5-E11.5 mouse lung single cells**. (**A**) Cell integration of E9.5, E10.5 and E11.5 mouse lung single cells. (**B**) Heatmap of top 30 genes of endoderm and mesoderm cells. (**C**) Cell type markers were visualized by UMAP.

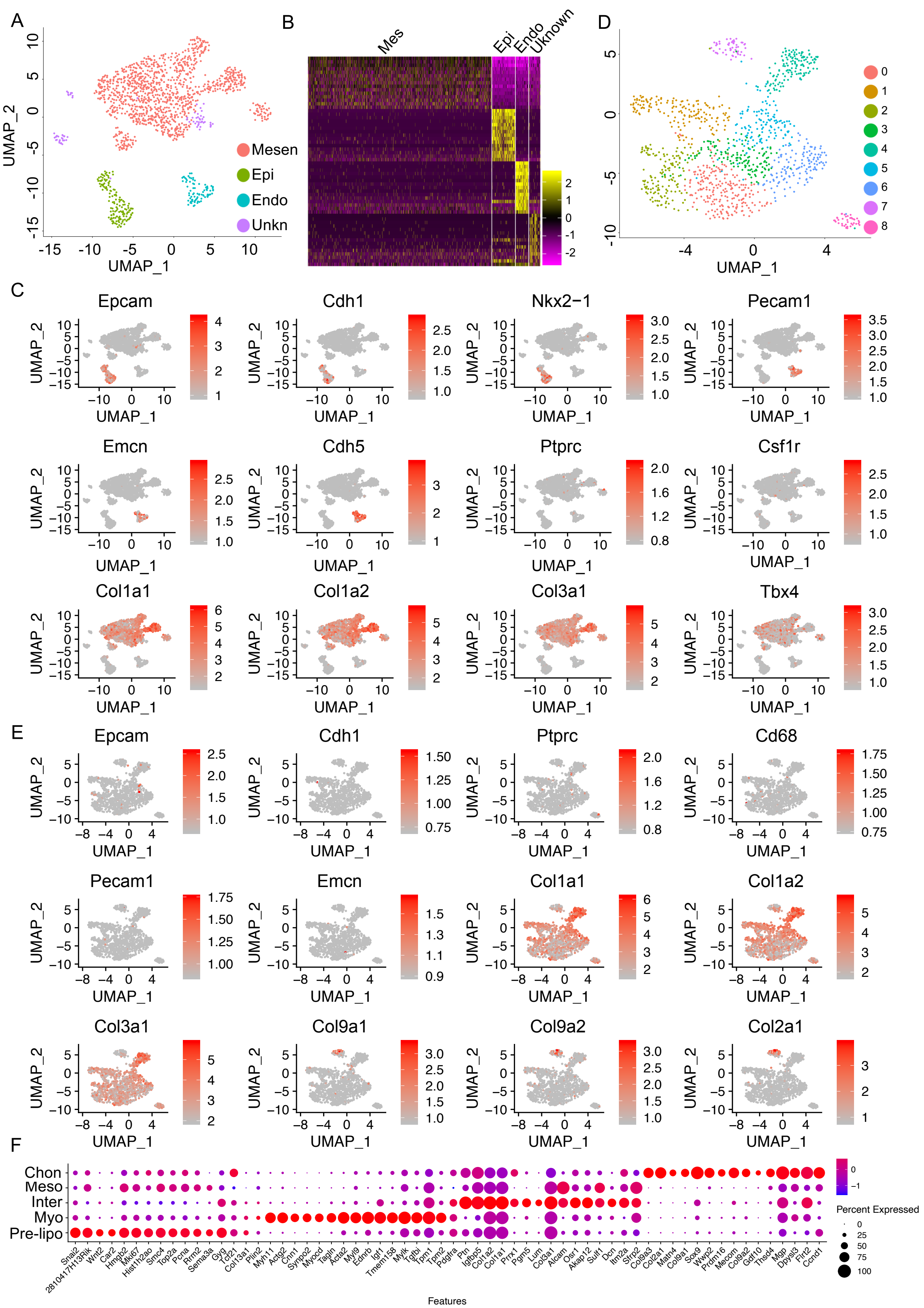


**Figure S7. Clustering of E12.5 mouse lung single cells.** (**A**) Cell type definition of E12.5 mouse lung single cell. (**B**) Heatmap of top 15 gene of E12.5 mouse lung cell type. (**C**) Visualization of cell type markers by UMAP. (**D**) Clustering of mesenchymal cells and (**E**) purity of mesenchymal cells. (**F**) Visualization of specific genes of each cluster by dot plots. Mesen, mesenchymal cell, Epi, epithelial cell, Immu, immune cell, Endo, endothelial cell, Unkn, unknown cell type.

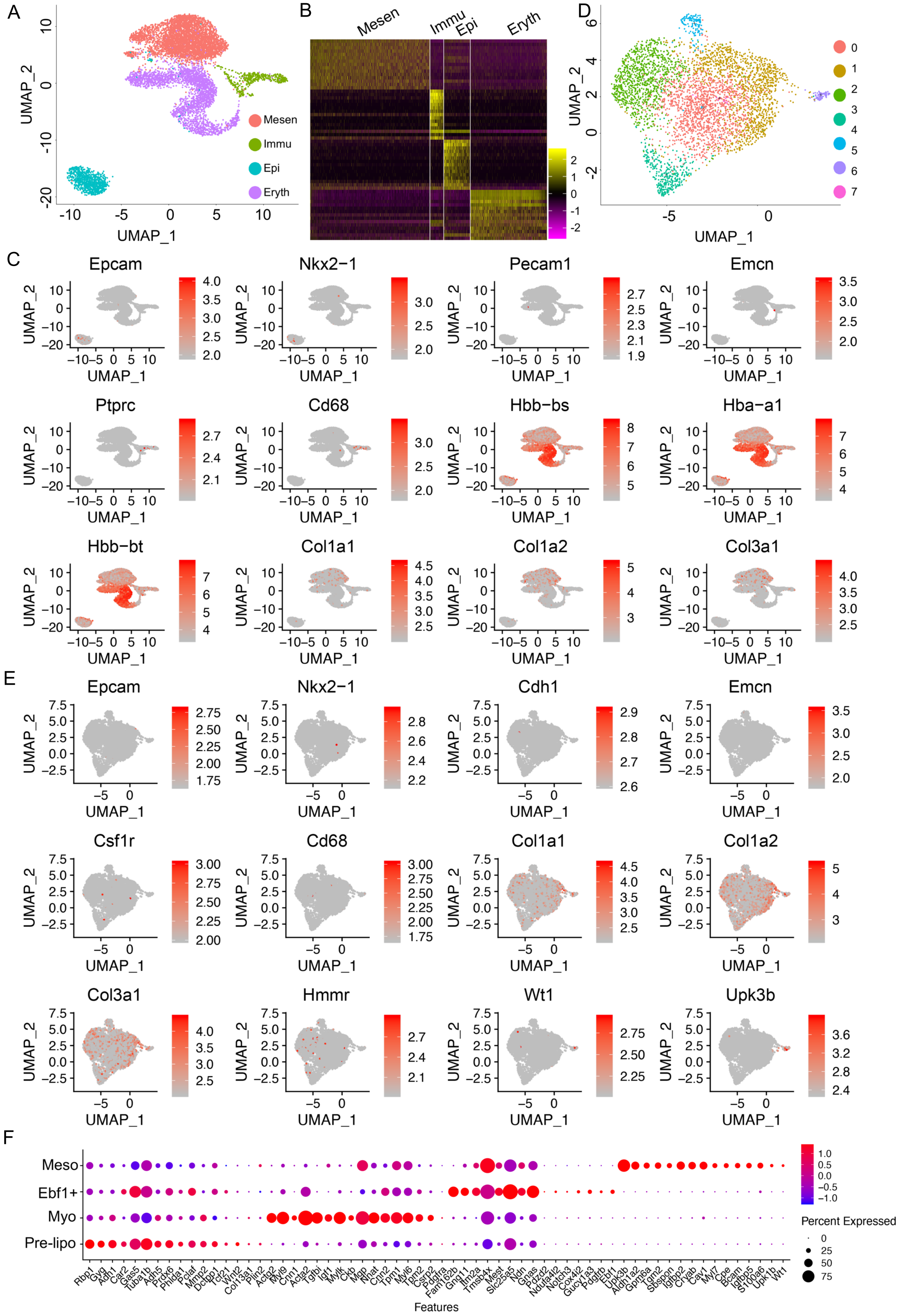


**Figure S8. Clustering of E14.5 mouse lung single cells.** (**A**) Cell type definition of E14.5 mouse lung single cell. (**B**) Heatmap of top 15 gene of E14.5 mouse lung cell type. (**C**) Visualization of cell type markers by UMAP. Clustering of mesenchymal cells (**D**) and purity of mesenchymal cells (**E**). (**F**) Visualization of specific genes of each cluster by dot plots. Mesen, mesenchymal cell, Epi, epithelial cell, Immu, immune cell, Eryth, erythroid cell.

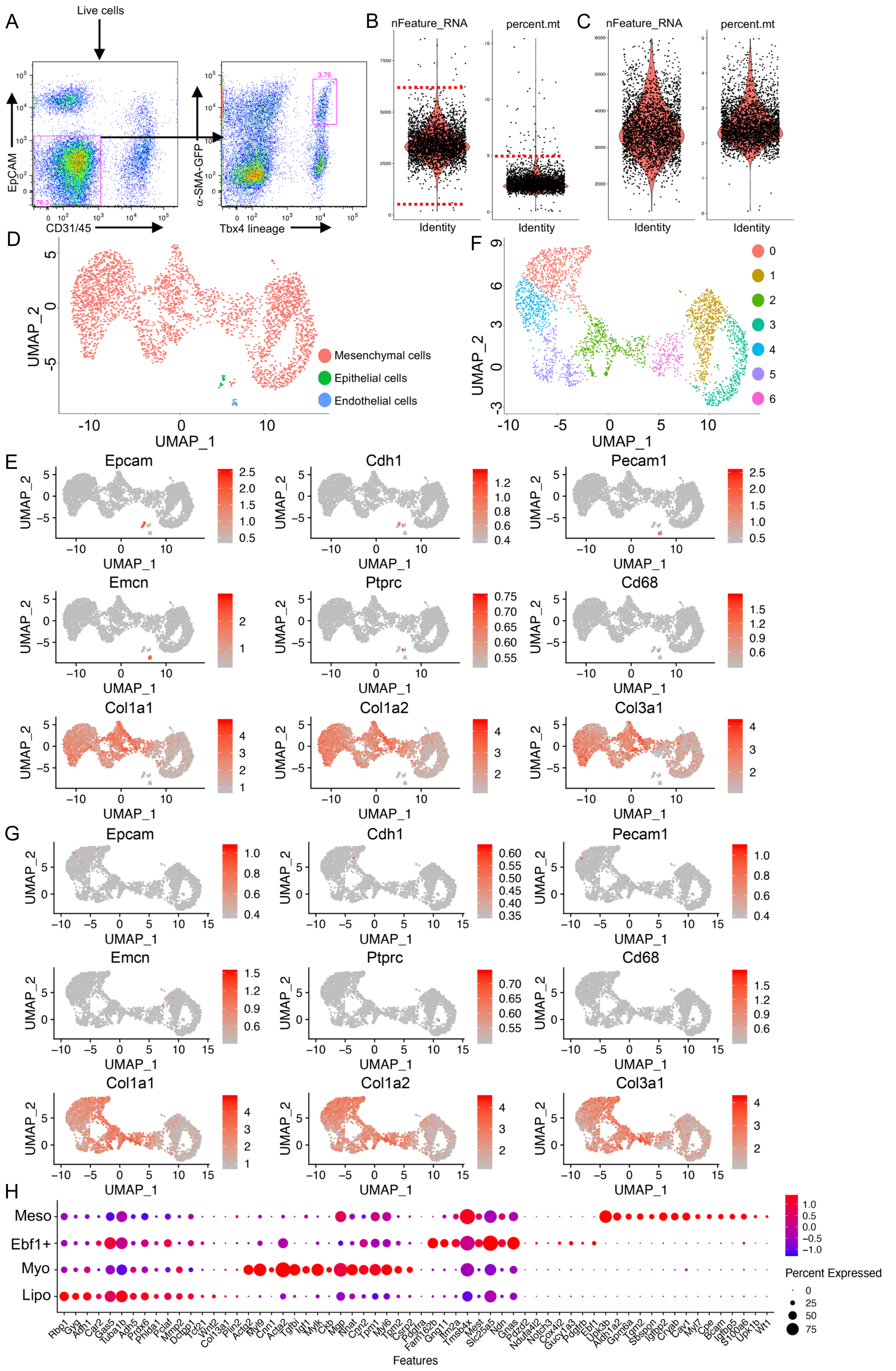

**Figure S9. scRNA-sequencing on α-SMA-GFP+, Tbx4-lineage fibroblasts from E16.5 mouse lung.** (**A**) Flow sorting for α-SMA-GFP+, Tbx4-lineage fibroblasts. Violin plot showing nFeature_RNA and percent.mt detected in each cell before (**B**) and after QC (**C**). (**D**) UMAP visualization of lung cell types. (**E**) UMAP visualization of cell type markers. Clustering (**F**) and purity (**G**) of mesenchymal cells. (**H**) Visualization of cluster specific genes by dot plots.

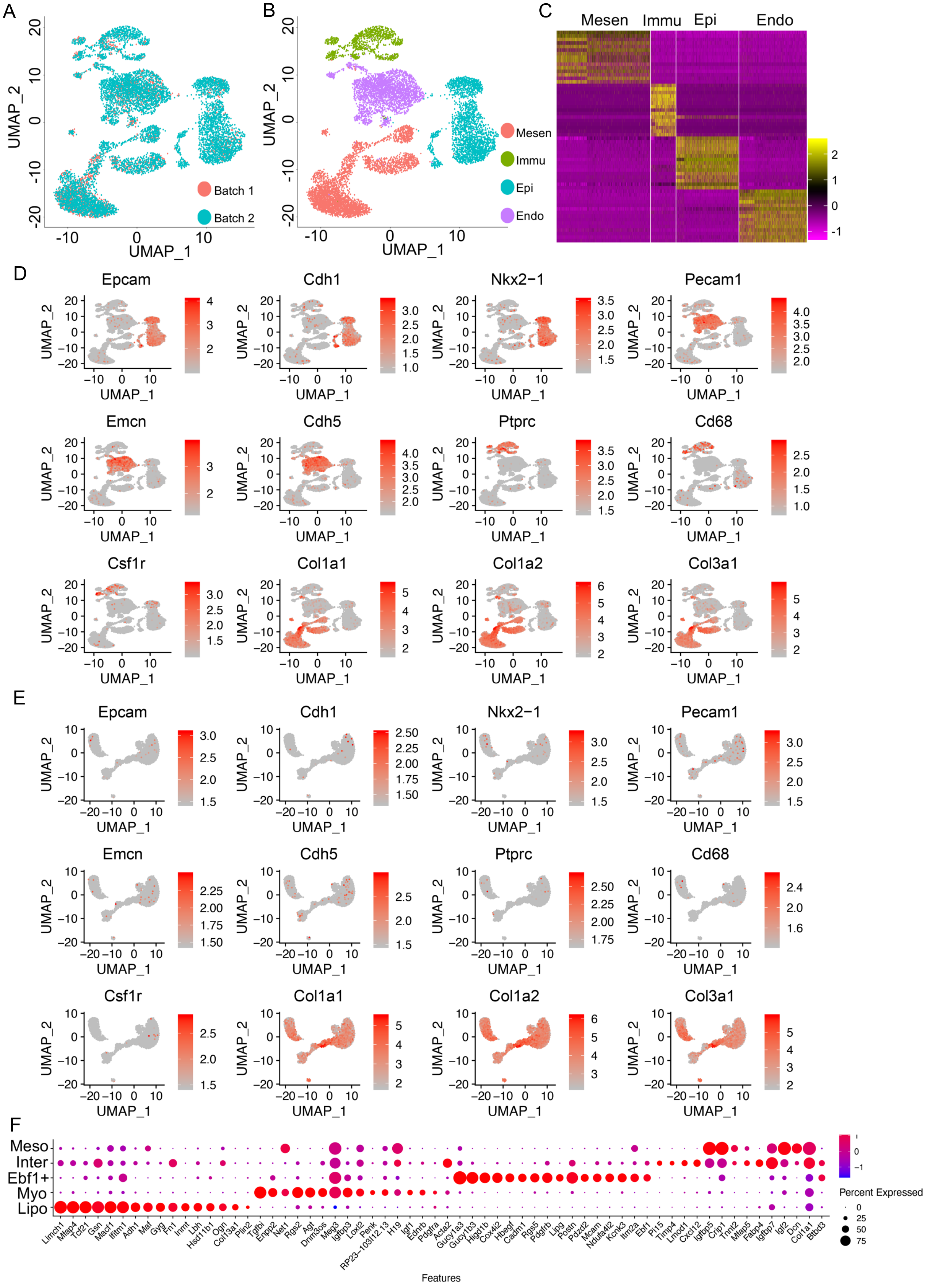


**Figure S10. Single cell RNA-sequencing on P1 mouse lung.** (**A**) Data integration of two batches of single cell RNA-seq data. (**B**) Cell type definition of P1 lung single cells. (**C**) Heatmap of top 15 differential expression genes comparing each cell types. (**D**) UMAP visualization of relative expression of cell type specific markers. (**E**) UMAP visualization of relative expression of known cell type specific markers to validate the purity of fibroblasts. (**F**) Visualization of specific genes of each cluster by dot plots. Mesen, mesenchymal cell, Epi, epithelial cell, Immu, immune cell, Endo, endothelial cell.

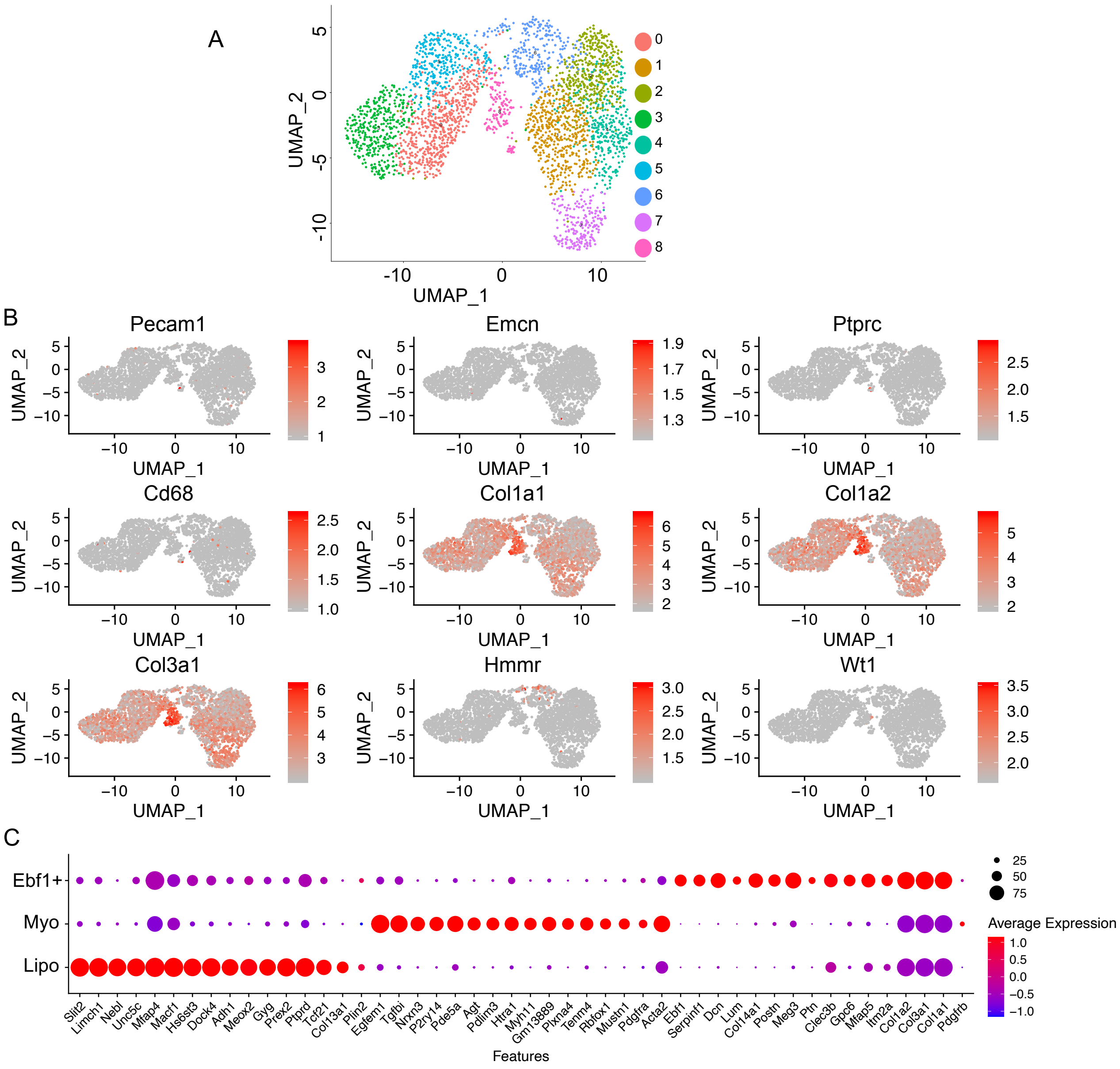


**Figure S11. Clustering of P7 mouse lung fibroblasts.** (**A**) UMAP visualization of clustering of the Pdgfra-GFP fibroblasts from P7 mouse lung. (**B**) UMAP visualization of relative expression of known cell type specific markers to confirm the purity of fibroblasts. (**C**) Visualization of mesenchymal subpopulation specific gene expression in P7 mouse lung mesenchymal cells by dot plots.

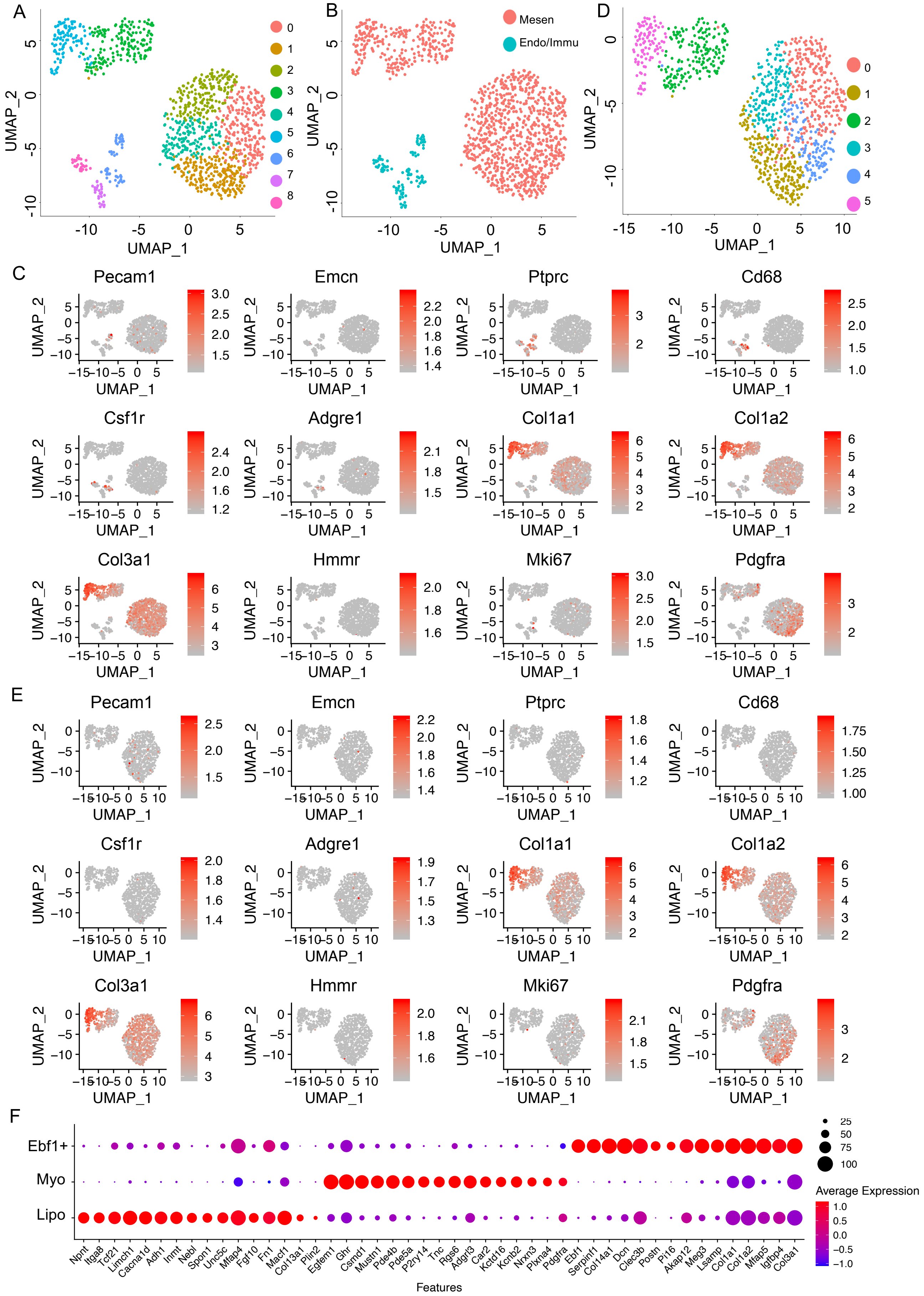


**Figure S12. Clustering Pdgfra-GFP cells of P15 mouse lung.** (**A-B**) UMAP visualization of clustering and cell type clustering in P15 mouse lung single cell. (**C**) UMAP visualization of relative expression of known cell type specific markers. Clustering (**D**) and purity (**E**) of mesenchymal cells in P15 mouse lung. (**F**) Specific genes of each mesenchymal cell clusters visualized by dot plots. Mesen, mesenchymal cell, Immu, immune cell, Endo, endothelial cell.

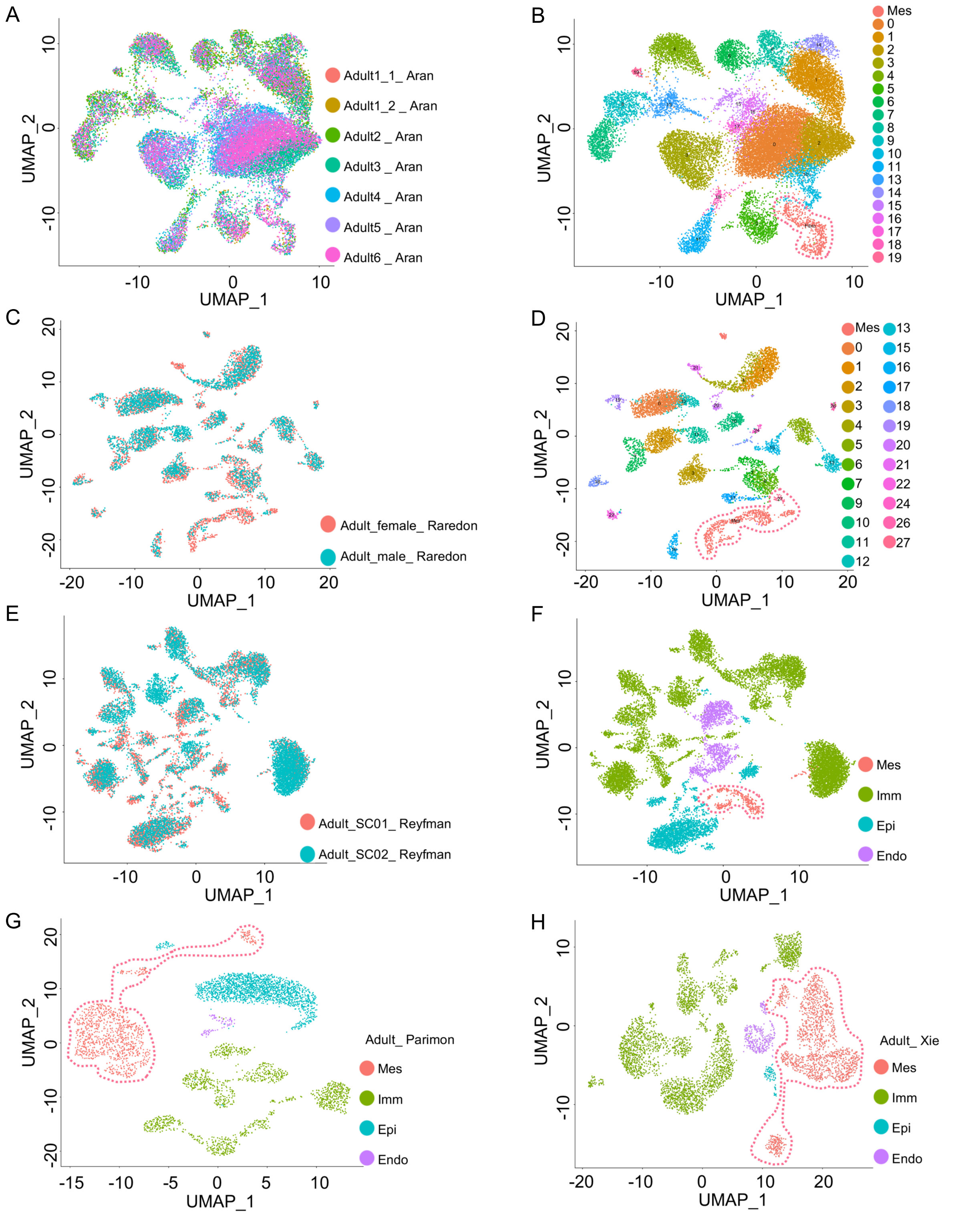


**Figure S13. Identification of adult mouse lung mesenchymal cells.** UMAP visualization of sample integration from published data by Aran (**A**), Raredon (**C**) and Reyfman (**E**). (**C**) UMAP visualization of mesenchymal cell identification from published data by Aran (**B**), Raredon (**D**), Reyfman (**F**), Parimon (**G**) and Xie (**H**). Mesenchymal cells were circled by dotted lines. Mesen, mesenchymal cell, Immu, immune cell, Epi, epithelial cell, Endo, endothelial cell.

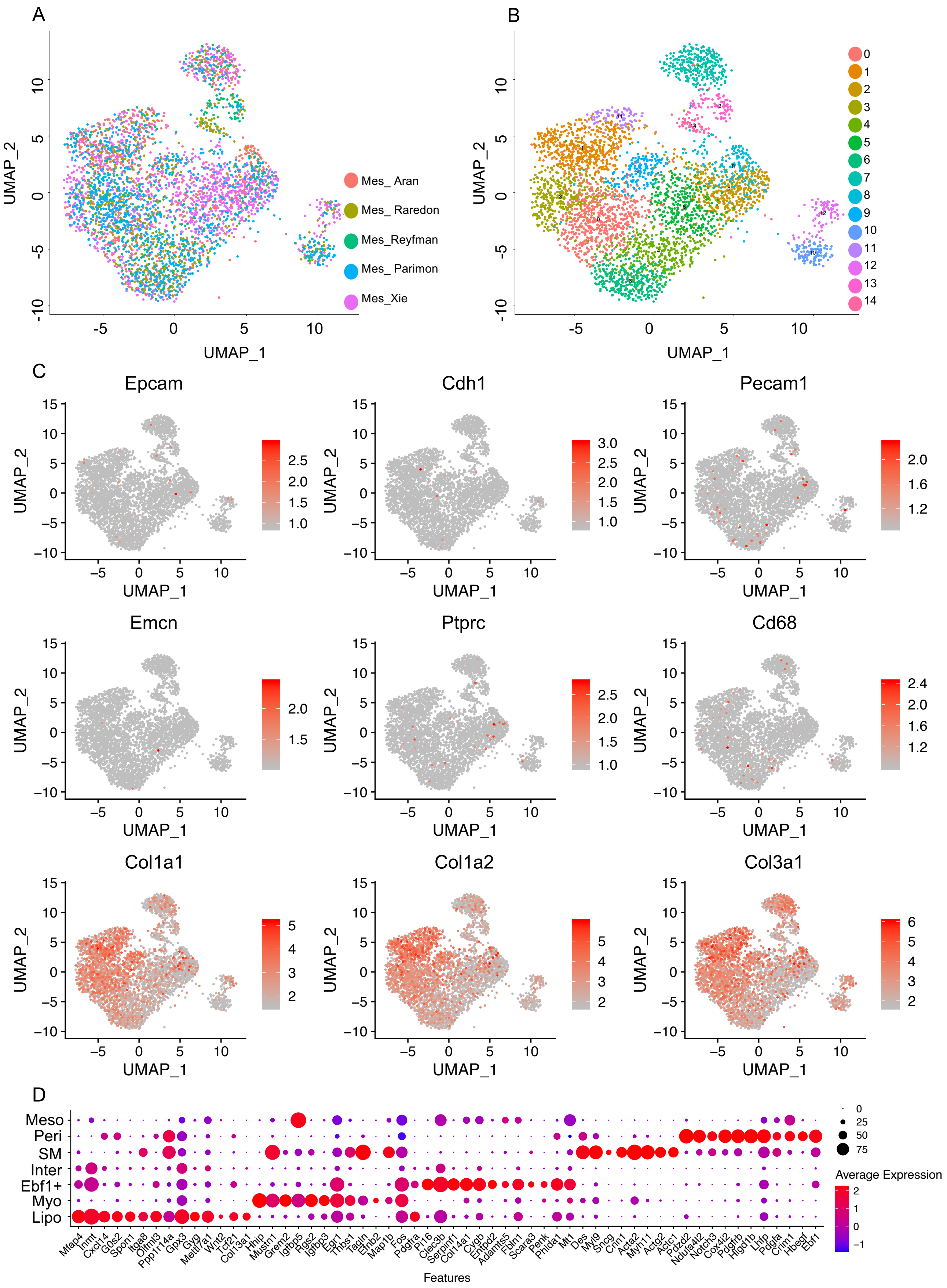


**Figure S14. Clustering of adult mouse lung fibroblasts.** UMAP visualization of mesenchymal cell integration (**A**) and clustering (**B**) of the fibroblasts from adult mouse lung. (**C**) UMAP visualization of relative expression of known cell type specific markers to validate the purity of fibroblasts. (**D**) Specific genes of each mesenchymal cell cluster visualized by dot plots.

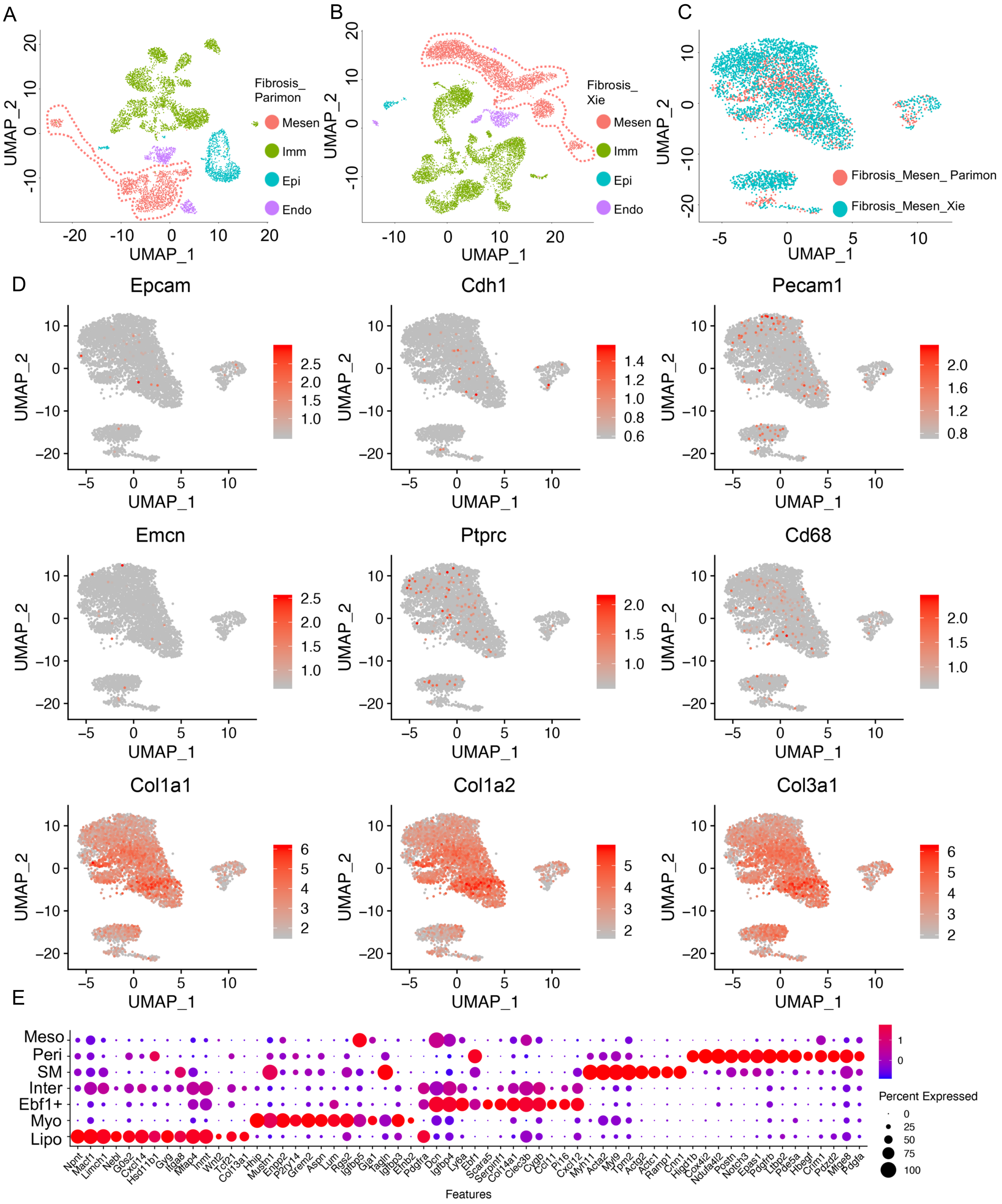


**Figure S15. Clustering of fibrosis mouse lung fibroblasts.** Cell types of fibrosis mouse lung from published papers were defined (**A**-**B**). Mesenchymal cells were circled by dotted lines. (**C**) UMAP visualization of mesenchymal cell integration and clustering of the fibroblasts from fibrosis mouse lung. (**D**) UMAP visualization of relative expression of known cell type specific markers to validate the purity of fibroblasts. (**E**) Specific genes of each mesenchymal cell cluster visualized by dot plots. Mesen, mesenchymal cells, Immu, immune cells, Epi, epithelial cells, Endo, endothelial cells.

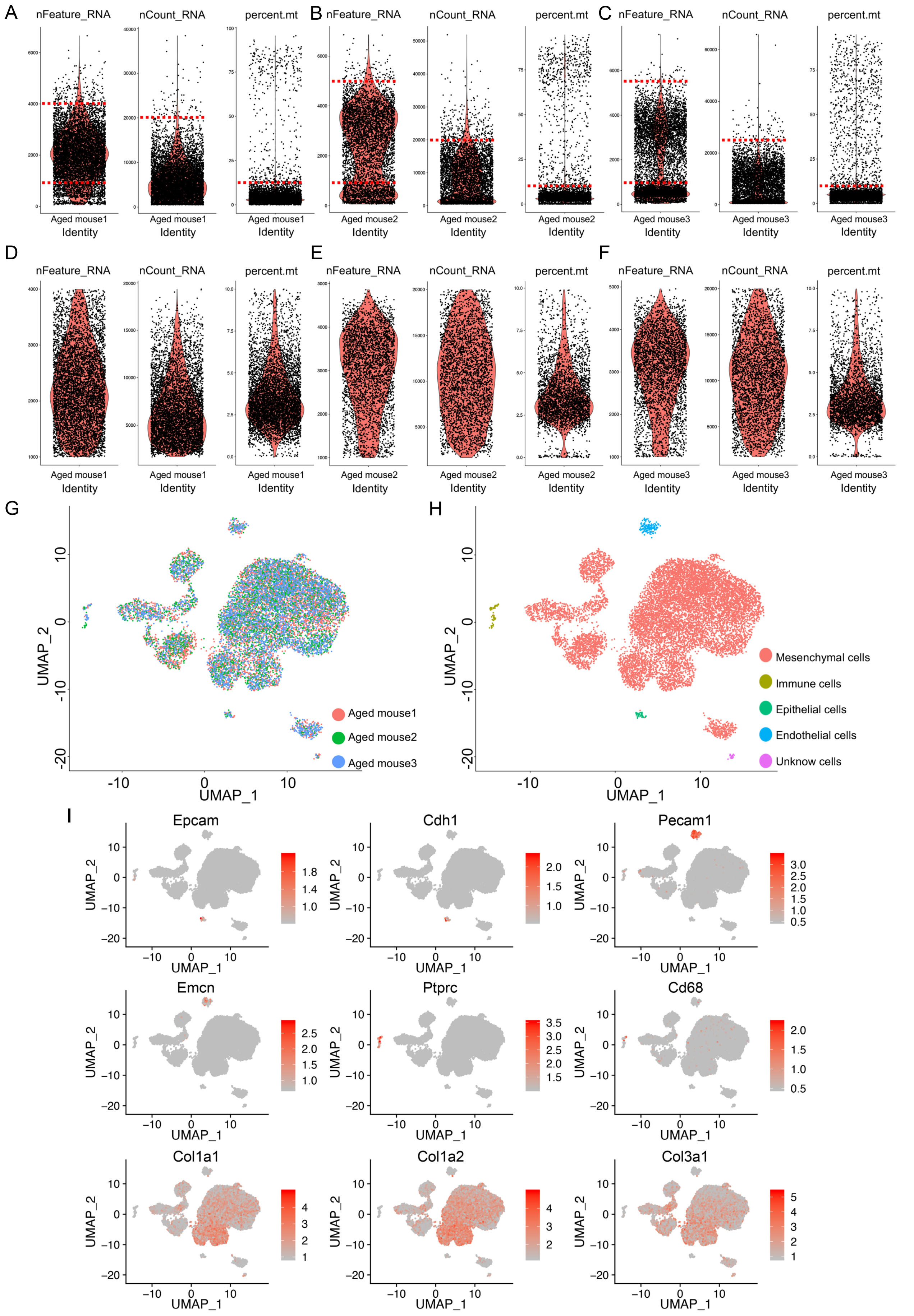


**Figure S16. Single cell RNA-seq on aged mouse lung.** Violin plot showing nFeature_RNA, nCount_RNA and percent.mt detected in each cell of each sample before (**A-C**) and after QC (**D-F**). UMAP visualization of mesenchymal cell integration (**G**) and cell types (**H**) of the single cells from aged mouse lung. (**I**) UMAP visualization of relative expression of known cell type specific markers.

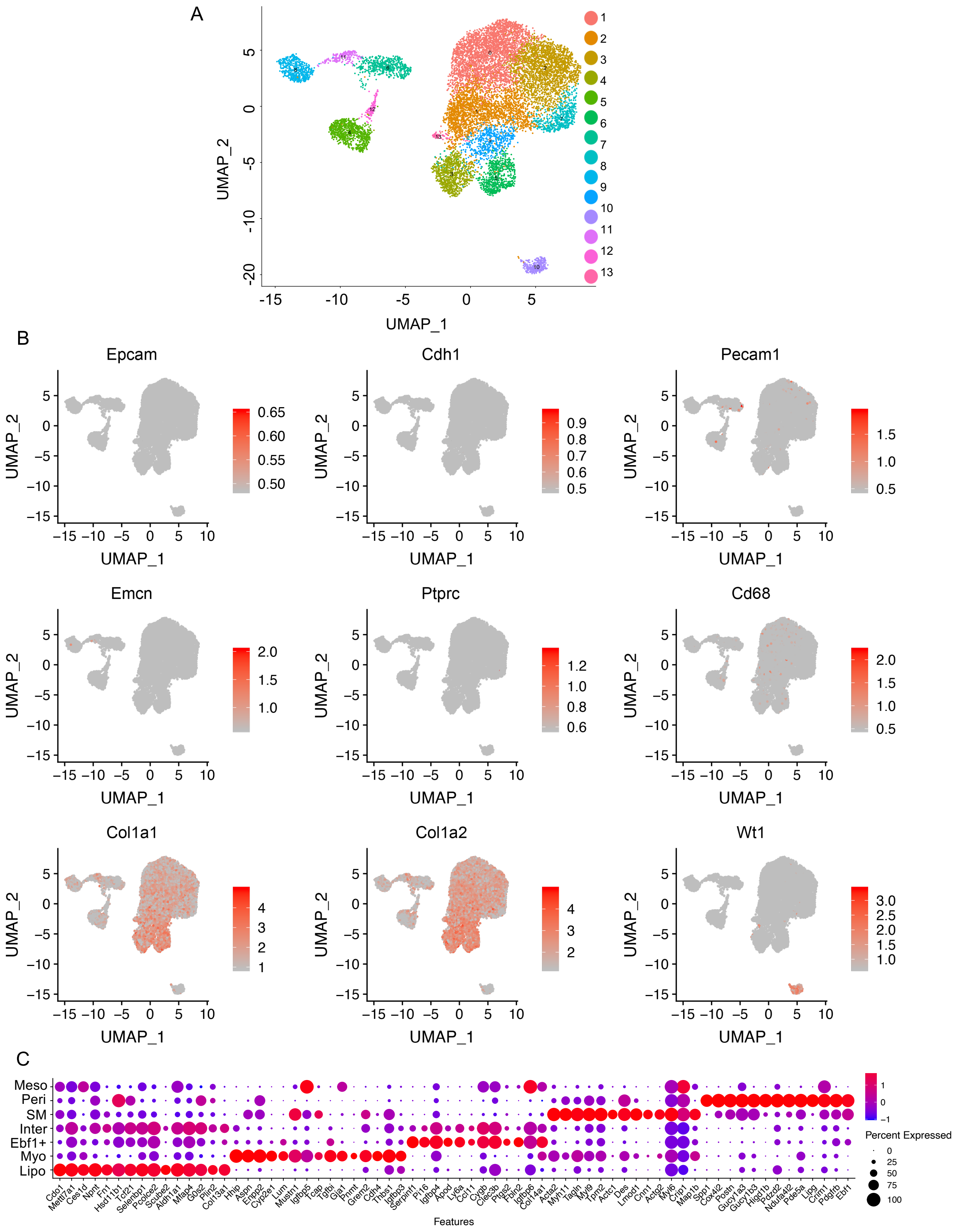


**Figure S17. Clustering of aged lung mesenchymal cells.** (**A**) UMAP visualization of mesenchymal cell clustering from aged mouse lung. (**B**) UMAP visualization of relative expression of known cell type specific markers to validate the purity of fibroblasts. (**C**) Specific genes of each mesenchymal cell cluster visualized by dot plot.

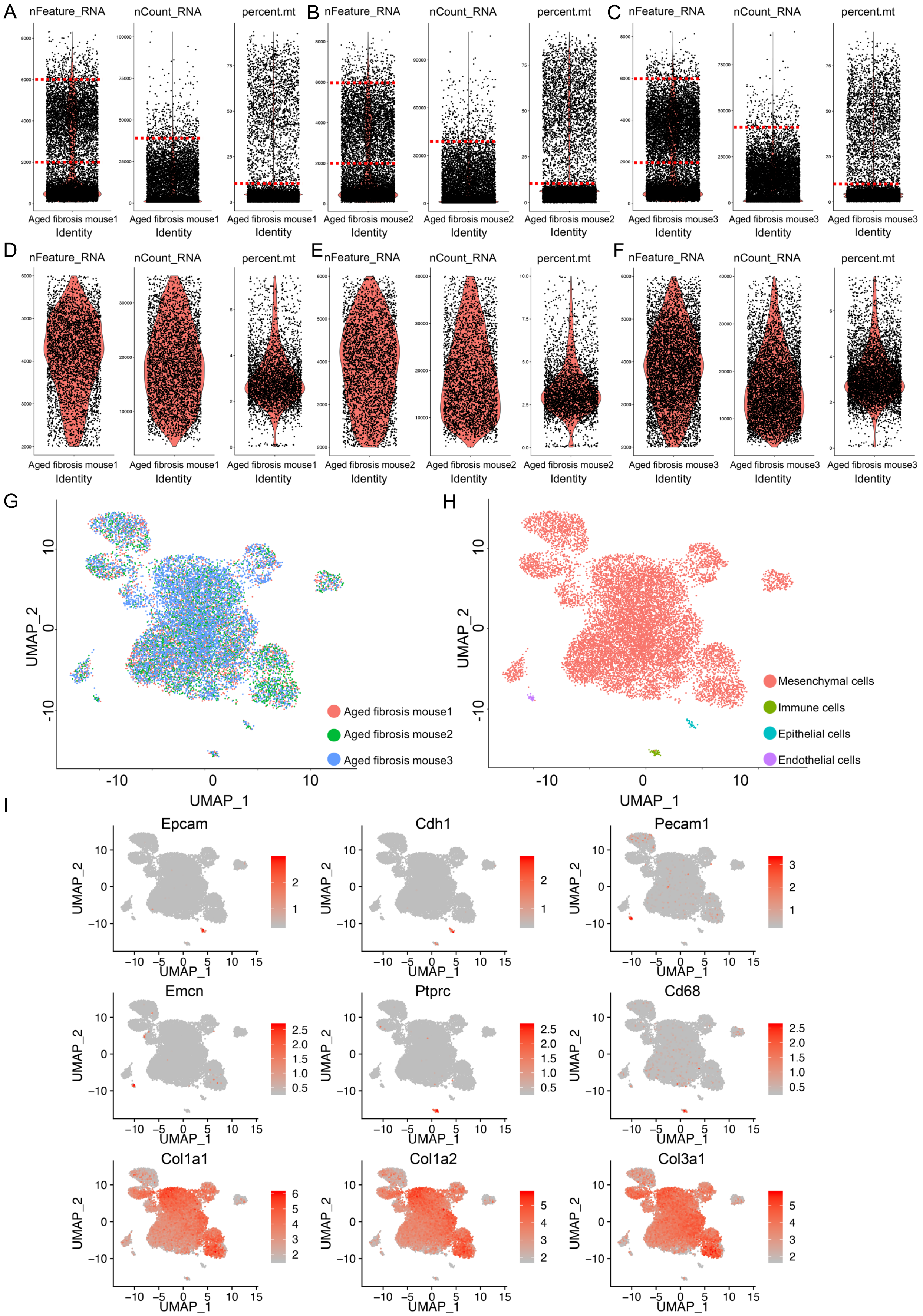


**Figure S18. Single cell RNA-seq on aged fibrosis mouse lung.** Violin plot showing nFeature_RNA, nCount_RNA and percent.mt detected in each cell of each sample before (**A-C**) and after QC (**D-F**). UMAP visualization of mesenchymal cell integration (**G**) and cell types (**H**) of the single cells from fibrosis mouse lung. (**I**) UMAP visualization of relative expression of known cell type specific markers.

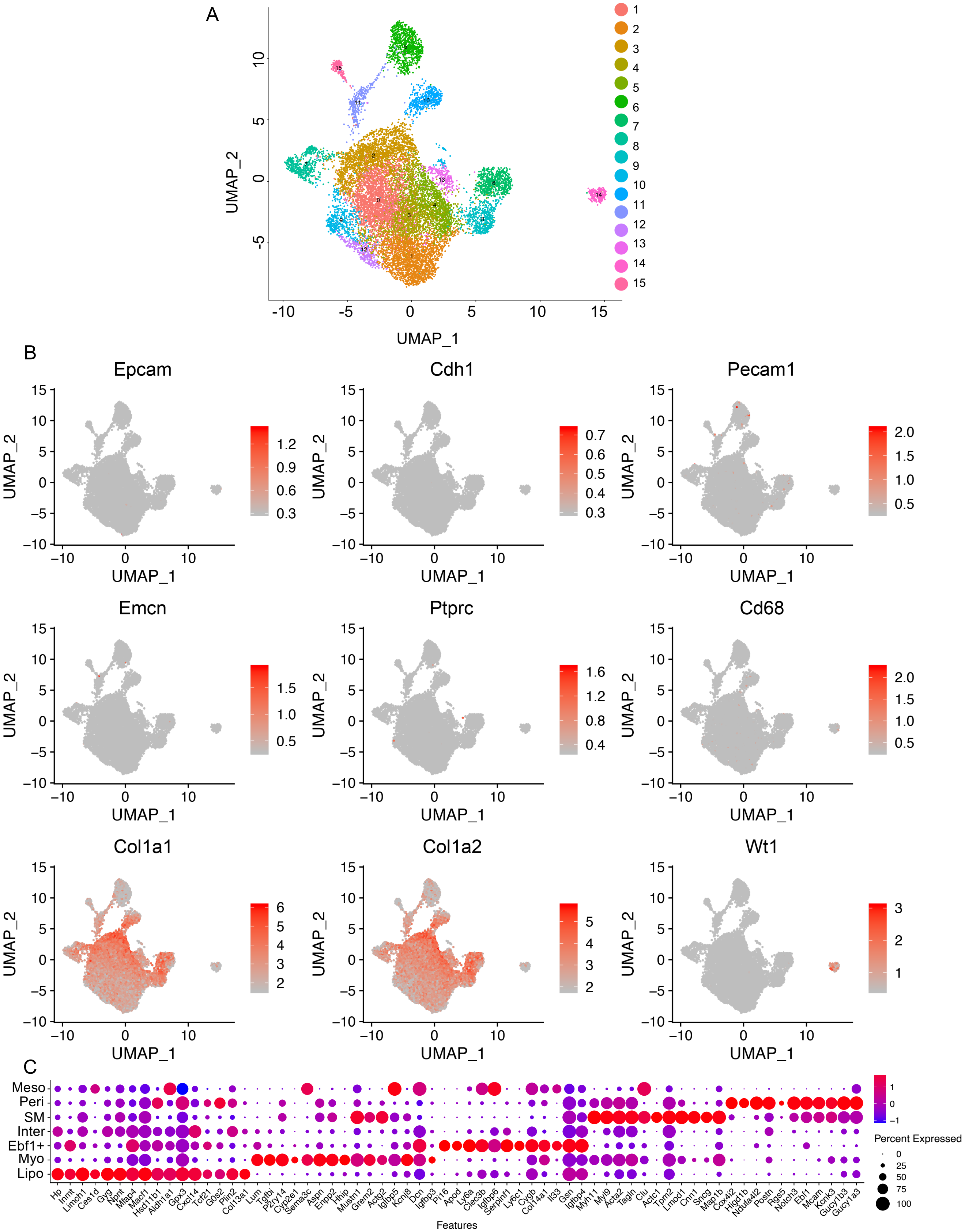

**Figure S19. Clustering of aged fibrosis lung mesenchymal cells.** (**A**) UMAP visualization of mesenchymal cell clustering from aged fibrosis mouse lung. (**B**) UMAP visualization of relative expression of known cell type specific markers to confirm the purity of fibroblasts. (**C**) Specific genes of each mesenchymal cell cluster visualized by dot plot.

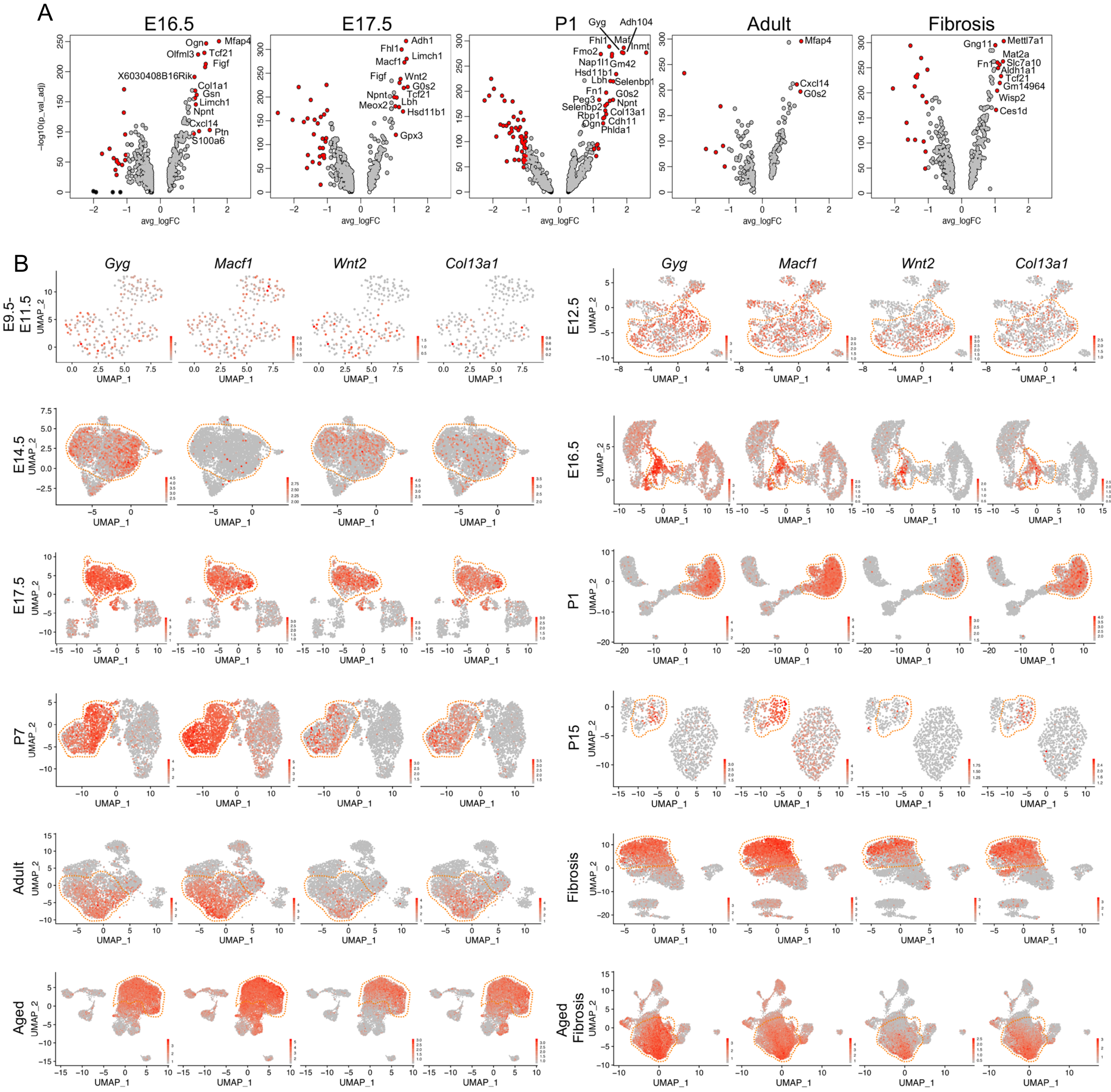


**Figure S20. Visualization of novel lipofibroblast marker expression in mouse lung lipofibroblasts of different timepoints.** (**A**) Visualization of differentially expressed genes of lipofibroblasts at each time point by Volcano plots. Genes highlighted in red have both a p-value < $10^{-5}$ and an average log fold-change (logFC) > 1, genes in black, p-value < $10^{-5}$, logFC < 1, genes in grey, p-value > $10^{-5}$, logFC > 1. (**B**) Visualization of Gyg, Macf1, Wnt2 and Col13a1 expression in mouse lung mesenchymal cells of different timepoints.

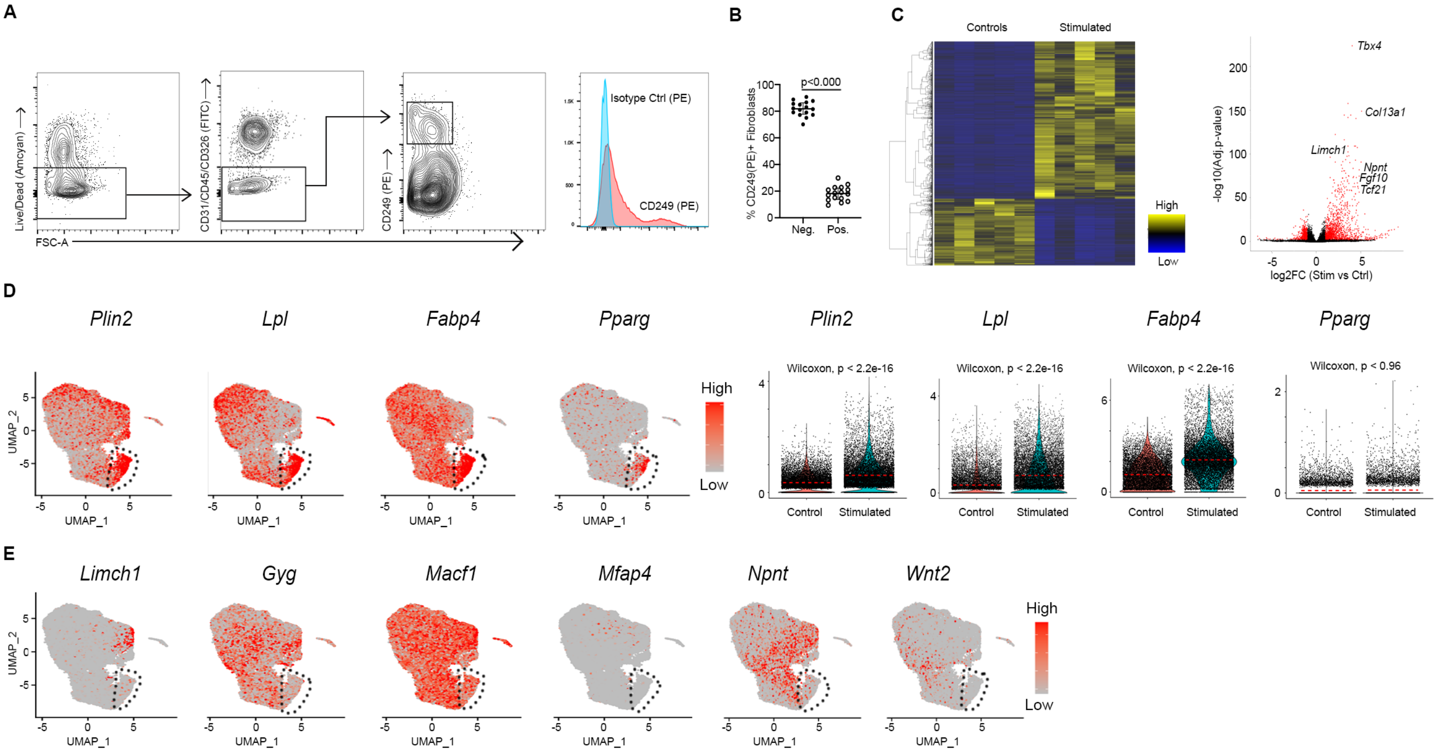


**Figure S21. Cd249⁺ fibroblast FACs and scRNA-seq of lipofibroblast-like cells.** (**A**) FACS gating strategy to obtain Cd249⁺ fibroblasts with concentration match isotype control overlay. (**B**) Mean (±SD) percentage Cd249⁺ fibroblasts vs Cd249⁻ fibroblasts obtained in each mouse lung. **(C**) Heatmap and volcano plot representation of the differentially expressed genes identified in bulk RNA-seq analysis of Cd249⁺ in comparison to Cd249⁻ fibroblasts. (**D**) UMAP visualization and violin plot representation of canonical lipofibroblast marker genes in cultured lipofibroblast-like cells and controls. Wilcoxon, $p < 2.2e-16$ per comparison. (**E**) UMAP visualization of the top differentially expressed genes identified in the scRNA-seq analysis of the in vivo lipofibroblast cluster in in vitro stimulated lipofibroblasts-like cells.

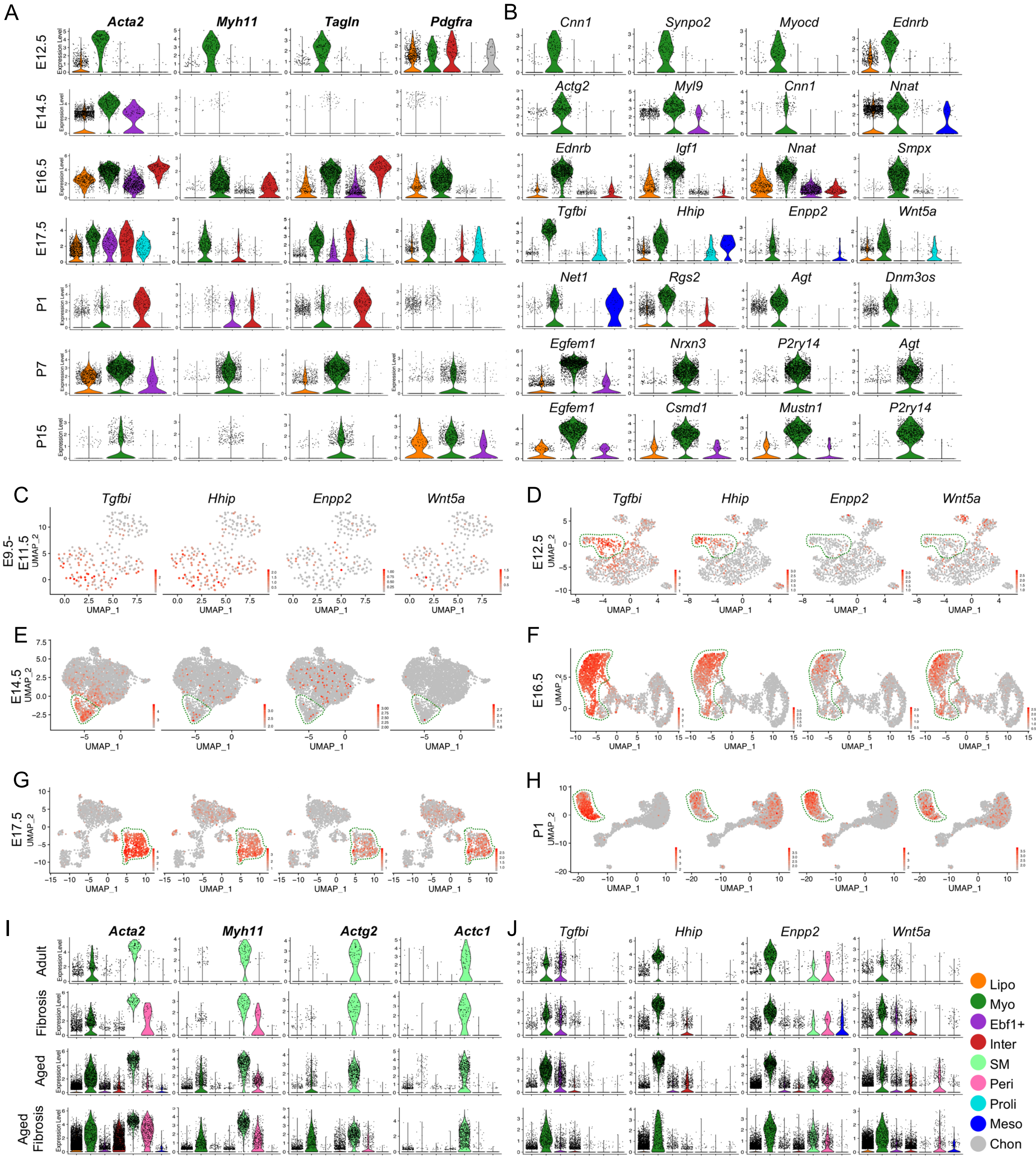


**Figure S22. Identification of myofibroblasts and SM cell markers.** Visualization of known (**A**) and timepoint specific genes for myofibroblasts (**B**) by violin plots in embryonic and postnatal mouse lung mesenchymal cells. UMAP visualization of novel myofibroblast markers in embryonic and postnatal mouse lung mesenchymal cells (**C**-**H**). Visualization of SMC specific markers (**I**) and novel myofibroblasts markers (**J**) in adult and aged normal and fibrosis mouse lung mesenchymal cells.

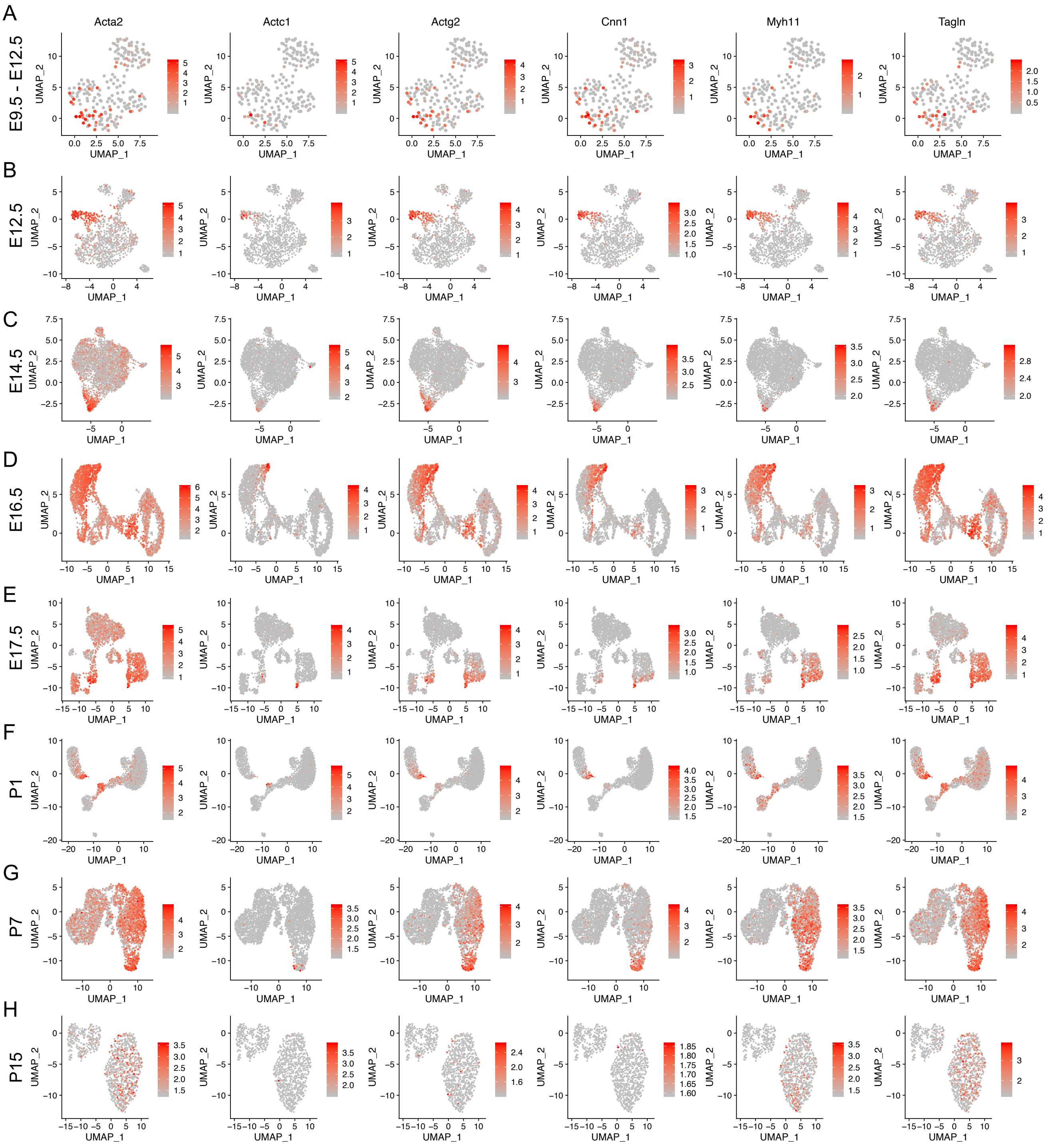


**Figure S23. SMC specific gene expression in embryonic and postnatal lung mesenchymal cells.** SMC specific gene expression in E9.5-E11.5 (**A**), E12.5 (**B**), E14.5 (**C**), E16.5 (**D**), E17.5 (**E**), P1 (**F**), P7 (**G**) and P15 (**H**) mouse lung mesenchymal cells. Broad or rare expression of these genes indicated no distinct SMC clusters in these mesenchymal cell datasets of these timepoints

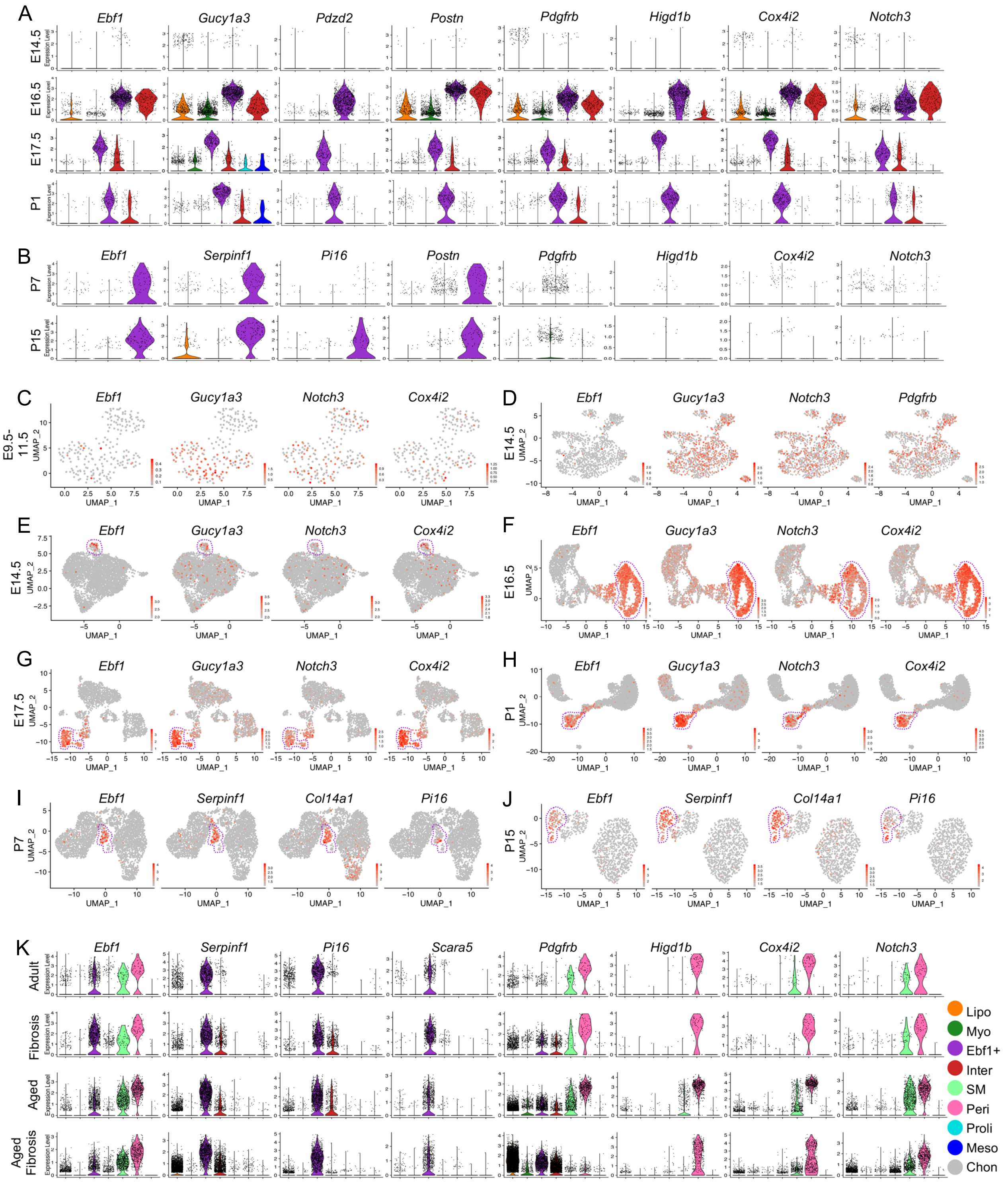


**Figure S24. Identification of *Ebf1*⁺ and pericyte subtypes.** (**A**-**B**) Visualization of timepoint specific genes for *Ebf1*⁺ fibroblasts by violin plots in embryonic and postnatal mouse lung mesenchymal cells. (**C**-**J**) UMAP visualization of *Ebf1*+ fibroblast specific markers in embryonic and postnatal mouse lung mesenchymal cells. (**K**) violin plot visualization of Ebf1⁺ fibroblast and pericyte specific genes in adult and aged normal and fibrosis mouse lung mesenchymal cells.

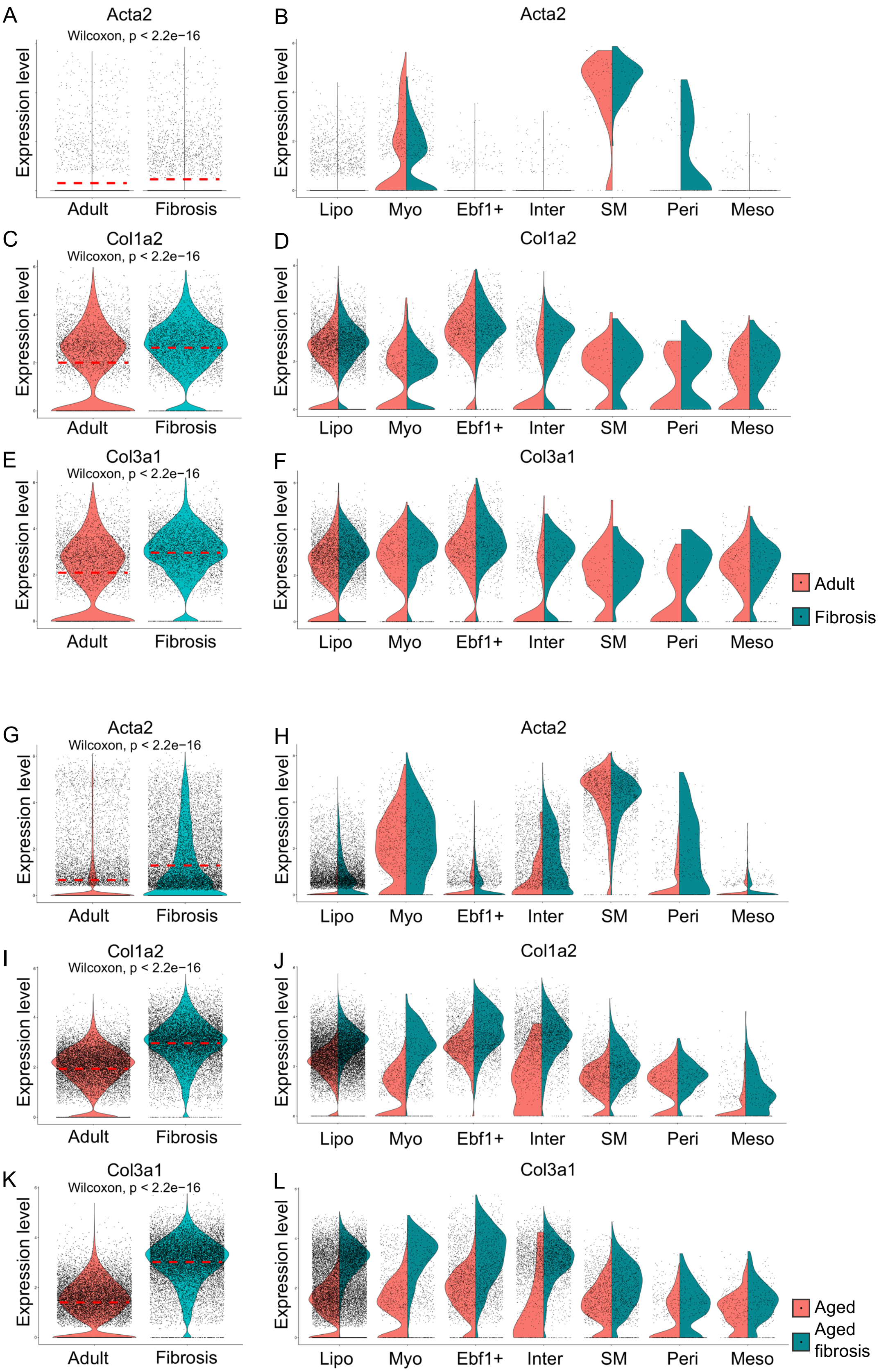


**Figure S25. ECM related genes expression in normal and fibrosis lung total mesenchymal cells and mesenchymal cell subtypes.** Acta2 (**A**, **B**, **G** and **H**), Col1a2 (**C**, **D**, **I** and **J**), and Col3a1 (**E**, **F**, **K** and **L**) transcripts in total normal and fibrosis lung mesenchymal cell (**A**, **C**, **E**, **G**, **I** and **K**) and mesenchymal cell subtypes (**B**, **D**, **F**, **H**, **J** and **L**) of adult (**A**-**F**) and aged (**G**-**L**) mouse were visualized by violin plots.

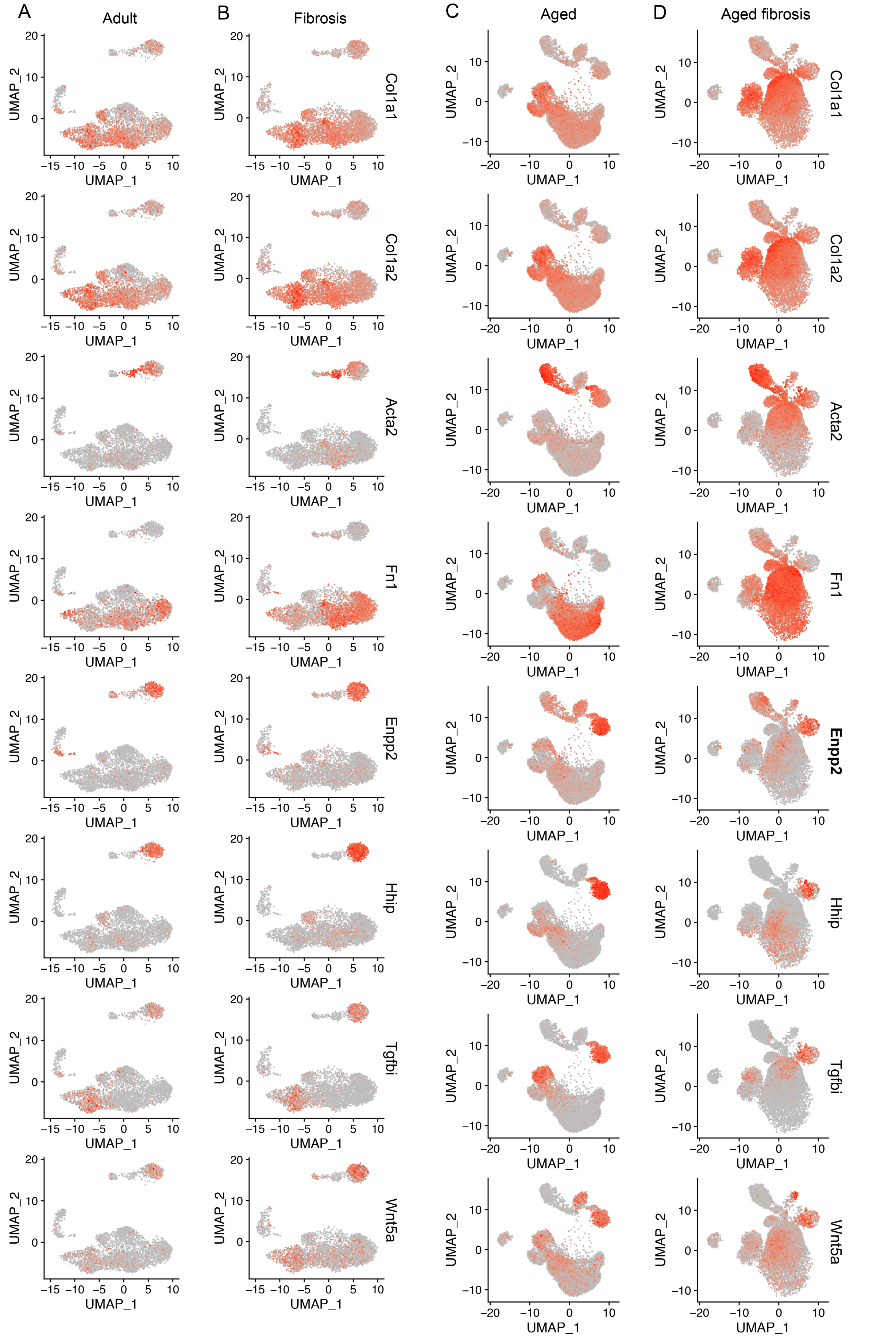


**Figure S26. Expression of fibrotic related gene and novel myofibroblast genes was visualized by UMAP.** Transcript of known fibrotic related genes, *Col1a1*, *Col1a2*, *Acta2*, *Fn1* and myofibroblast specific genes, *Enpp2*, *Hhip*, *Tgfbi*, *Wnt5a* in adult (**A**), fibrosis (**B**), aged (**C**) and aged fibrosis (**D**) mesenchymal cells.

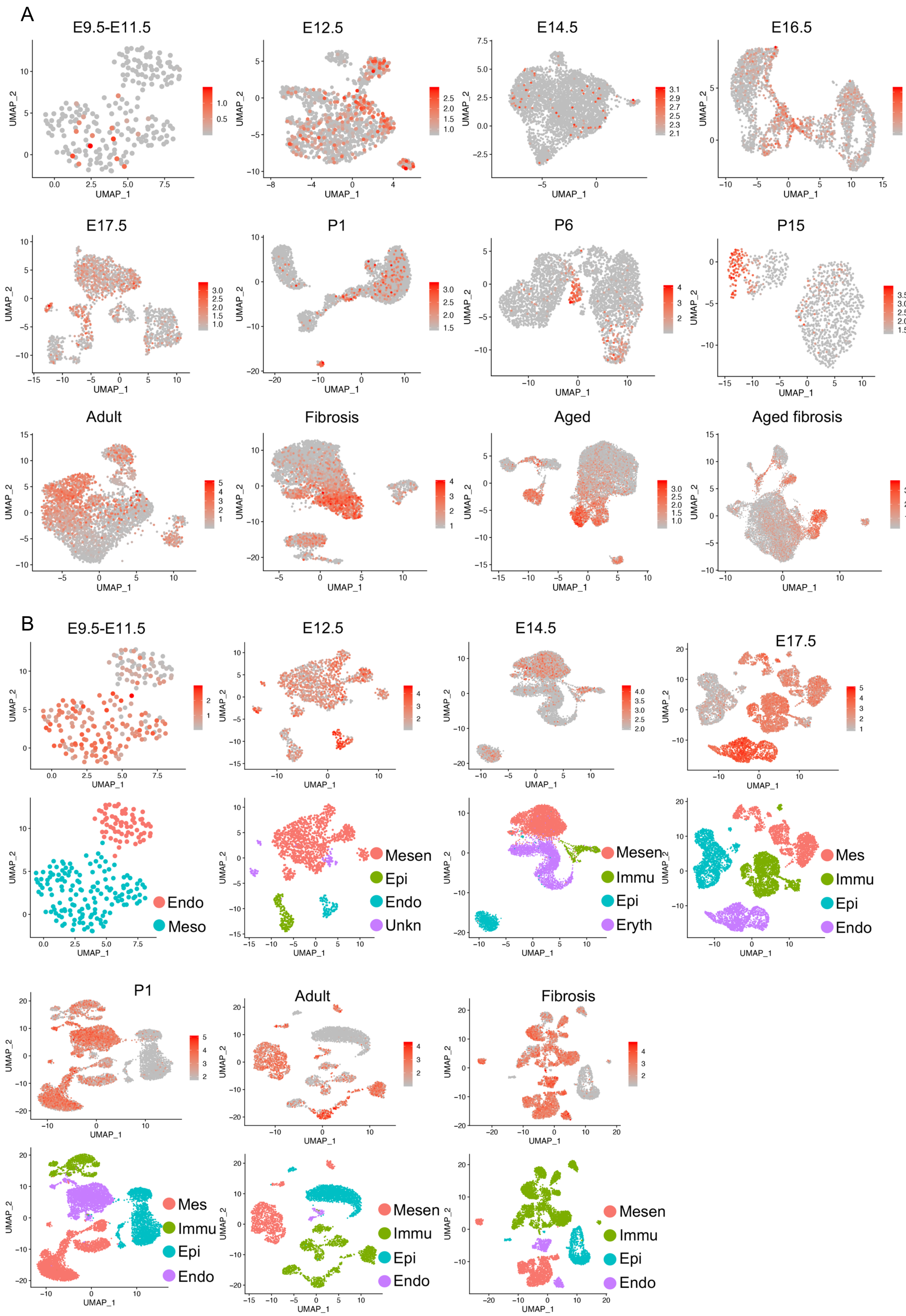


**Figure S27. Transcript of *Col14a1* in lung mesenchymal cells and *Vim in lung total cells* of different timepoints**. (**A**) In embryonic stages, *Col14a1* was mainly expressed in pre-lipofibroblasts/lipofibroblasts, however, after birth, *Col14a1* expression switched to Ebf+ fibroblasts and in adult and aged lungs *Col14a1* kept its transcripts in Ebf1+ fibroblasts. (**B**) In early embryonic stages (E9.5-E11.5), *Vim* showed transcripts in both mesoderm and endoderm cells. In later embryonic, postnatal, adult and fibrosis stages, *Vim* showed highest transcripts in endothelial cells, showed weakened transcripts in immune cells and mesenchymal cells and was rarely detectable in epithelial cells. Endo, endoderm, Meso, mesoderm, Mesen, mesenchymal cells, Immu, immune cells, Epi, epithelial cells, Endo, endothelial cells, Unkn, unknown.

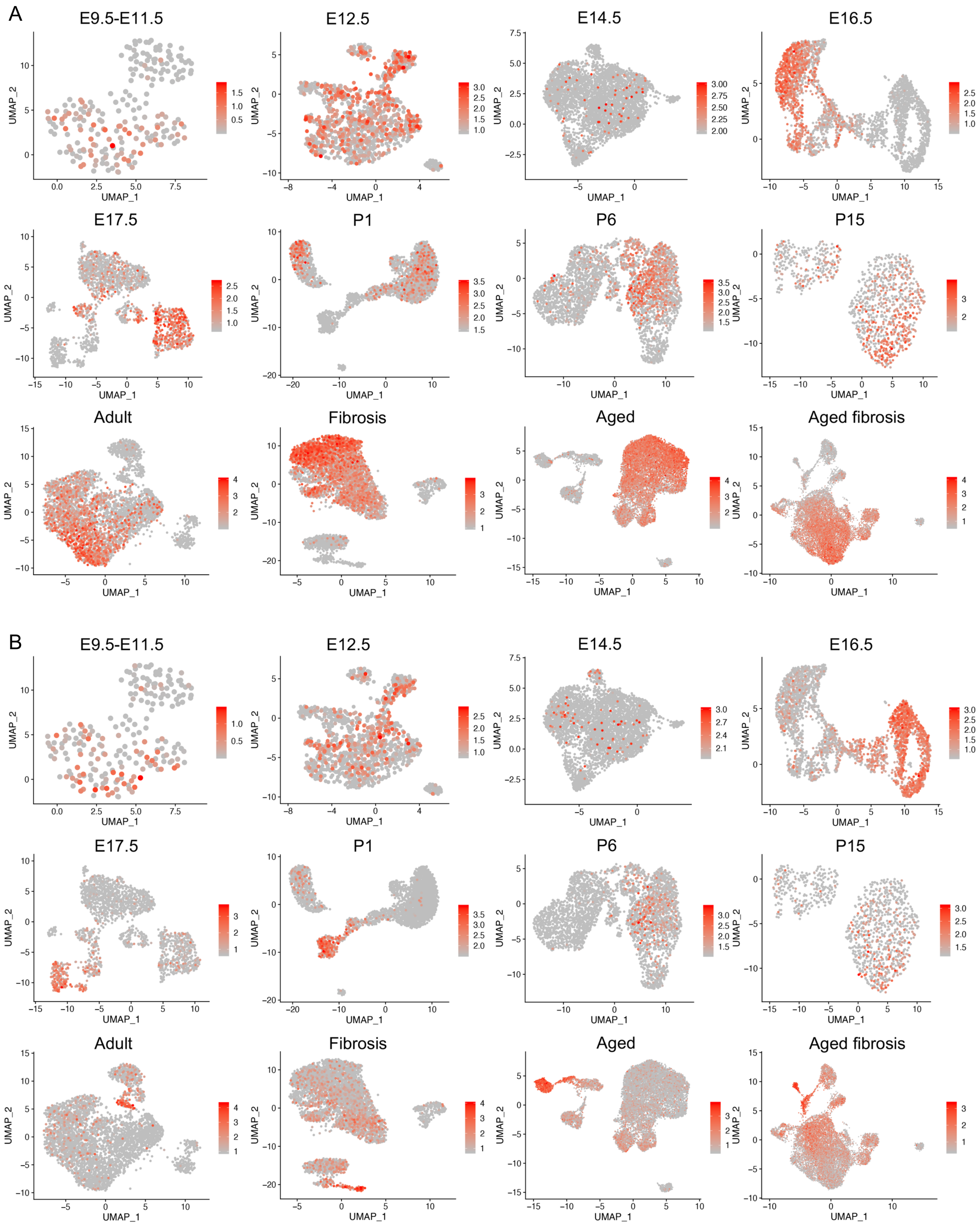


**Figure S28. Expression of *Pdgfra* and *Pdgfrb* in mouse lung mesenchymal cells of different timepoints.** (**A**) *Pdgfra* transcripts in mesenchymal cells of different time points. *Pdgfra* expression was mainly in myofibroblasts in embryonic and postnatal lung mesenchymal cells and switched to lipofibroblasts in adult and aged normal and fibrosis mesenchymal cells with high background in other subtypes. (**B**) *Pdgfrb* showed higher expression in Ebf1+ fibroblasts and pericytes with some background in other subtypes.

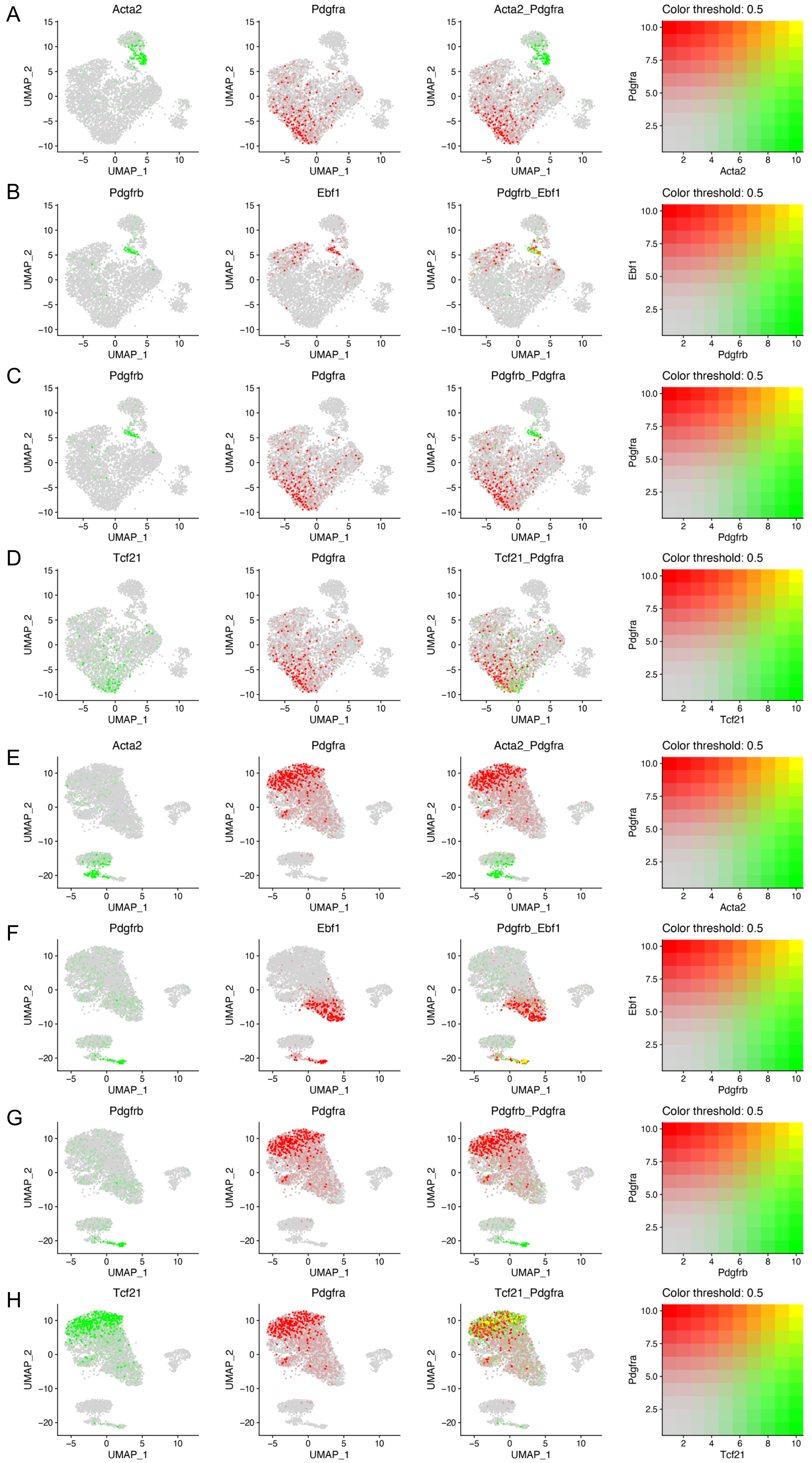


**Figure S29. Co-expression of genes in normal and fibrosis lung mesenchymal cells**. Visualization of blend expression of *Acta2* and *Pdgfra* (**A** and **E**), *Pdgfrb* and *Ebf1* (**B** and **F**), *Pdgfrb* and *Pdgfra* (**C** and **G**), *Tcf21* and *Pdgfra* (**D** and **H**) in normal (**A** - **E**) and fibrosis (**E** - **H**) mouse lung mesenchymal cells.

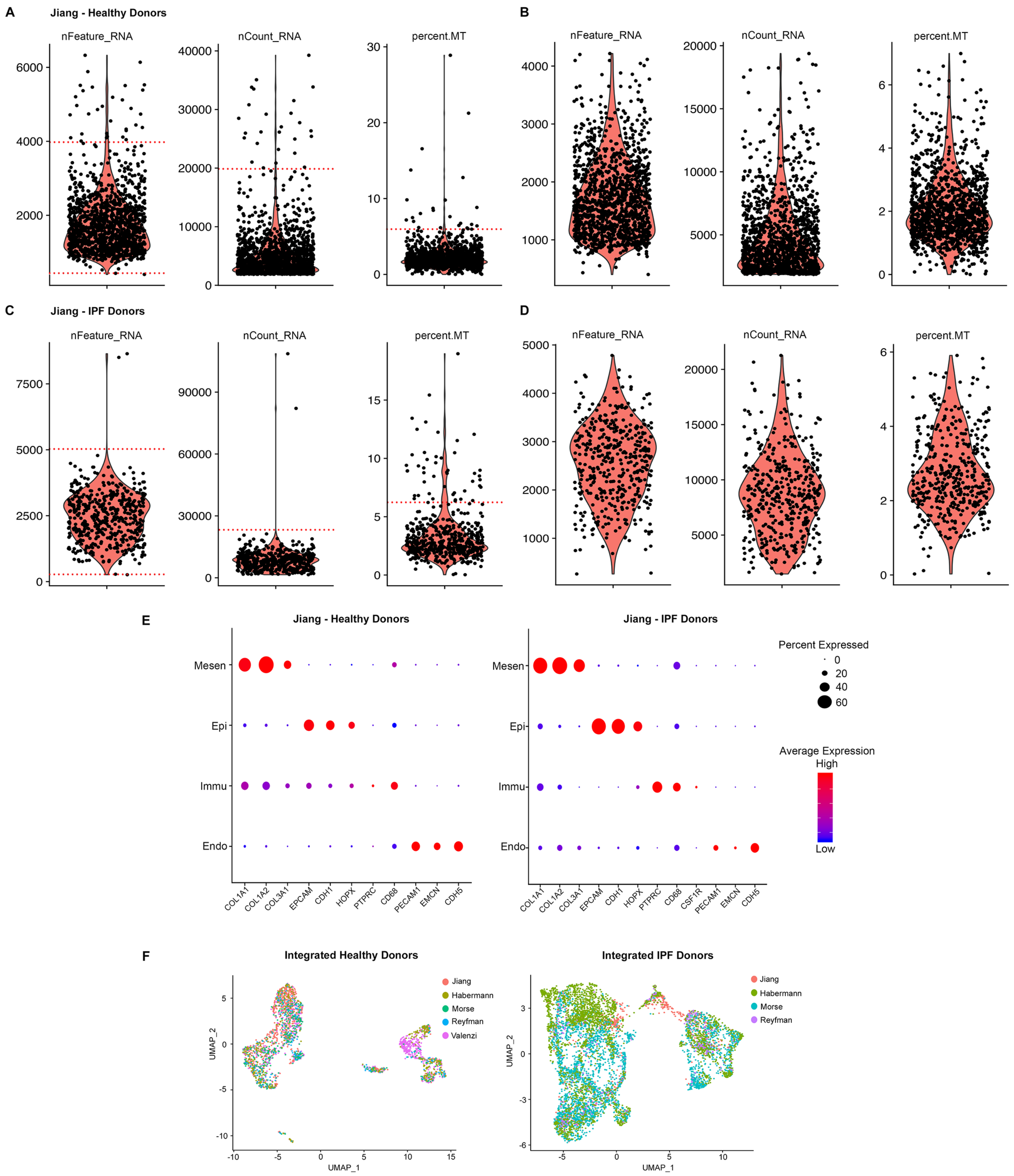


**Figure S30. Representative quality control, sub-setting and integration of human scRNA-seq data.** Violin plot showing number of genes (nFeature_RNA), number of read counts (nCount_RNA) and the percentage of transcripts mapping to mitochondrial genes (percent.MT) detected in each cell before **(A, C)** and after **(B, D)** QC in healthy **(A, B)** and IPF **(C, D)** donor samples from Jiang lab. **(E)** Dot plot representation of common marker genes used to identify and subset mesenchymal from epithelial, endothelial and immune cells in healthy **(E)** and IPF **(F)** donor samples . Dot size corresponds to percentage of cells in a cluster expressing the gene and color to expression level. UMAP visualization to the integration of scRNA-seq data sets of healthy **(G)** and IPF **(H)** donor lung mesenchymal cells. Mesen, Mesenchymal, Epi, Epithelial, Immu, Immune, Endo, Endothelial.

Supplementary Table 1 Mouse Single Cell Sample Details

| Data origin | | Cell number | | Cell number/cell type | | | | | |
|---|---|---|---|---|---|---|---|---|---|
| | n | Before QC | After QC | Mesoderm | | | Endoderm | | Sequencing platform |
| E9.5-E11.5 (Pijuan-Sala) | 4 | 224 | 215 | 149 | | | 66 | | Illumina HiSeq 2500 |
| Data origin | | Before QC | After QC | Mesen | Imm | Endo | Epi | Eryth/Ukn | Sequencing platform |
| E12.5 (Cohen) | 4 | 1,498 | 1,454 | 1,158 | | 80 | 145 | 71 | Illumina NextSeq 500 |
| E14.5 (Han) | 1 | 8,395 | 8,351 | 4,246 | 470 | | 921 | 2,714 | Illumina Hiseq system |
| E16.5 | 1 | 3,085 | 3,013 | 2,981 | | 14 | 18 | | Illumina NextSeq 500 |
| E17.5 | 3 | 7,479 | 7,477 | 2,002 | 2,193 | 1,383 | 1,899 | | Illumina NextSeq 500 |
| P1 (Guo) | 2 | 9,944 | 9,722 | 3,664 | 976 | 2,645 | 2,437 | | Drop-seq |
| P7 (Li) | 1 | 3,213 | 3,125 | 3,125 | | | | | HiSeq4000 |
| P15 (Li) | 1 | 1,233 | 1,202 | 1,097 | 105 | | | | HiSeq4000 |
| Adult (Aran) | 7 | 25,900 | 25,216 | 606 | | | 24,610 | | Drop-seq |
| Adult (Xie) | 1 | 5,481 | 5,306 | 1,547 | 3,371 | 305 | 83 | | Illumina NextSeq 500 |
| Adult (Raredon) | 2 | 7,952 | 7,397 | 621 | | | 6,776 | | Illumina HiSeq 4000 |
| Adult (Reyfman) | 2 | 14,518 | 14,169 | 356 | 9,966 | 1,290 | 2,537 | | Illumina HiSeq 4000 |
| Adult (Parimon) | 1 | 4,523 | 4,448 | 1,063 | 1,450 | 81 | 1,854 | | Illumina Novaseq 6000 |
| Fibrosis (Xie) | 1 | 9,813 | 9,756 | 3,399 | 5,729 | 514 | 114 | | Illumina NextSeq 500 |
| Fibrosis (Parimon) | 1 | 6,465 | 5,741 | 1,329 | 3,019 | 504 | 889 | | Illumina Novaseq 6000 |
| Aged | 3 | 20,452 | 12,563 | 12,304 | 56 | 137 | 40 | 26 | Illumina NextSeq 500 |
| Aged fibrosis | 3 | 40,243 | 13,425 | 13,335 | 35 | 23 | 32 | | Illumina NextSeq 500 |

Pijuan-Sala, Blanca Pijuan-Sala publication; Cohen, Merav Cohen publication; Han, Xiaoping Han publication; Guo, Minzhe Guo publication; Li, Rongbo Li publication; Aran, Dvir Aran publication; Xie, Ting Xie publication, Raredon, Micha Sam Brickman Raredon publication; Reyfman, Paul A. Reyfman publication; Parimon, Tanyalak Parimon publication; n, sample number per condition; QC, quality control, Mesen, mesenchymal cells, Immu, immune cells, Endo, endothelial cells, Eryth, erythroid cells, Ukn, uknown cell type.

Supplementary Table 2 Human scRNA-seq sample details

| Data origin | Pub (Y/N) | Status | Smoker (Y:N) | M:F | N | Cell number Pre QC | Post QC | Cell type (number, post QC) Immu | Epi | Endo | Mesen | Sequencing platform |
|---|---|---|---|---|---|---|---|---|---|---|---|---|
| Jiang | N | Healthy | - | - | 5 | 2,282 | 2,171 | 300 | 854 | 161 | 850 | Illumina NextSeq 500 |
| | N | IPF | - | - | 5 | 1,412 | 1,266 | 14 | 590 | 38 | 624 | |
| Val | Y | Healthy | - | 3:2 | 5 | 26,036 | 24,760 | 20,128 | 2,132 | 1,638 | 862 | Illumina NextSeq 500 |
| Mor | Y | Healthy | 0:6 | 4:2 | 6 | 29,687 | 27,151 | 21,941 | 2,346 | 2,300 | 564 | Illumina NextSeq 500 |
| | Y | IPF | 1:2 | 1:2 | 3 | 37,935 | 33,808 | 18,789 | 6,290 | 4,656 | 4,073 | |
| Haber | Y | Healthy | 8:2 | 7:3 | 10 | 31,644 | 31,281 | 19,493 | 8,139 | 3,156 | 493 | Illumina Novaseq 6000/HiSeq 4000 |
| | Y | IPF | 11:9 | 13:7 | 20 | 57,682 | 57,522 | 25,696 | 24,891 | 4,080 | 2,855 | |
| Reyf | Y | Healthy | 3:5 | 2:6 | 8 | 43,632 | 41,175 | 16,814 | 23,312 | 870 | 179 | Illumina HiSeq 4000 |
| | Y | IPF | 3:2 | 3:2 | 5 | 15,650 | 14,790 | 10,292 | 3,726 | 527 | 245 | |

Jiang, Jiang/Noble Cohort Cedars-Sinai Medical Center; Val, Valenzi publication; Mor, Morse publication lung; Haber, Haberman publication; Reyf, Reyfmann publication; Pub, Published, Y: Yes; N, No; Healthy, Healthy (non-fibrotic) donor; IPF, Idiopathic pulmonary fibrosis patient; M, Male; F, Female; N, sample number per group; QC, Quality control; Endo, Endothelial cells; Epi, Epithelial cells; Immu, Immune cells; Mesen, Mesenchymal cells.

Supplementary Table 4 antibodies for Flow cytometry, FACs and immunofluorescence

Antibodies for Flow cytometry or FACs

| Antibody | Conjugate | Clone | Manufacturer | Catalogue Number |
|---|---|---|---|---|
| Cd24 | Biotin | M1/69 | Biolegend | 101804 |
| Cd31 | Biotin | 390 | Biolegend | 102404 |
| Cd31 | FITC | Mec13.3 | BD | 553080 |
| Cd34 | Biotin | MEC14.7 | Biolegend | 119304 |
| Cd45 | Biotin | 30-F11 | Biolegend | 103104 |
| Cd45.2 | FITC | 104 | BD | 553372 |
| Cd326 | FITC | G8.8 | Invitrogen | 11-5791-82 |
| Cd326 | PE/Cy7 | G8.8 | Biolegend | 118216 |
| Cd249 | PE | BP-1 | BD | 553735 |
| Sca-1 | PE | D7 | BD | 553108 |
| IgG2a, κ Isotype Control | PE | G155-178 | BD | 553457 |
| Strepavidin | PE | - | eBiosciences | 12-4317-87 |
| CD326 | Alexa Fluor® 647 | 9C4 | Biolegend | 324212 |
| CD31 | PE/Cy7 | VM59 | Biolegend | 303118 |
| CD45 | PE/Cy7 | HI30 | Biolegend | 304016 |
| Fixable viability | eFluor 780 | - | Invitrogen | 65-0865-14 |
| Fixable viability | eFluor 506 | - | Invitrogen | 65-0866-14 |

Antibodies for immunofluorescence

| Antibody | Conjugate | Clone | Manufacturer | Catalogue Number |
|---|---|---|---|---|
| EBF-1 | - | - | R&D system | AF5165 |
| VWF | - | - | Abcam | ab6994 |
| αSMA | - | 1A4 | Santa Cruz Biotechnology | sc-32251 |
| Donkey Anti-Goat IgG (H+L) | Alexa Fluor® 488 | - | Jackson Immuno Research | 705-545-003 |
| Donkey anti-Rabbit IgG (H+L) | Alexa Fluor® 647 | - | Thermo Fisher Scientific | A-31573 |
| Donkey anti-Mouse IgG (H+L) | Alexa Fluor® 555 | - | Thermo Fisher Scientific | A-31570 |
